# Temporal Dynamics of Microbial Communities in Anaerobic Digestion: Influence of Temperature and Feedstock Composition on Reactor Performance and Stability

**DOI:** 10.1101/2025.01.12.632589

**Authors:** Ellen Piercy, Xinyang Sun, Peter R. Ellis, Mark Taylor, Miao Guo

## Abstract

Anaerobic digestion (AD) offers a sustainable biotechnology to recover resources from carbon- and nutrient-rich wastewater streams, such as food-processing wastewater. Despite the wide adoption of crude wastewater characterisation, the impact of detailed chemical fingerprinting on AD remains underexplored. This study investigated the influence of fermentation-wastewater composition and operational parameters on AD over time to identify critical parameters influencing microbiome diversity and reactor performance. Eighteen bioreactors were operated under various operational conditions using mycoprotein fermentation wastewater. Detailed chemical analysis fingerprinted the molecules in the fermentation-wastewater throughout the AD process including sugars, sugar alcohols and volatile fatty acids (VFAs). High-throughput sequencing revealed distinct microbiome profiles linked to temperature and reactor configuration, with mesophilic conditions supporting a more diverse and densely connected microbiome. Importantly, significant elevations in *Methanomassiliicoccus* were correlated to high butyric acid concentrations and decreased biogas production, further elucidating the role of this newly discovered methanogen in AD. Reactors from different experimental runs had distinct VFA profiles, which impacted microbial taxonomy and diversity. Dissimilarity analysis demonstrated the importance of individual VFAs, sugars and sugar alcohols on microbiome diversity, highlighting the need for detailed chemical fingerprinting in AD studies of microbial trends. Furthermore, machine learning models predicting reactor performance achieved high accuracy based on operational parameters and microbial taxonomy. Operational parameters were found to have the most substantial influence on chemical oxygen demand removal, whilst *Oscillibacter* and two *Clostridium* species were highlighted as key factors in biogas production. By integrating detailed chemical and biological fingerprinting with explainable machine learning models this research presents a novel approach to advance our understanding of AD microbial ecology, offering insights for industrial applications of sustainable waste-to-energy systems.

## 1. Introduction

Global wastewater is estimated to have reached 400 billion m^3^/year ^1^. Alarmingly, 48% of wastewater is inadequately treated, posing significant environmental and public health risks^2^. Compared with municipal wastewater, food-processing wastewater is often characterised by high organic matter concentrations, which require advanced treatment technologies ^3^. Aerobic wastewater treatments are energy-intensive, cost ineffective, and contribute to greenhouse gas emissions ^4^. This realisation has prompted a shift towards more sustainable approaches, which recover valuable compounds from wastewater. Anaerobic digestion (AD) offers a biological process that converts organic matter to energy-carrier gas (biogas and nutrient-rich digestate), which has the potential to reduce environmental impacts and operational costs ^5,6^.

Many studies have focused on lab-scale reactors using synthetic feedstock, such as acetic acid or glucose-based growth mediums (Figure 1a). However, synthetic feedstock is not representative of the fluctuations and contaminants within real-world wastewater. Detailed chemical composition of wastewater feedstock remains largely undefined, especially within studies including detailed biological characterisation. Diverse microbes show different substrate affinities; therefore, it is important to understand how the molecules present in wastewater streams impact microbial diversity (Figure 1b). Moreover, previous studies often focus on singular time points, which do not capture the dynamic nature of the microbiome in response to fluctuating feedstock structure and composition. Understanding the interactions between these parameters and microbial communities over temporal scales is essential for enhancing biogas production, process stability, and overall efficiency.

**Figure 1.**
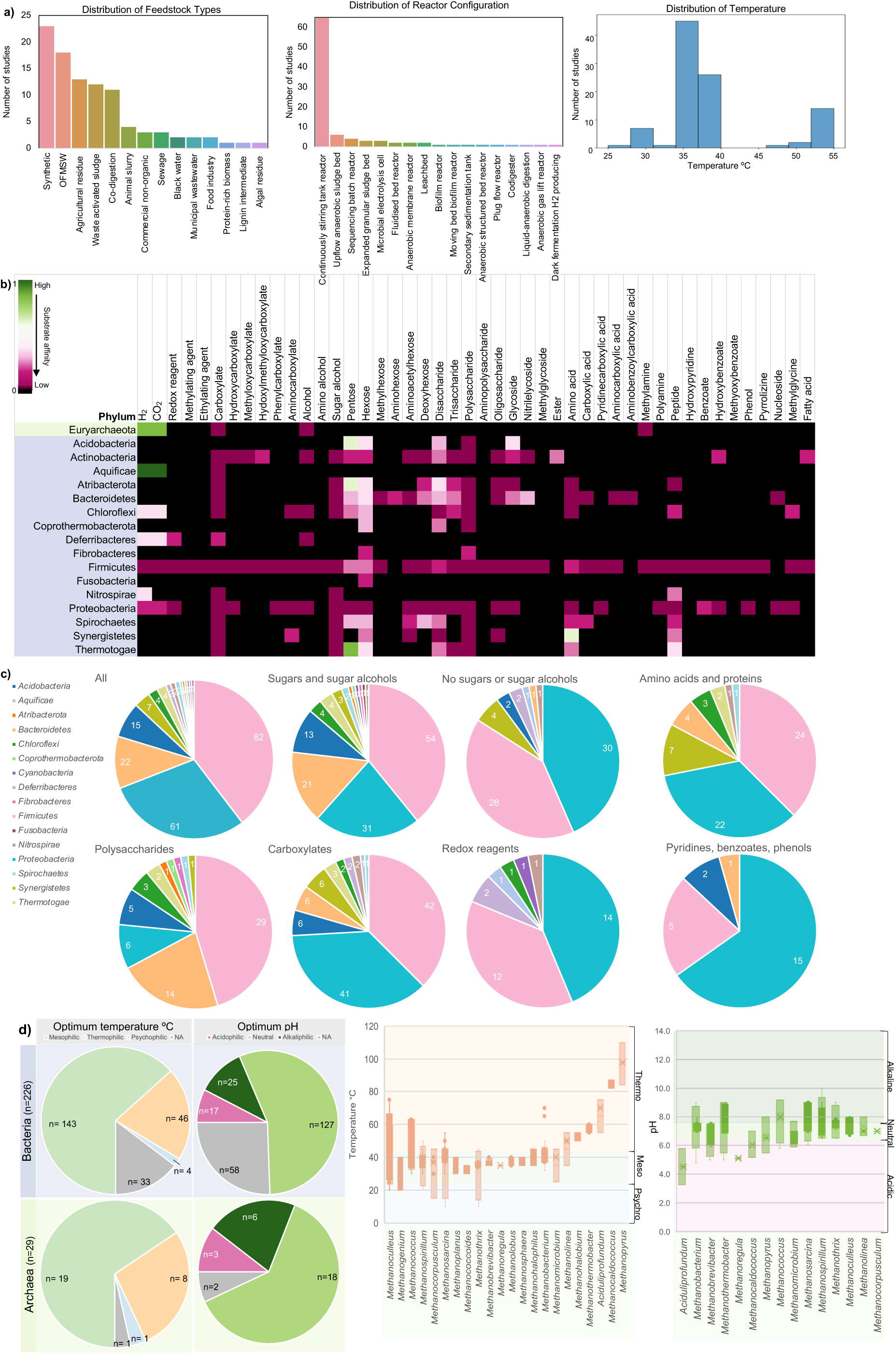
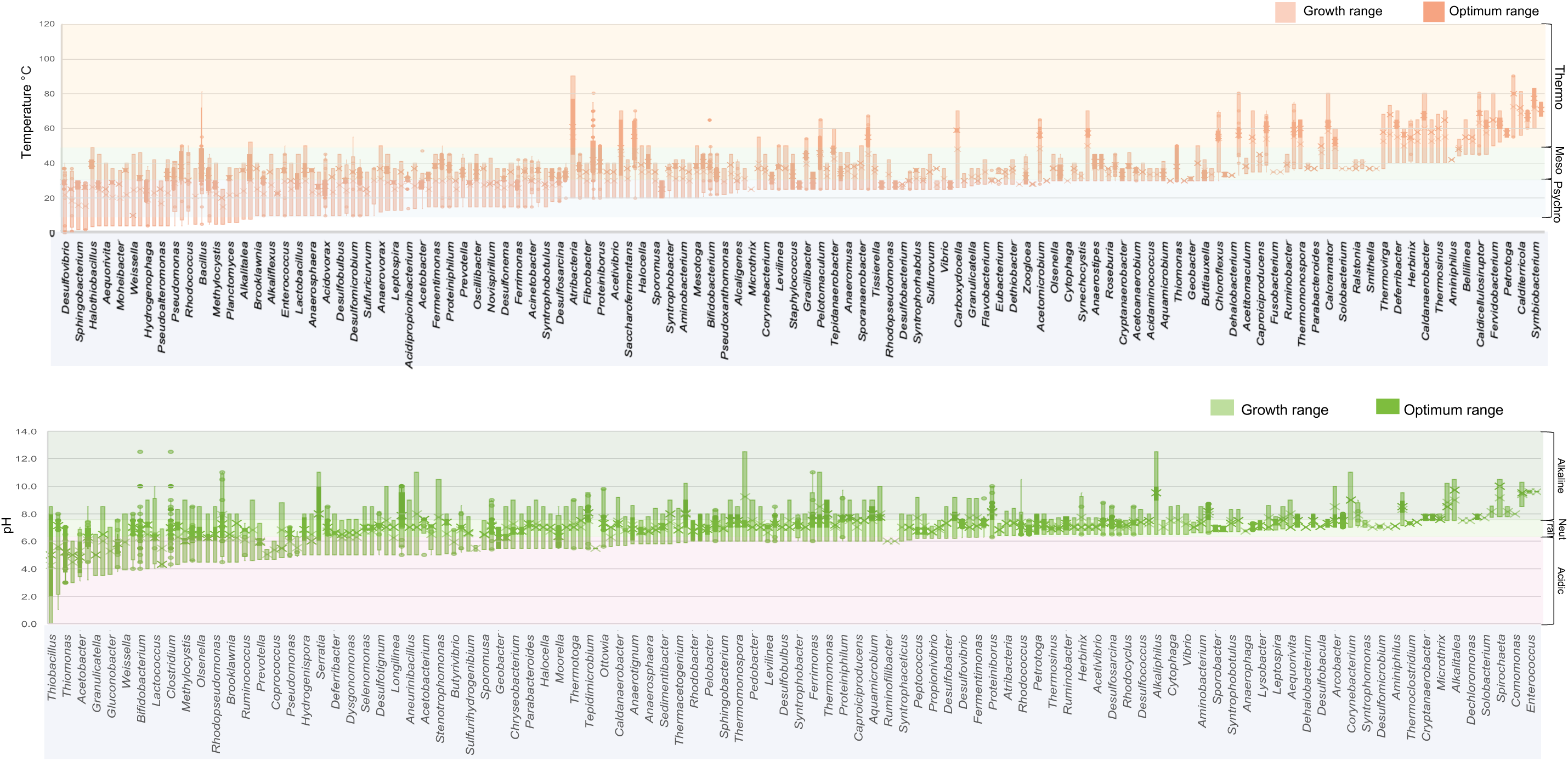
Growth conditions reported to support microbial growth for anaerobic digestion from a literature review of 197 papers (supplementary materials and methods 1.1). **(a)** Distribution of feedstock, reactor configuration and operational temperature. **(b)** Substrates (187) reported to support microbial growth were investigated. Affinity was based on average substrate utilisation on a scale of 0 to 1, where 1 (green) indicates that substrate supports the growth of all species within a phyla, and 0 (black) indicates no reported growth on a given substrate. Substrates are categorised by key functional groups (Supplementary Table 1.1). **(c)** Pie charts displaying the number of genera belonging to 16 bacteria phyla reported to utilise different categories of substrate. **(d)** Temperature and pH range reported to support growth. Temperature is categorised as psychrophilic (<25°C), mesophilic (25°C> temperature >42°C) or thermophilic (>42°C). The pH range is categorised as acidic (≤ 6.5, pink), neutral (6.5< pH <7.5, light green), or alkaliphilic (≥7.5, dark green). NA (grey) indicates information not available; n represents number of genera.

The findings of a comprehensive literature review comprising 197 publications of AD species growth and experimental conditions are synthesised in Figure 1. On the phylum level, *Firmicutes* has the most diverse substrate utilisation, which could help to explain the wide diversity of *Firmicutes* reported in AD (Figure 1b). The majority of *Firmicutes* and *Bacteroidetes* species utilised sugars, sugar alcohols, polysaccharides and carboxylates. Specific substrate categorisation is detailed in according to Supplementary Table 1.1. Conversely, *Proteobacteria* showed complementary substrate utilisation to *Bacteroidetes*, with higher utilisation of amino acids, proteins, redox substrates (such as hydrogen and nitrates) and pyridines, benzoates and phenols (Figure 1c). These findings provide insights into how different feedstocks influence microbial community identity. However, the results are not comprehensive, and further research is required to relate detailed feedstock chemical characterisation with various microbial enrichments.

Operational parameters play a crucial role in AD performance and stability. In industrial-scale AD plants, microbial communities are subjected to fluctuating conditions, including feedstock structure and composition, temperature, and pH variations ^7,8^. These fluctuations could lead to community changes, potentially causing process instability ^9^. Most species had a mesophilic optimum temperature (63%); however, many species had a wide range of temperatures that could support growth (Figure 1d). Most bacteria (56%) had an optimum pH range at or close to neutral, although one-quarter of species did not have a specified pH growth range (Figure 1d). Archaea had a higher proportion of alkaliphilic species compared to more acidophilic bacteria (21% versus 11%, respectively, Figure 1d).

Furthermore, machine learning (ML) is a rapidly expanding field that is being applied to human microbiome studies ^10,11^. Nevertheless, the application of ML in applied and industrial microbiology remains an emerging field. Current studies have shown promise for predicting AD performance from operational and biological parameters ^10^. However, these studies suffer from limited sample sizing (between 17 to 50 samples), potentially limiting statistical power, and also lack detailed feedstock and biological characterisation ^11,12^. Furthermore, model interpretability is often limited due to the need to reduce high dimensionality datasets. Applying explainable ML techniques, such as Shapley Additive exPlanations (SHAP) values, to datasets including detailed chemical and biological fingerprinting, allows interpretability of factor impact to improve our understanding of how operational, chemical and biological factors contribute to AD reactor performance.

To address this knowledge gap, our research aimed to understand the resource recovery potential of complex wastewater streams from fermentation processes and investigate the dynamics of the microbiome underpinning AD across different operational parameters and temporal scales. Here, we present the results of experiments exploring the effect of feedstock composition on AD using characterised mycoprotein fermentation wastewater (MFWW) obtained from the food industry.

## 2. Materials and methods

MFWW, a by-product of Quorn™ food production, was obtained from Marlow Ingredients (Billingham, UK). Four batches of MFWW were characterised to evaluate batch variation and the effect of the feedstock on AD (Figure 2a). Two experiments were designed to address the research objectives. In the first experimental run, mesophilic and thermophilic reactors were compared to evaluate the effect of temperature. In run 2, single- and two-stage reactors were assessed to determine the impact of reactor configuration (Figure 2b-c). Influent composition was monitored in a time-series sampling schedule (Figure 2d). Reactor chemistry, performance and microbiome biodiversity were analysed to address how MFWW composition affects these parameters across different operational parameters and temporal scales. Predictive ML models were developed to forecast AD reactor performance in terms of chemical oxygen demand (COD) removal, cumulative biogas and specific daily biogas based on biological, chemical and operational parameters.

**Figure 2.**
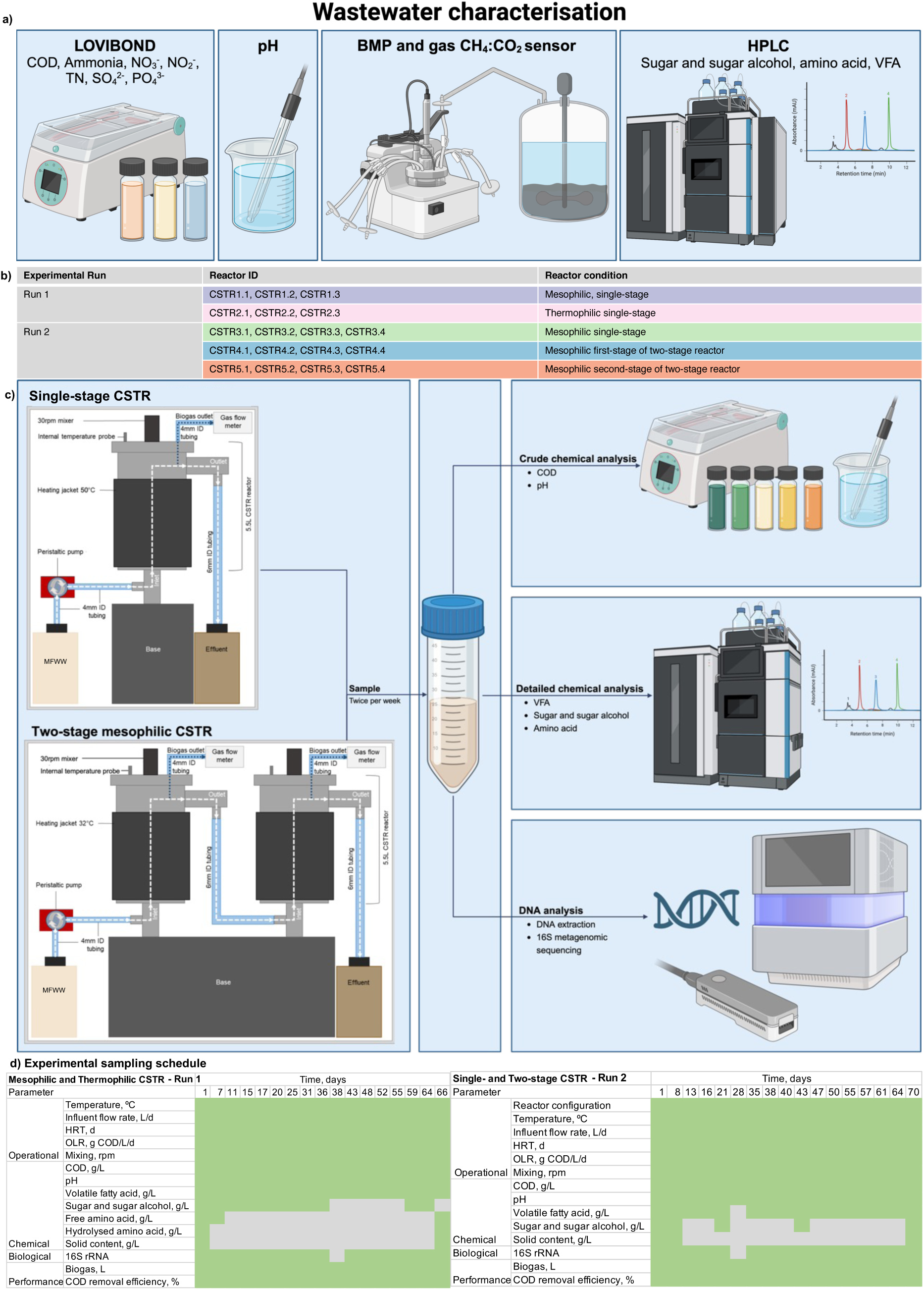
Experimental design. **(a)** Mycoprotein fermentation wastewater characterisation. **(b)** Experimental grouping. **(c)** Bioreactor set up, sampling and analysis. **(d)** Sampling schedule. *Abbreviations: chemical oxygen demand (COD), total nitrogen (TN), biomethane potential (BMP), high performance liquid chromatography (HPLC), volatile fatty acid (VFA), and continuously stirred tank reactor (CSTR).

### 2.1. Digester description

Eighteen 5.5L continuously-stirred tank reactors (CSTR) were operated with a 14.7-day hydraulic retention time and 0.32L/d flow rate with MFWW (Figure 2c). Reactors were operated for a two-month adaptation period with MFWW as feedstock at 0.18L/d. CSTR were grouped by operational conditions. Group 1 represented single-stage mesophilic reactors (37±1°C), group 2 represented single-stage thermophilic reactors (50±1°C) and group 3-5 represented mesophilic single-, first- and second-stage reactors, respectively (Figure 2b). Groups 1 and 2 contained biological triplicates, and groups 3, 4 and 5 contained four biological replicates each. 2mL samples were taken in a time-series schedule in triplicate, and operational parameters were monitored throughout (Figure 2d). Samples were evaluated using the methods detailed below. Groups 1-2 and 3-5 were set up as two independent runs using the same batch of inoculum obtained from a commercial food-waste AD plant treating municipal source-separated domestic kitchen waste, industrial food waste and a fraction of abattoir waste (Devon, UK). Volatile suspended solid content was quantified as a measure of active biomass using standard methods (Supplementary Figure 2.1) ^13^.

### 2.2. Analytical methods

#### 2.2.1. Biogas production

Real-time biogas production and composition (CH_4_ and CO_2_) from each individual reactor was measured in real-time using an Anaero technology gas flow meter and sensor.

#### 2.2.2. Chemical oxygen demand (COD)

COD was measured directly from MFWW and reactor samples using Vario HR/COD, MR/COD, LR/COD Lovibond kits (as appropriate), according to standard methods (ISO, 1989; Supplementary materials and methods 2.3). COD removal was calculated using Supplementary Equation S3.

#### 2.2.3. pH

Fresh MFWW and reactor samples were measured directly in triplicate with a pH meter (Mettler Toledo).

#### 2.2.4. High performance liquid chromatography

Analysis of sugars and sugar alcohols (melibiose, glucose, maltitol, glycerol, mannitol and arabitol) and VFAs (acetic, propionic, isobutyric, butyric, isovaleric and isovaleric acid) for MFWW and reactor samples were conducted using a Shimadzu U-HPLC with SPD-M40 photodiode array and RID-20A refractive index detectors (Shimadzu). Prior to analysis samples were filtered through a 0.22µm pore filter (Claristep® Filtration system, Sartorius). Analysis was carried out in triplicate, and the mean and standard deviation were calculated. Calibration curves were constructed by plotting peak area against concentration. The linearity of the line of best fit was calculated using the least square regression method within the LabSolutions software. The limit of detection and quantification were calculated using LabSolutions software to verify quality. Detailed methodologies are provided in Supplementary Materials 2.4.

### 2.3. Metagenomic Sequencing and Bioinformatics

#### 2.3.1. DNA extraction

Reactor samples were centrifuged at 10,000rpm for 5 minutes to obtain a pellet which was used for the DNA extraction, using the DNeasy® PowerSoil® Pro Kit according to standard protocol (QIAGEN GmbH, Germany). PCR amplification was carried out using 16S rRNA primers (Supplementary Figure 2.5.1c). Amplicon barcoding PCR was conducted using BiomekFX robot and automated software. Sequencing was conducted using Illumina MiSeq v3 kit and the run quality was assessed based on quality score (% bases>Q30), raw cluster density (k/mm^2^), data output (Gbp), read number and % PhiX aligned (Supplementary Figure 2.5.1).

#### 2.3.2. Sequence analysis

Taxonomic classification was conducted using the Mothur pipeline in Galaxy referenced against the Silva_v4 database with ≥97% sequence identity threshold ^14,15^. Additional pre-processing steps were conducted to filter for sequences ≥1200bp with Phred qualities >20 per base (Supplementary Figure 2.5.2).

#### 2.3.3. Phylogenetic tree construction

A Newick tree of genera was constructed from the taxonomic classification data using the anytree library in python and visualised using the international tree of life ^16^.

#### 2.3.4. Network analysis

The relative abundance of classified microbial taxa for each reactor sample were analysed across all time points, using the Molecular Ecology Network Analysis pipeline (Supplementary Materials and Methods 2.6) ^17^. Networks were visualised in Cytoscape at the genus level ^18^.

#### 2.3.5. Alpha diversity

Shannon, Simpson’s and Chao1 diversity indices were calculated using Supplementary Equation S4-6.

#### 2.3.6. Beta diversity and dissimilarity index

The calculation of Bray-Curtis similarity and db-RDA was implemented using the scipy.spatial, skbio and statsmodels library in python (Supplementary Eq. S7) ^19^. The dissimilarity matrix served as the response variable, while environmental variables were used as predictors. Detailed methodology is available in Supplementary Materials 2.8.

### 2.4. Machine learning models

Lasso regression, random forest and bagging regression models were evaluated using algorithms defined in Eq.S9-11 (Supplementary Materials 2.9). The models were evaluated based on their performance in predicting reactor performances. Root mean squared error (RMSE), and coefficient of determination (R²) were used to assess performance.

### 2.5. Statistical analysis

For normally distributed data with two comparisons a pairwise t-test was applied, for three or more comparisons a one-way ANOVA analysis was performed with Tukey *post-hoc* analysis. For non-normally distributed independent data a Kruskal-Wallis H test was conducted with Dunn’s *post-hoc* test for significant results. Confidence intervals of 95%, 99% and 99.9% were used. To investigate correlations between syntrophic relationships, a non-parametric Spearman’s rank correlation analysis was conducted.

## 3. Results

### 3.1. MFWW represents a complex feedstock suitable for AD

The composition of four different batches (A-D) of MFWW, compared with a previously reported value (sample R) obtained from the food industry, were measured over the time course of four fermentation cycles to determine resource recovery potential (Figure 3). We quantified physicochemical parameters driving the AD process, including ammonium, protein, pH, COD, and soluble organic compounds in MFWW (Figure 3). MFWW presented as a complex but balanced feedstock with a neutral pH (6.2±0.1) and high potential for AD (Figure 3a). Inhibitory ammonium was present at low concentrations (0.03±0.01g/L, Figure 3b). High COD (11.32±4.09g/L) and a mixed sugar profile, paired with high protein content (2.19±1.31g/L) facilitating good C/N ratios and suggests high resource recovery potential (Figure 3c-e). Biomethane potential assays showed high biodegradability (92±6%, Figure 3f).

**Figure 3.**
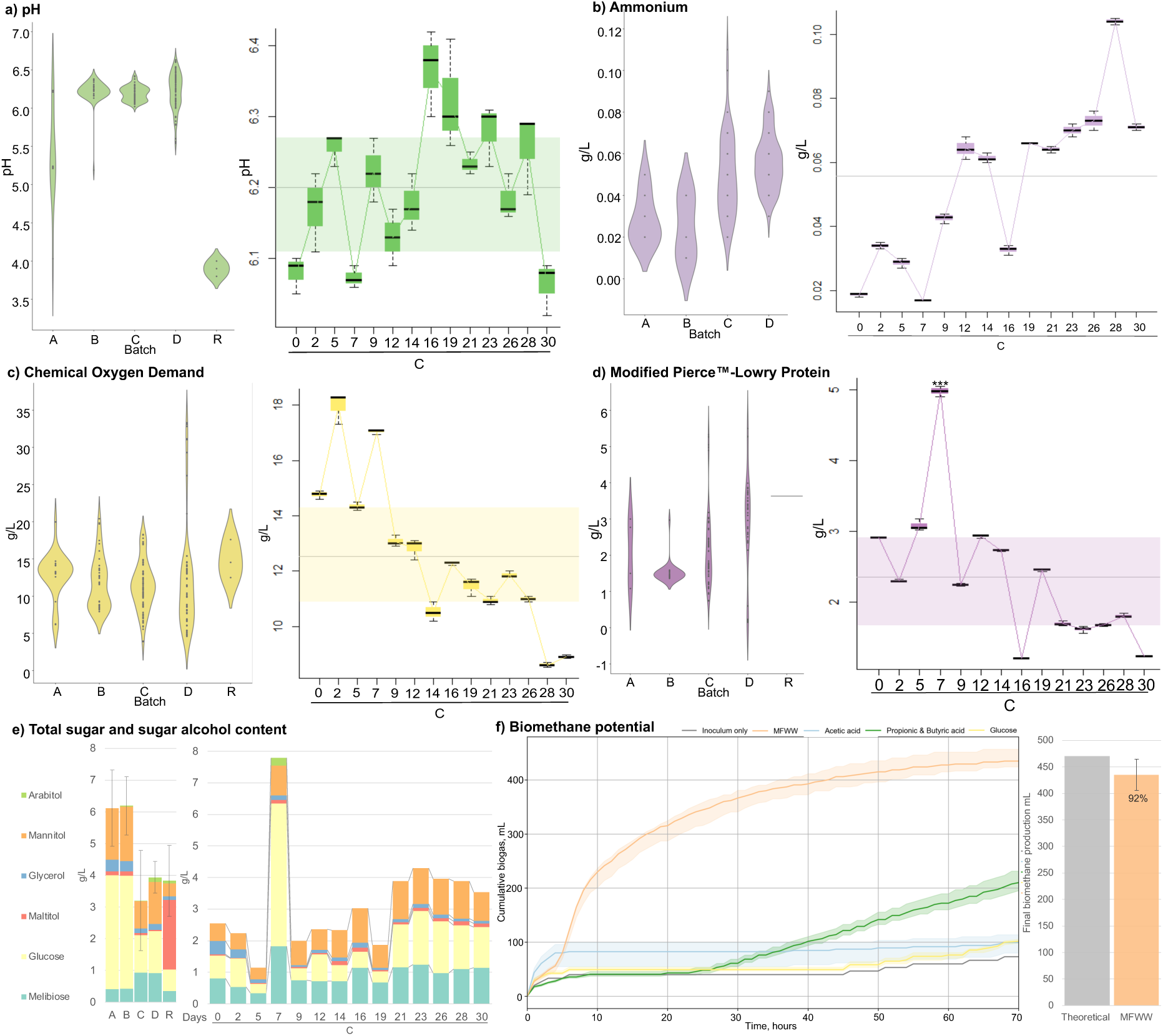
Chemical characterisation for four different batches (A-D) of mycoprotein fermentation wastewater (MFWW) compared to previously internally reported values (batch R). Batch C was measured at 14 times points within a 30-day fermentation cycle. Violin plots represent the distribution of results, box plots represent the interquartile range, the central bar represents the mean average, error bars represent the standard deviation and dots represent outliers. Statistical significance compared to R are shown at 95% *, 99% ** and 99.9% *** confidence intervals^20,21^. **(a)** pH. **(b)** Ammonium. **(c)** Chemical oxygen demand. **(d)** Protein. **(e)** Sugars and sugar alcohols. **(f)** Biomethane potential assay results including cumulative biogas of MFWW compared to inoculum only, acetic acid, propionic and butyric acid, and glucose. Final biomethane production compared as a percentage of the theoretical value calculated based on the COD to CH_4_ mass balance equation. Error bars represent standard deviation.

### 3.2. Bioreactor performance

#### 3.2.1. Run 1 reactors had better COD removal but poorer biogas production than Run 2 reactors

COD removal in samples obtained from run 1 reactors were consistently greater than the 75% regulatory threshold and run 2 reactor samples (p=0.000, Figure 4b) ^22^. COD removal efficiency in run 2 reactor samples regularly fell below the regulatory threshold, particularly at the beginning of the experiment up to day 28 and again at day 38, but gradually increased over time (Figure 4b). Conversely, run 2 reactors produced higher cumulative biogas than run 1 (p=0.000, Figure 4b). We compared the chemical and biological differences between reactors with high and low biogas production (Figure 4-5).

**Figure 4.**
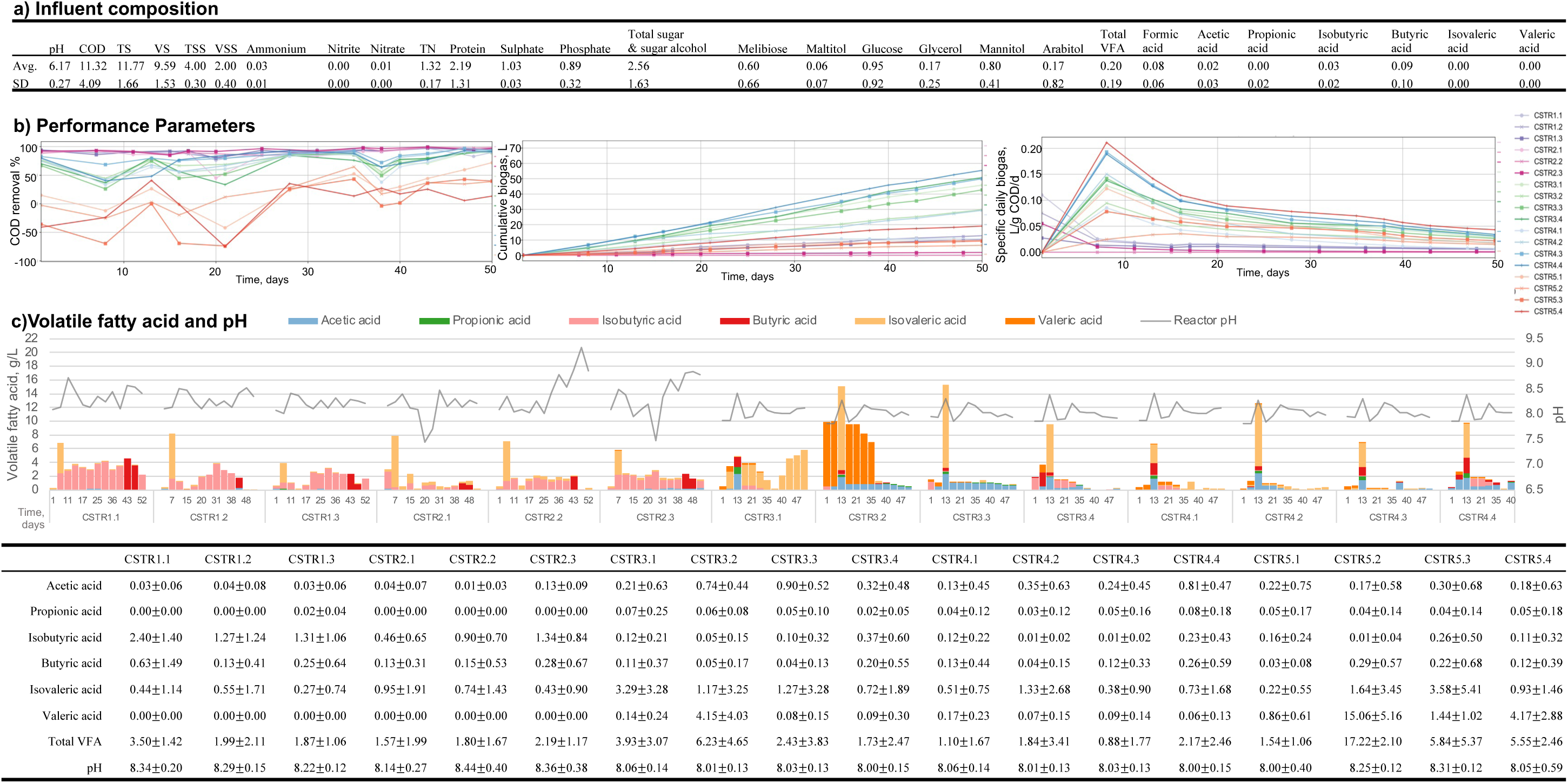
Reactor operation, performance and chemistry **(a)** Influent composition including mean (Avg.) ± standard deviation (SD) for pH, chemical oxygen demand (COD), total solids (TS), volatile solids (VS), total suspended solids (TSS), volatile suspended solids (VSS), ammonium, nitrites, nitrates, total nitrogen (TN), protein, sulphate, phosphate, total sugar & sugar alcohol, melibiose, maltitol, glucose, glycerol, mannitol, arabitol, total volatile fatty acid (VFA), formic acid, acetic acid, propionic acid, isobutyric acid, butyric acid, isovaleric acid and valeric acid. All units are g/L (except pH). (**b)** Performance parameters including COD removal, cumulative biogas, and specific daily biogas. (**c)** Reactor pH and VFA content. Bar charts represent the cumulative sum of individual VFA over time (days). Line chart represents pH (grey). Mean values ± standard deviation are presented for VFA concentrations and pH.

**Figure 5.**
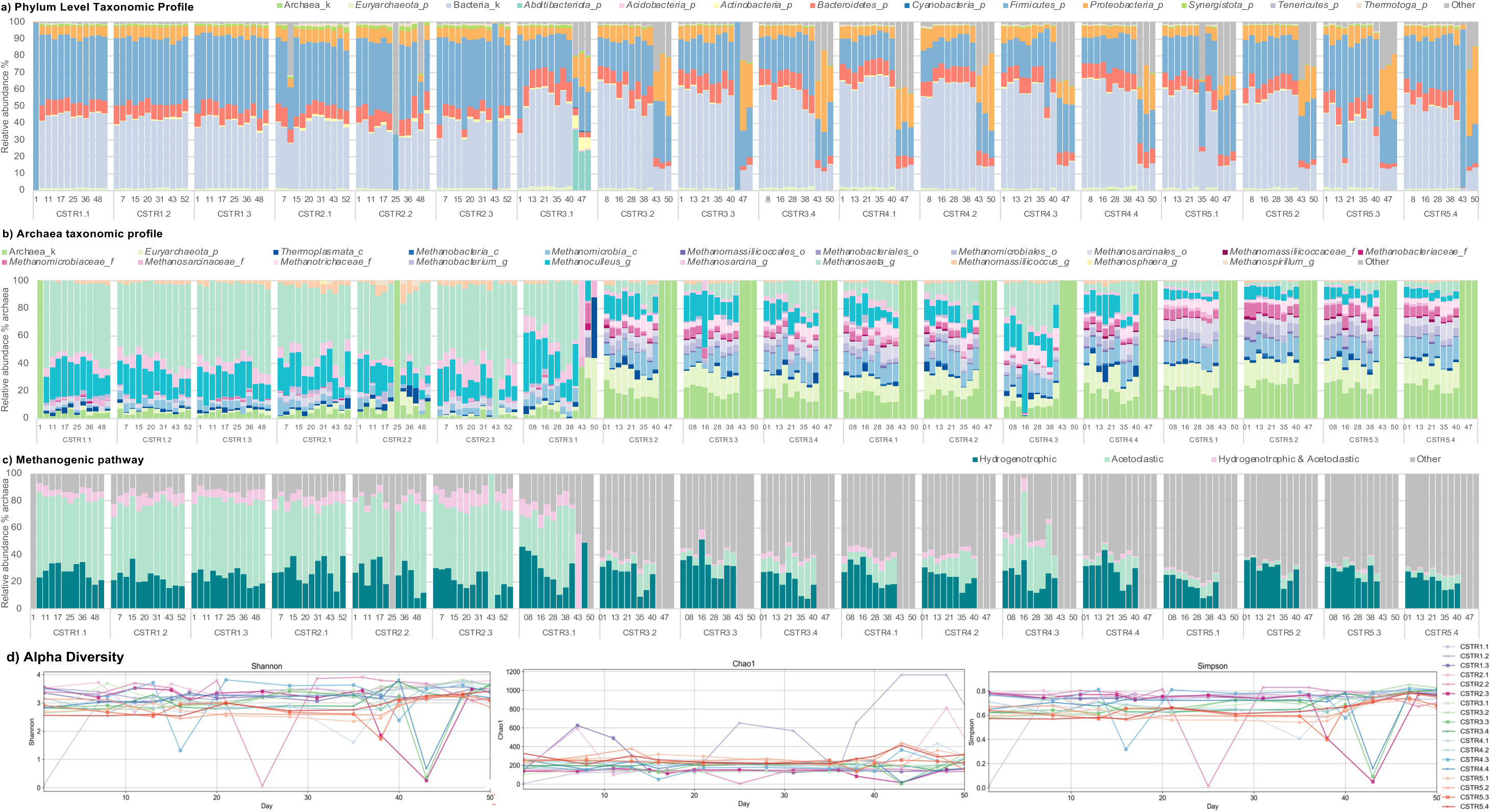
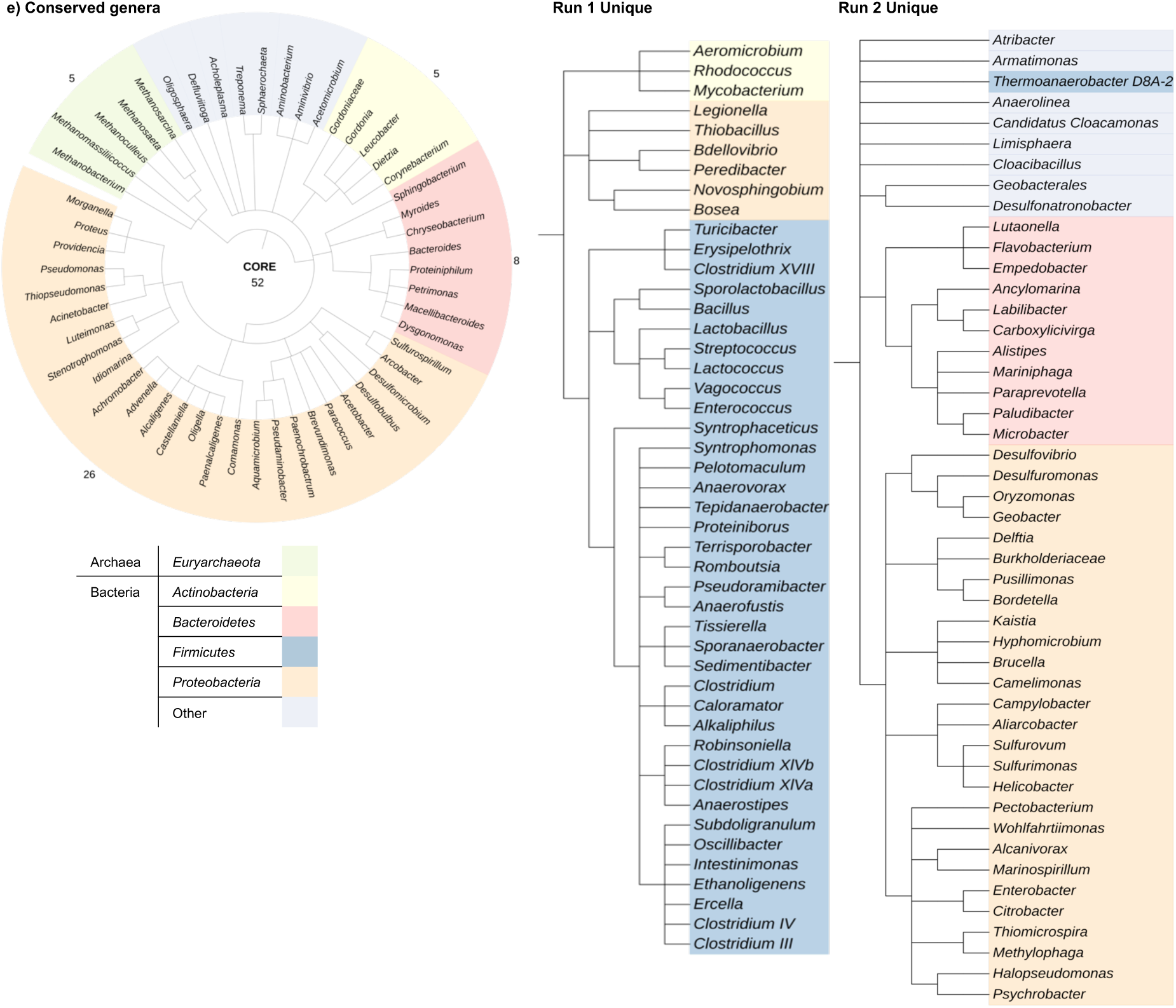
CSTR taxonomy **(a)** Phylum level relative abundance (% of total population), phyla with >1% relative abundance for at least one sample are represented and phyla representing <1% of the total population for any sample are aggregated into ‘other’ **(b)** Relative abundance of archaea (% of total archaea population). Taxa with >1% relative abundance for at least one sample are represented and taxa representing <1% of the total population for any sample are aggregated into ‘other’. Taxa are represented at the kingdom (_k), phylum (_p), class (_c), order (_o), family (_f) and genus (_g) level. **(c)** Relative abundance of archaea categorised by methanogenic pathway, where other represents ambiguous taxa. **(d)** Alpha diversity for Shannon, Simpson’s and Chao1 diversity indices over time, days. **(e)** Core microbiome of species shared across all reactors and unique species from experimental run 1 and 2. Archaea are highlighted in green, *Actinobacteria* in yellow, *Bacteroidetes* in red, *Firmicutes* in blue, *Proteobacteria* in orange, and other bacteria are light purple.

Mesophilic reactors had better biogas production than thermophilic (p=0.000) despite similar COD removal (p=1.000, Figure 4b). This result indicates that thermophilic conditions created an unfavourable environment for biogas production, possible due to poor adaptation increased temperature (Figure 4b). This could be indicative of the inoculum coming from a mesophilic source, meaning that thermophilic genera were underrepresented. Another possibility is that the temperature was low enough to support mesophilic genera with thermotolerant traits, but caused thermal stress, leading to underperformance, suggesting a specialised microbiome with little flexibility for adaptation (Figure 4b).

#### 3.2.2. VFA content showed marked differences between Run 1 and 2 reactor samples

Samples obtained from run 2 had elevated VFA concentration compared to run 1 (p=0.000) and were characterised by higher acetic (p=0.000), isovaleric (p=0.020) and valeric acid (p=0.000, Figure 4c). Isobutyric acid constituted a large proportion of the VFA content produced for all run 1 reactors, likely owing to the influence of the influent containing amino acids, such as valine and leucine, which can be converted during AD to isobutyric acid and isovaleric acid, respectively ^23–25^. The influent contained relatively high concentrations of isobutyric acid and isovaleric acid (Figure 4a) ^26^.

CSTR3.1-3.2, had elevated total VFA contents compared to CSTR3.3-3.4 (p=0.007 and p=0.001, respectively, Figure 4c). CSTR3.1 had elevated isovaleric acid, and CSTR3.2 had significantly elevated valeric acid compared to other group 3 reactors (p=0.000, Figure 4c). Samples in run 1 and 2 reactors had a spike in VFA content on day 7 and 13, respectively (Figure 4c). This peak consisted of elevated isovaleric content in samples for most reactors (except CSTR3.1) and increased acetic acid for run 2 reactors (Figure 4c). For second-stage reactors CSTR5.2, had elevated valeric acid (p=0.000), whilst CSTR5.1 had lower total VFA content than other group 5 reactors (p<0.05, Figure 4c).

The average reactor pH ranged from 8.00 to 8.44 and was slightly higher for groups 1, 2 and 5 than 3 and 4 (p=0.000, Figure 4c). Group 2 reactors pH showed high variability, suggesting potential instability (Figure 4c). Reactor pH is a key factor influencing microbial activity, and different pH ranges support different species’ growth (Figure 1d). Therefore, it will be interesting to investigate the effect of this pH difference by comparing microbial communities between reactors (Figure 1d, Figure 4c).

### 3.3. Microbiome composition and function

#### 3.3.1. Run 1 reactor samples had a distinct taxonomic profile, characterised by increased *Methanosaeta* compared to run 2 samples

In total, 1,982,343 unique sequences were retained after quality control over 25 times points. Archaea accounted for 1.23±0.72% of the total population. *Proteobacteria, Firmicutes, Bacteroidetes,* and *Actinobacteria* represented the major phyla across all reactors, although run 1 had a distinct taxonomic profile compared to run 2 reactors (Figure 5a). Notably, the relative abundance of *Firmicutes* was significantly elevated in run 1 compared to run 2 reactors, indicating elevated *Firmicutes* is associated with reduced biogas production (p<0.05, Figure 5a). Additionally, the relative abundance of *Synergistota* was significantly elevated in group 1 reactors compared to groups 2, 3 and 4 (*p*=0.000, Figure 5a).

#### 3.3.2. The *Firmicutes* taxonomic profile was highly dynamic

Core microbiome construction and times series network analysis revealed that *Firmicutes* was highly dynamic, with little conservation over time for run 2 reactors with high biogas production (Figure 4b, Figure 5e). This result suggests that a dynamic *Firmicutes* community improves reactor performance (Figure 5e). Interestingly, *Firmicutes* had the most diverse substrate utilisation of key AD phyla, which could likely be contributing to their dynamic profile (Figure 1b). By contrast, there was a much higher presence of conserved *Proteobacteria* and *Bacteroidetes* across run 2 reactors that were not shared by run 1 (Figure 5e).

#### 3.3.3. Taxonomic profile appeared to be decoupled from reactor performance

Despite having the same feedstock, operational conditions and starting inoculum, CSTR3.1 had a distinct taxonomic profile compared to other run 2 reactors, but comparable biogas production and COD removal (Figure 4b, Figure 5). Run 1 reactors had a higher abundance of *Methanosaeta* (*p*=0.000), corresponding to lower acetic acid concentration, suggesting active acetoclastic methanogenesis (Figure 5b). *Methanosaeta* is an obligate acetoclastic methanogen associated with low organic loading rate (OLR), acetic acid, and solid and ammonium contents^27^. Additionally, *Methanosaeta* are reported to have an alkaliphilic optimum pH of 7.0 to 9.0 (Figure 1d). The higher pH and lower acetic acid concentration of run 1 compared to group 3 and 4 could explain the elevated *Methanosaeta* (Figure 4c, Figure 5b).

Notably, CSTR3.1 had similar performance to run 2 reactors but a taxonomic profile that was more comparable to run 1; for example, CSTR3.1 and run 1 reactors had comparable *Methanosaeta* abundance (Figure 5). This result indicates that reactor performance was decoupled from the microbiome profile. Despite low OLR, acetic acid concentration and high pH, *Methanosaeta* content was reduced in group 5 reactors, indicating potentially less reliance on acetoclastic methanogenesis (Figure 5b-c). However, second-stage reactors had a lower percentage of classified genera compared to first- and single-stage reactors making it difficult to conclude the true relative abundance of acetoclastic methanogens (Figure 5).

#### 3.3.4. Despite some fluctuations, reactors had similar and stable species richness and evenness

Shannon, Simpson’s and Chao1 alpha diversity indices were compared as measures of community evenness and richness (Figure 5d). There was a significant drop in Shannon and Simpson’s diversity for only CSTR3.3 on day 43, which recovered by day 48 (Figure 5d). By day 28, Simpson’s diversity was similar for all reactors (Figure 5d). Chao1 diversity was similar for CSTR1.1 and Group 3; however, CSTR1.2 and CSTR1.3 showed large fluctuations, resulting in periods of elevated Chao1 diversity index values (Figure 5d). Reactor CSTR1.2 had elevated Chao1 diversity past day 20, whereas CSTR1.3 had elevated Chao1 diversity between days 1 and 15, indicating more rare species than other reactors (Figure 5d). Species richness and evenness were reduced in group 5 reactors compared to group 1-4 reactors, indicating that alpha diversity plays a crucial role in reactor resilience to stress conditions. This result agrees with the literature that diversity improves performance (Figure 5d) ^28^.

### 3.4. Microbiome dynamics and interaction with bioreactor parameters

#### 3.4.1. Denser networks were associated with increased biogas production

Run 1 reactor samples were found to represent less dense networks than those in run 2 (Figure 6a). These results suggest that network density contributes to reactor performance, supporting previous findings from Dalantai *et al.,* ^29^, who reported that sparser networks were associated with reactor instability. Time series analysis of node centrality showed that groups 3 and 5 had a higher node degree than groups 1, 2 and 4 (Figure 6b). Group 4 showed greater variation in node betweenness and a lower clustering coefficient than other reactor groups (Figure 6a). Interestingly, run 1 had a higher node event centrality than run 2, suggesting that node event centrality could be important in reactor performance (p= 0.000, Figure 6b).

**Figure 6.**
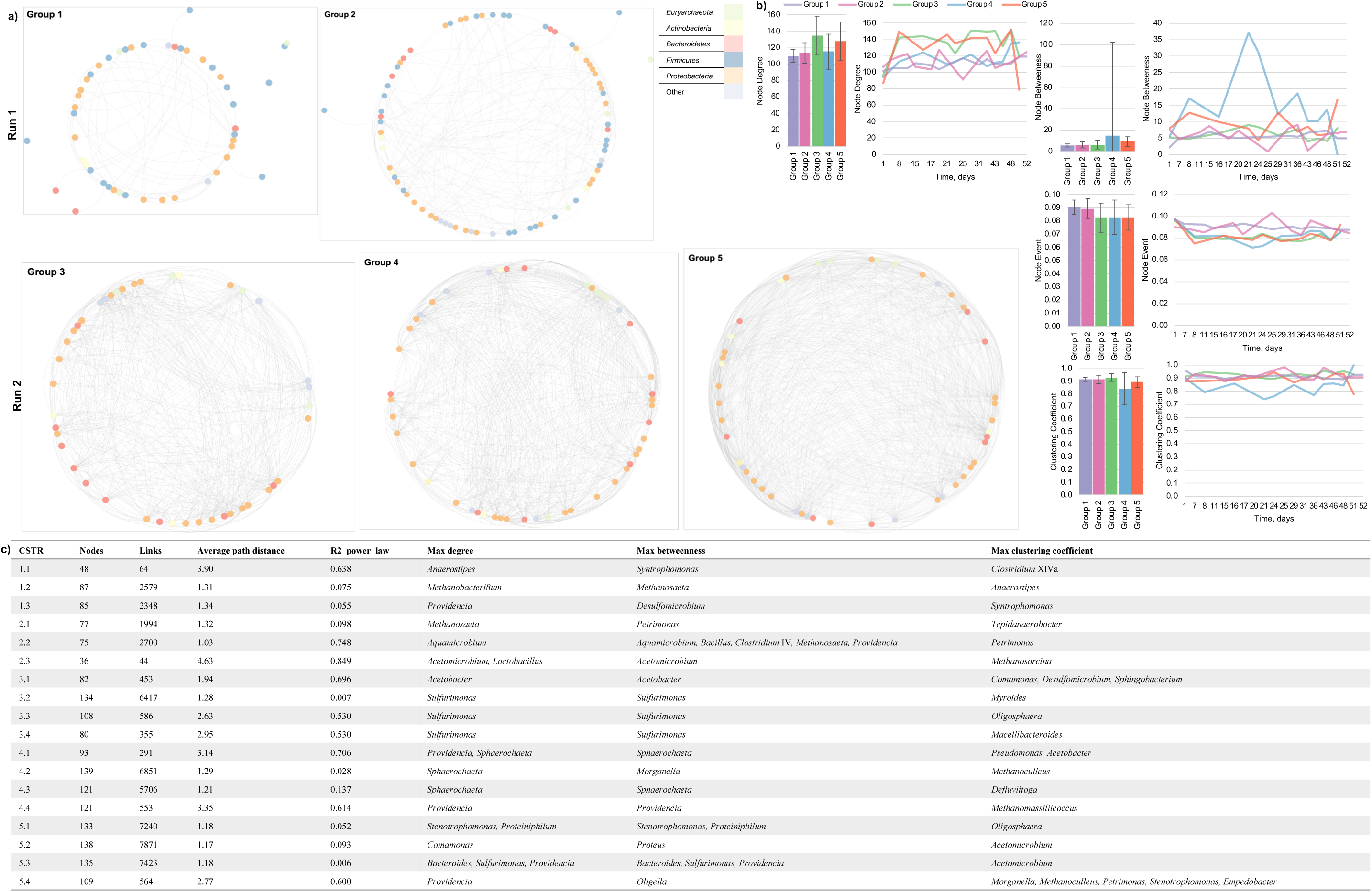
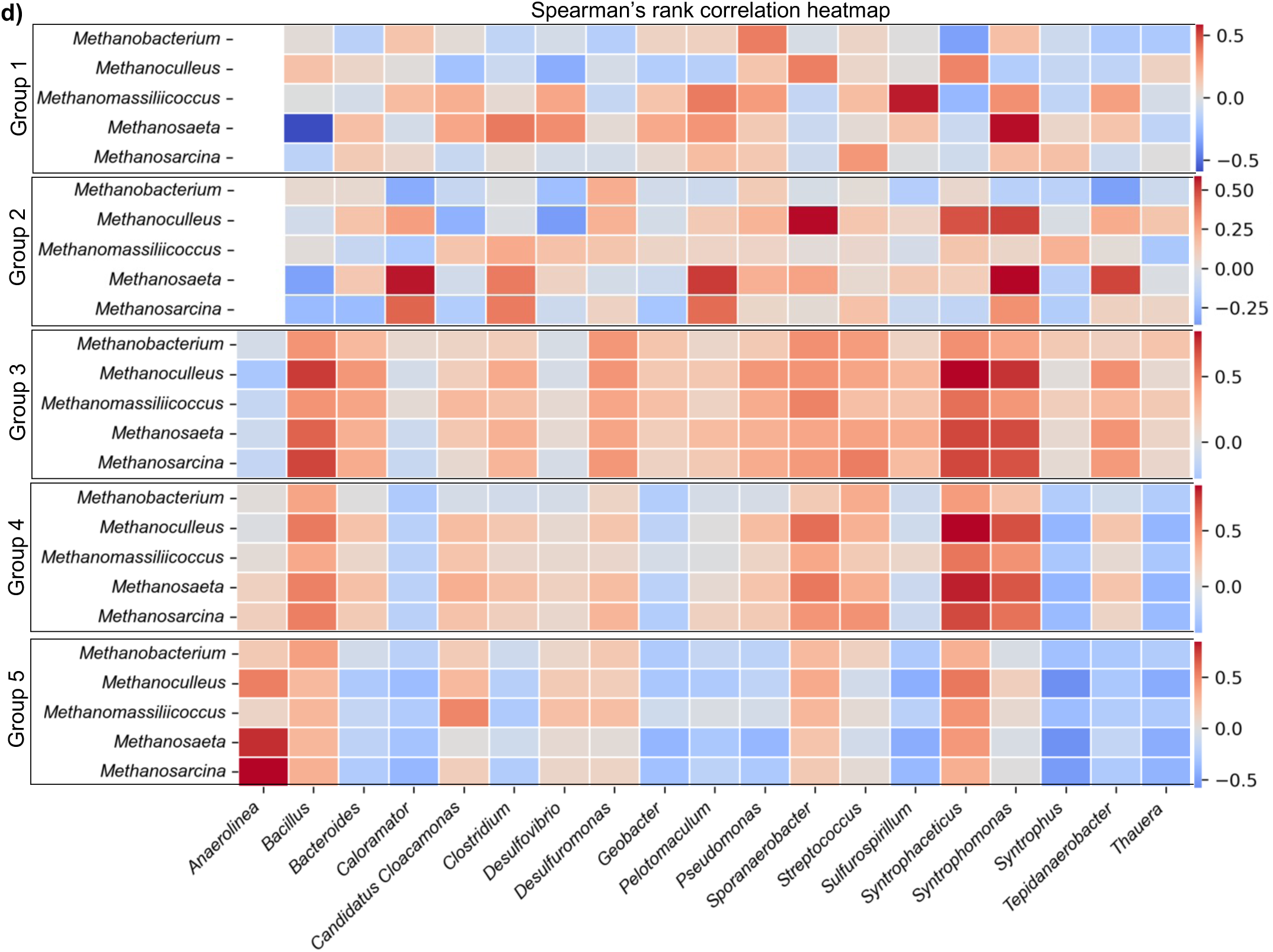
Microbiome interaction analysis. **(a)** Time series network analysis of each reactor group at genus level, where nodes represent genera. **(b)** Node centrality is presented where bar charts represent the average, and error bars represent the standard deviation, and line charts represent node centrality over time for node degree, betweenness, clustering coefficient and event centrality. **(c)** Descriptive statistics of network analysis and the identity of nodes with maximum degree, betweenness and centrality are described. **(d)** Spearman rank correlation coefficient of dominant methanogens and syntrophic genera presented as a heatmap for group 3-5 reactors, where *c*orrelations are reported as positive (red) or negative (blue). Blank values indicate insufficient data. Only correlations with sufficient data for at least one genera are presented.

#### 3.4.2. Syntrophic relationships were associated with increased biogas production

Syntrophic bacteria and methanogens, including *Syntrophomonas, Methanosarcina* and *Methanosaeta*, represented group 1 central network nodes (Figure 6c). CSTR2.3 also had high connectivity of syntrophic genera (Figure 6c) and elevated biogas production compared to other group 2 reactors, indicating the potential importance of syntrophic genera connectivity in reactor performance (Figure 4b, Figure 6c). Mesophilic reactors had elevated syntrophic genera compared to thermophilic reactors, and higher connectivity of syntrophic genera and methanogens was associated with improved biogas production. Syntrophic genera belonging to *Firmicutes*, which have been related to stress conditions, such as high OLR, were present across all run 1 reactors, which could have contributed to lower biogas production ^30,31^. Spearman’s rank correlation analysis revealed significant correlations between thermophilic *Caloramator* and *Methanosaeta* and *Methanosarcina* in thermophilic reactors only (Figure 1, Figure 6d).

Spearman rank correlation analysis revealed a positive correlation between hydrogen-producing syntrophic bacteria and hydrogenotrophic methanogens in single- (group 3) and first-stage (group 4) reactors, which was diminished in the second-stage (group 5) reactors (Figure 6d). Most syntrophic bacteria showed significant correlations to dominant methanogens for single- and first-stage reactors (p<0.05), except for *Candidatus Cloacamonas* (p>0.05, Figure 6d). However, the positive correlation between *Candidatus Cloacamonas* and *Methanomassiliicoccus* was significant for second-stage reactors (p=0.000, Figure 6d). *Anaerolinea* and acetoclastic methanogens were significantly positively correlated; however, the strength of this correlation was lower in single-compared to first- and second-stage reactors (p<0.05, Figure 6d).

The relationship between syntrophic genera were less clear in samples from second-stage reactors, which were found to have fewer significant correlations (Figure 6d). However, *Anaerolinea, Methanosarcina,* and *Methanosaeta* showed significant positive correlations (p=0.000, Figure 6d). *Desulfuromonas* and *Pseudomonas* were also significantly positively correlated to *Methanoculleus, Methanosaeta* and *Methanomassiliicoccus* and *Geobacter* was significantly correlated with *Methanoculleus* (p=0.000, Figure 6d). Importantly, samples in the group 5 reactors exhibited decreased acetic acid concentrations and acetoclastic methanogenesis, which could reduce the strength of syntrophic acetate oxidising bacteria relationships within the reactors (Figure 4c, Figure 6d). This result highlights the importance of syntrophic relationships in reactor performance.

#### 3.4.3. *Methanomassiliicoccus* was associated with increased butyric acid concentration and decreased biogas production

Group 2 reactors had a greater abundance of *Methanomassiliicoccus* than other reactors (p=0.000, Figure 5b). Interestingly, *Methanomassiliicoccus* were associated with periods of enriched butyric acid, suggesting a potential correlation between *Methanomassiliicoccus* and isobutyric and butyric acid concentrations (Figure 4c, Figure 5b). This result is consistent with data from Nikitina *et al.,* ^32^, who found that *Methanomassiliicoccus* were enriched under butyric acid dominant VFA accumulation. *Methanomassiliicoccus* has been associated with reduced biogas yields, which agrees with the results that run 1 reactors had lower biogas production than run 2 reactors (Figure 4c, Figure 5) ^33^.

### 3.5. Effects of chemical and operational parameters on bioreactor performance and biodiversity

#### 3.5.1. Random forest models performed best for predicting anaerobic digestion reactor performance

Supervised ML models were developed to investigate the potential of operational parameters and reactor chemistry and biology for predicting reactor performance in terms of COD removal and biogas production. Lasso regression models provided a baseline with moderate predictive capability. Random forest regression is a robust ensemble method that creates and merges multiple decision trees for a more accurate and stable prediction. Bagging regression is an ensemble technique that improves the stability and accuracy of ML algorithms by reducing variance and preventing overfitting. Random forest regression showed the best performance with the highest RMSE and lowest R², reflecting high accuracy and substantial variance explanation (Figure 8b-d). Overall, the evaluation revealed that the random forest outperformed the two other models across all experiments (Figure 8b-d).

**Figure 8.**
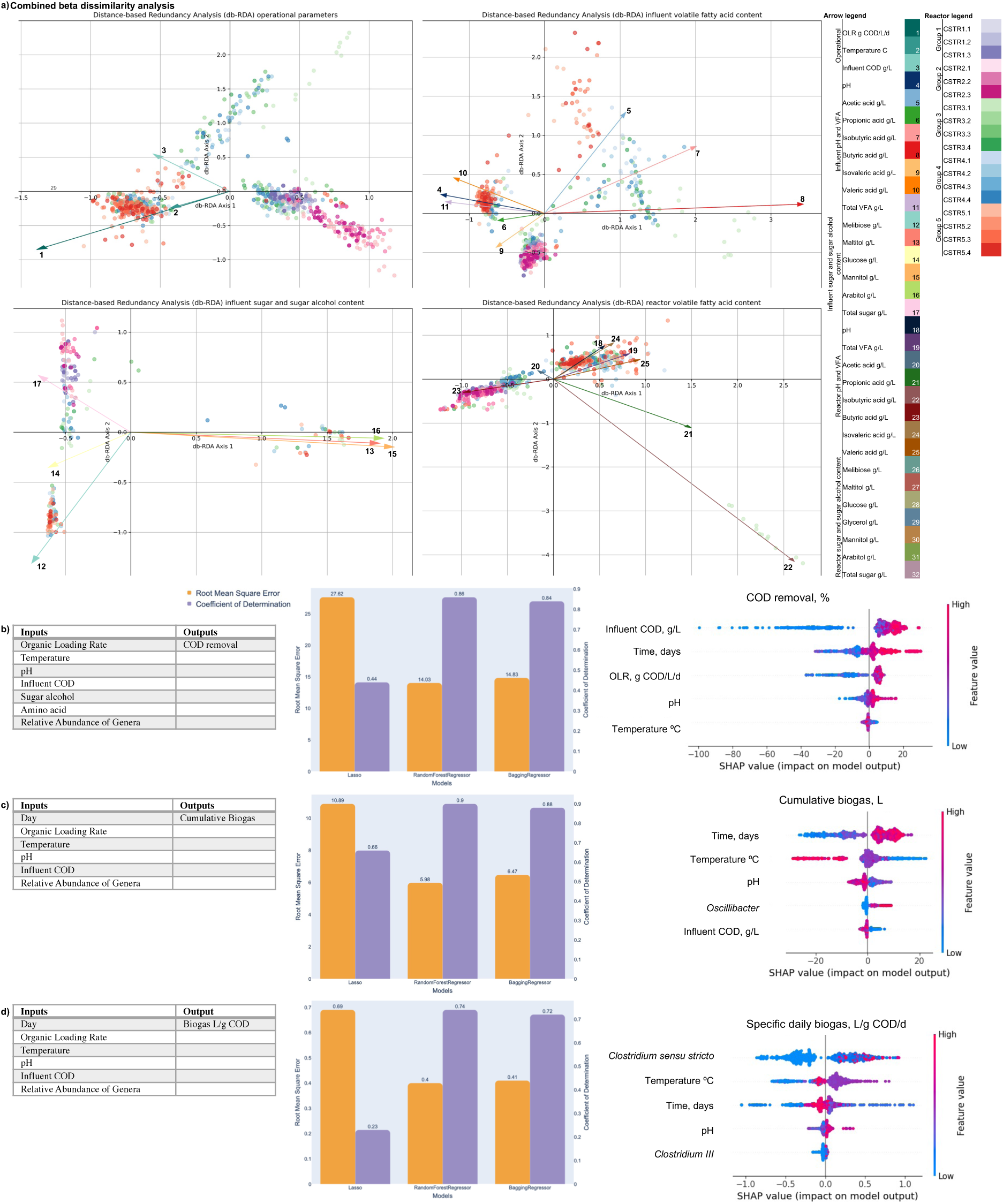
Effect of operational and chemical parameters on microbiome biodiversity and performance. **(a)** Dissimilarity analysis of mesophilic (M1 to M3, purple) and thermophilic (T1 to T3, pink) reactors assessing the influence of operational parameters, influent VFA content, influent sugar and sugar alcohol content, and reactor VFA content. Machine learning predictive models includes Lasso regression, random forests and bagging regression. The inputs and outputs of each model are described in the tables, the RMSE and R^2^ values used to evaluate the models are presented as bar charts and factor influence is displayed in terms of SHAP values for each model predicting reactor performance in terms of **(b)** COD removal. **(c)** Cumulative biogas, L. **(d)** Specific daily biogas, L biogas/g COD/d. *Abbreviations: organic loading rate (OLR), chemical oxygen demand (COD), volatile fatty acid (VFA), root mean squared error (RMSE).

#### 3.5.2. Influent COD and OLR were determined to be key influences on reactor biodiversity and performance

The average impact of features on model output was measured using SHAP values. The top five most important features affecting COD removal included influent COD, time, OLR, temperature, and pH (Figure 8b). Influent COD was the most significant feature, with the highest SHAP value (Figure 8b). The most influential factors affecting biogas production were time, temperature, pH*, Oscillibacter* and influent COD (Figure 8c). The horizontal placement of these lines reflects the SHAP values, offering a numerical assessment of their impact on the model’s results. Analysing the distribution along the x-axis reveals that higher temperature values are associated with lower cumulative biogas and decreased reactor performance (Figure 8c). Two *Clostridium* species, temperature, time, pH were the top five factors influencing specific daily biogas (Figure 8d). Overall, the predictive models emphasised the importance of operational parameters on reactor performance, and biological genera within *Firmicutes* (*Clostridium* and *Oscillibacter*) on biogas production (Figure 8b-d).

Dissimilarity analysis revealed that influent COD and OLR were the most influential operational parameters on beta diversity (Figure 8a). Influent butyric acid content, and mannitol, arabitol, and maltitol concentrations strongly influenced beta diversity (Figure 8a). Group 3 reactors were distributed across three main clusters, which appeared to be strongly influenced by individual sugars and sugar alcohols. Group 1 formed one central cluster and was strongly influenced by total sugars and sugar alcohol content (Figure 8a). Influent isobutyric, butyric, and acetic acid content appeared to strongly influence diversity (Figure 8a). Notably, individual VFAs appeared to have a more substantial influence on beta diversity than total VFA content, highlighting the importance of detailed chemical analysis for understanding microbiome diversity (Figure 8a).

Feedstock composition was identified as a critical driver of biological parameters across all reactors. The use of ML models, particularly random forests, has proven highly effective in this work. The agreement between the dissimilarity analysis and ML models emphasised the robustness of the ML models and highlighted that influent COD and OLR were highly influential on microbiome diversity and reactor performance (Figure 8).

### 3.6. Discussion

Biological repeats of various anaerobic digestion reactors fed with MFWW were monitored over a time series to evaluate the effect of feedstock composition on the microbial community underpinning anaerobic digestion across different operational parameters and time scales.

Influent COD and OLR were also identified as key influences on reactor performance by ML and dissimilarity analysis. Dissimilarity analysis showed strong clustering of reactor type by operational parameters, implying a strong impact on microbial diversity (Figure 8a). Additionally, individual VFAs were found to have a stronger influence on beta diversity than pH or total VFA, as did individual sugars and sugar alcohols compared to total sugar and sugar alcohol concentrations (Figure 8a). Isobutyric acid content in particular was found to have a major influence on microbiome biodiversity, as were influent acetic, isobutyric, and butyric acid levels (Figure 8a).

Bioreactor chemistry highlighted distinct VFA profiles between runs 1 and 2 (Figure 4c). VFA profiles in run 1 consisted of a majority of isobutyric acid, which was correlated to the relative abundance of *Methanomassiliicoccus* ^32^. The detection of both acetoclastic and hydrogenotrophic methanogens in various reactors underscores the flexibility and resilience of AD microbial communities ^34^. It is important to note that *Methanomassiliicoccus* lack the Wood-Ljungdahl pathway for methyl group oxidation to carbon dioxide ^35^. *Methanomassiliicoccus* have instead been found to use hydrogen-dependent methylotrophic methanogenesis pathways, utilising methylamines and methanol, which highlights an unusual metabolic pathway compared to classic hydrogenotrophic or acetoclastic methanogens ^36,38^.

Additionally, *Methanomassiliicoccus* have been associated with reduced biogas production ^33^, and were significantly correlated with syntrophic bacteria only in group 5, which had the lowest biogas production (Figure 6d). Therefore, this study provides further evidence to support the hypothesis that *Methanomassiliicoccus* is associated with butyric acid and reduced biogas production. Moreover, this study has also highlighted the importance of *Firmicutes* in relation to reactor performance. The relative abundance of *Firmicutes* was elevated and showed little conservation over time in run 1 compared to run 2 samples. This result implies that a relatively low abundance of *Firmicutes* and a dynamic *Firmicutes* population was associated with increased biogas production. Furthermore, ML models identified three genera of *Firmicutes* (*Oscillibacter*, and two *Clostridium*) as primary factors that influence biogas production (Figure 8c-d). These combined results implicate *Firmicutes* as key bacteria for influencing in biogas production (Figure 5, Figure 8).

A shift in taxonomic profile observed between days 43 and 50 for almost all reactors corresponded to increased influent COD and increased OLR (Figure 4c, Figure 5). Dissimilarity analysis and ML outcomes revealed that OLR had the most substantial effect on microbiome diversity, and that influent COD was a key factor in modifying reactor performance (Figure 8). The shift in taxonomy was associated with improved performance and microbiome diversity stability, indicating that flexibility within the microbiome improves performance in response to fluctuating feedstock composition (Figure 4b, Figure 5).

Despite having identical feedstock (MFWW), operational conditions and starting inoculum, CSTR3.1 had a distinct taxonomic profile compared to other single-stage reactors but comparable biogas production and COD removal (Figure 4b, Figure 5). This result indicates that reactor COD removal and biogas production was decoupled from the microbiome profile. Syntrophic relationships showed stronger correlations in single- and first-stage compared to second-stage reactors (Figure 6d). Additionally, network analysis revealed that syntrophic species had higher node connectivity in mesophilic compared to thermophilic reactors (Figure 6). Several bacterial genera are involved in syntrophic relationships essential for AD. *Syntrophobacter* converts butyrate and propionate to acetate, which is then utilised by methanogens such as *Methanosarcina* and *Methanosaeta* ^37^. Similarly, *Syntrophomonas wolfei* degrades butyrate in association with hydrogen-utilising methanogens, maintaining low hydrogen partial pressure, which is critical for the energetics of the reaction ^38^. Additionally, *Smithella* species grow on propionate only in the presence of methanogenic bacteria that remove hydrogen and formate, facilitating propionate oxidation ^36^. *Candidatus Cloacamonas*, part of the WWE1 candidate phylum, and *Syntrophomonas* are involved in the fermentation of amino acids and butyrate, respectively, indicating their significant roles in syntrophic degradation processes (Figure 1) ^39^. Furthermore, the presence of syntrophic bacteria like *Thermotoga* and *Thermacetogenium* in thermophilic digesters supports syntrophic acetate oxidation under high-temperature and high-ammonia conditions, highlighting the adaptability and importance of these bacteria in diverse AD environments ^27^. Additionally, *Cytophaga*, *Herbaspirillum*, *Symbiobacterium*, *Comamonas*, and *Allochromatium* have been associated with enhanced biogas production, although their exact roles remain unclear ^40,41^.

A greater understanding of the microbiome underpinning AD could improve reactor stability and reduce start-up times ^28,42,43^. The well-described process of acetoclastic methanogenesis has formed the basis for much top-down AD microbiome optimisation. However, our results agree with the findings that the obligate acetoclastic methanogen *Methanosaeta* has been associated with reactor dysfunction, and acetoclastic methanogenesis is not always the dominant pathway for methane production ^41,43–45^.

## 4. Conclusions and Future Directions

This study investigated the effect of complex feedstock (MFWW) from the microbial protein industry on microbiome diversity and reactor performance in a time series. ML models and dissimilarity analysis highlighted the importance of crude and detailed chemical analysis for influencing reactor biodiversity and performance. The findings also indicated that *Methanomassiliicoccu*s is associated with high butyric acid concentrations and reduced biogas production, providing new insights into the newly discovered methanogen. Finally, *Firmicutes* were implicated in reactor performance and were found to be highly dynamic in reactors with high biogas production.

Future research should focus on building upon the findings of this study by elongating the run times, including untargeted chemical analysis and additional -omics tools. HPLC analysis was limited to known compounds, including VFAs and sugars and sugar alcohols, which failed to identify all compounds of interest within the reactors. Untargeted analysis would be recommended to identify all compounds of interest within the reactors. Furthermore, individual compound manipulation of characterised feedstocks (MFWW) could be conducted to validate the findings of the dissimilarity analysis and ML models. Preliminary exploration of bioreactor chemistry and operational parameters as predictors of microbiome structure was used to investigate whether microbial community assembly is deterministic or stochastic (Supplementary Materials 3.3). However, more data will be required to further explore this topic. Moreover, advanced time series analysis was challenging to implement. An extended run time of at least 6 months would be recommended to further explore the temporal trends and whether community can be predicted based on feedstock composition and operational conditions. Finally, we recommend additional -omics to investigate the role of structure and function in relation to reactor chemistry and performance parameters. Meta-transcriptomics is a common approach used in microbiome studies to elucidate the functional profile of the microbiome and should be applied to further explore some interesting relationships observed during the experimental run, such as the metabolic basis for the elevated production of valeric acid in CSTR3.1. A meta-pan genomic approach could also be applied to compare reactor metagenomes and their relationship to bioreactor chemistry.

This work developed robust predictive ML models based on AD chemical and biological fingerprinting. The combined ML approach derived actionable insights for the potential of *Firmicutes*, *Methanomassiliicoccus* and butyric acid as biological and chemical markers to monitor and predict system performance. By integrating biological, chemical, and computational methods, researchers can develop robust monitoring methods that consider whole systems approaches to enhance AD performance. Detailed feedstock analysis in tandem with the application of metagenomic sequencing and reactor chemical analysis are key methods for capturing the complexity of these communities. By investigating substrates ranging from single molecules to complex macroparticles, we may be able uncover microbial structure and function trends. By effectively managing operational parameters and bioreactor chemistry, it is possible to harness the full potential of AD, ensuring efficient wastewater treatment and resource recovery.

## Supporting information

Supplementary Database 1

## Data Availability

All sequencing data have been submitted to the European Nucleotide Archive (ENA) under the project ID PRJEB80086. Sequences reported in this study are deposited in the EMBL-EBI database under accession numbers ERS22193437-ERS22335023. The correspondence between accession numbers, sequences and metadata is provided in Supplementary Database 1. Machine learning models are open source and available at https://github.com/MGuo-Lab/AnaerobicDigestionML.

## Author contributions

EP and MG conceived the study. EP acquired the data. EP and XS conducted the experiments, analysed and interpreted the data. EP, XS, PE, MT and MG drafted or revised the manuscript.

## Acknowledgements

We are grateful to Marlow Ingredients and the research and development team, especially Dr Mark Taylor, for supplying the feedstock for the experiments. EP and MG would like to acknowledge the UK Engineering and Physical Sciences Research Council (EPSRC) and Monde Nissin Corporate for providing fundings under the EPSRC iCASE programme.

