## Supplementary Materials and Methods for "Temporal Dynamics of Microbial Communities in Anaerobic Digestion: Influence of Temperature and Feedstock Composition on Reactor Performance and Stability"

### 1. Supplementary Introduction and Literature Review

#### 1.1. Reported substrate utilisation and categorisation

**Supplementary Table 1.1.** Functional categorisation of substrates reported to support the growth of species involved in AD. The functional categorisation was based on the molecular formula, condensed formula, ionic charge, and elemental composition in terms of carbon (C), hydrogen (H), nitrogen (N), oxygen (O), sulphur (S), chlorine (Cl) and iron (Fe), molecular weight, and the number of functional groups. Ionic charge is presented on a colour scale of red (strongly negative) to blue (strongly positive). The number of functional groups is colour-coded from light blue (low) to dark blue (high), where black represents a given functional group's absence and NA indicates information unavailable.

| | | | | | Elemental composition | | | | | | | | | | | | | | | | | | | | | | | | | | | | | | | | | | | | | | |
|---|---|---|---|---|---|---|---|---|---|---|---|---|---|---|---|---|---|---|---|---|---|---|---|---|---|---|---|---|---|---|---|---|---|---|---|---|---|---|---|---|---|---|---|
| Functional Category | Name | Molecular formula | Condensed Formula | Ionic Charge | C | H | N | O | S | Cl | Fe | Molecular weight | R-COOH, carboxylic acid | R-COO$^-$, carboxylate | R-OH, hydroxyl | R-NH$_2$, amine | R-NH | R-N-R | R-NH$_3^+$ | R-NO$_3^-$ | Nitrile, CN | R-N-(C=O)-R, amide | R-C=O, ketone | R-HC=O, aldehyde | R-O-C=O, Ester | R-CH$_3$, methyl | R-O-CH$_3$, oxymethyl | R-Cl | R-Fe$^{3+}$ | R-S | C=C | Benzene | Phenyl | Pyridine | Imidazole | Indole | Oleate | pyranose | furanose | pyrrolizidine | Glucosamine | Purine | Pteridine |
| Reducing agent | Hydrogen | $H_2$ | $H_2$ | 0 | 0 | 2 | 0 | 0 | 0 | 0 | 0 | 2.02 | | | | | | | | | | | | | | | | | | | | | | | | | | | | | | | |
| Oxidising agent | Carbon dioxide | $CO_2$ | $CO_2$ | 0 | 1 | 0 | 0 | 2 | 0 | 0 | 0 | 44.01 | | | | | | | | | | | | | | | | | | | | | | | | | | | | | | | |
| | Fe(III) | $Fe^{3+}$ | $Fe^{3+}$ | 3 | 0 | 0 | 0 | 0 | 0 | 0 | 1 | 55.85 | | | | | | | | | | | | | | | | | 1 | | | | | | | | | | | | | | |
| | Carbon monoxide | CO | CO | 0 | 1 | 0 | 0 | 1 | 0 | 0 | 0 | 28.01 | | | | | | | | | | | | | | | | | | | | | | | | | | | | | | | |
| Redox agent | Nitrate | $NO_3^-$ | $NO_3^-$ | -1 | 0 | 0 | 1 | 3 | 0 | 0 | 0 | 62.00 | | | | | | | | 1 | | | | | | | | | | | | | | | | | | | | | | | |
| Methylating agent | Dichloromethane | $C_2H_2Cl_2$ | $CH_2Cl_2$ | 0 | 2 | 2 | 0 | 0 | 0 | 2 | 0 | 96.94 | | | | | | | | | | | | | | | | 1 | | | | | | | | | | | | | | | |
| Ethylating agent | Chloroethane | $C_2H_5Cl$ | $CH_3CH_2Cl$ | 0 | 2 | 5 | 0 | 0 | 0 | 1 | 0 | 64.51 | | | | | | | | | | | | | | | | 2 | | | | | | | | | | | | | | | |
| | Oxalate | $C_2O_4^{(2-)}$ | $(COO^-)_2$ | -2 | 2 | 0 | 0 | 4 | 0 | 0 | 0 | 88.02 | | 2 | | | | | | | | | | | | | | | | | | | | | | | | | | | | | |
| | Formate | $CHO_2^-$ | $HCOO^-$ | -1 | 1 | 1 | 0 | 2 | 0 | 0 | 0 | 45.02 | | 1 | | | | | | | | | | | | | | | | | | | | | | | | | | | | | |
| | Acetate | $C_2H_3O_2^-$ | $CH_3COO^-$ | -1 | 2 | 3 | 0 | 2 | 0 | 0 | 0 | 59.04 | | 1 | | | | | | | | | | | | | | | | | | | | | | | | | | | | | |
| | Propionate | $C_3H_5O_2^-$ | $CH_3CH_2COO^-$ | -1 | 3 | 5 | 0 | 2 | 0 | 0 | 0 | 73.07 | | 1 | | | | | | | | | | | | | | | | | | | | | | | | | | | | | |
| | Butyrate | $C_4H_7O_2^-$ | $CH_3CH_2CH_2COO^-$ | -1 | 4 | 7 | 0 | 2 | 0 | 0 | 0 | 87.10 | | 1 | | | | | | | | | | | | | | | | | | | | | | | | | | | | | |
| | Isobutyrate | $C_4H_7O_2^-$ | $(CH_3)_2CHCOO^-$ | -1 | 4 | 7 | 0 | 2 | 0 | 0 | 0 | 87.10 | | 1 | | | | | | | | | | | | 1 | | | | | | | | | | | | | | | | | |
| | Valerate | $C_5H_9O_2^-$ | $CH_3(CH_2)_3COO^-$ | -1 | 5 | 9 | 0 | 2 | 0 | 0 | 0 | 101.12 | | 1 | | | | | | | | | | | | | | | | | | | | | | | | | | | | | |
| | Caproate | $C_6H_{11}O_2^-$ | $CH_3(CH_2)_4COO$- | -1 | 6 | 11 | 0 | 2 | 0 | 0 | 0 | 115.15 | | 1 | | | | | | | | | | | | | | | | | | | | | | | | | | | | | |
| | Haptonoate/Enanthic acid | $C_7H_{13}O_2^-$ | $CH_3(CH_2)_5COO^-$ | -1 | 7 | 13 | 0 | 2 | 0 | 0 | 0 | 129.18 | | 1 | | | | | | | | | | | | | | | | | | | | | | | | | | | | | |
| | Isoheptanoate | $C_7H_{13}O_2^-$ | $CH_3(CH_2)_3CH(COO^-)CH_3$ | -1 | 7 | 13 | 0 | 2 | 0 | 0 | 0 | 129.18 | | 1 | | | | | | | | | | | | | | | | | | | | | | | | | | | | | |
| | Octonoate/Caprylate | $C_8H_{15}O_2^-$ | $CH_3(CH_2)_6COO^-$ | -1 | 8 | 15 | 0 | 2 | 0 | 0 | 0 | 143.20 | | 1 | | | | | | | | | | | | | | | | | | | | | | | | | | | | | |
| | Nonanoate/Pelargonate | $C_9H_{17}O_2^-$ | $CH_3(CH_2)_7COO^-$ | -1 | 9 | 17 | 0 | 2 | 0 | 0 | 0 | 157.23 | | 1 | | | | | | | | | | | | | | | | | | | | | | | | | | | | | |
| | Decanoate/Caprate | $C_{10}H_{19}O_2^-$ | $CH_3(CH_2)_8COO^-$ | -1 | 10 | 19 | 0 | 2 | 0 | 0 | 0 | 171.26 | | 1 | | | | | | | | | | | | | | | | | | | | | | | | | | | | | |
| | Dodecanoate/Laurate | $C_{12}H_{23}O_2^-$ | $CH_3(CH_2)_{10}COO^-$ | -1 | 12 | 23 | 0 | 2 | 0 | 0 | 0 | 199.31 | | 1 | | | | | | | | | | | | | | | | | | | | | | | | | | | | | |
| | Tetradecanoate/Myristate | $C_{14}H_{27}O_2^-$ | $CH_3(CH_2)_{12}COO$ | -1 | 14 | 27 | 0 | 2 | 0 | 0 | 0 | 227.36 | | 1 | | | | | | | | | | | | | | | | | | | | | | | | | | | | | |
| | Palmitate | $C_{16}H_{31}O_2^-$ | $CH_3(CH_2)_{14}COO^-$ | -1 | 16 | 31 | 0 | 2 | 0 | 0 | 0 | 255.42 | | 1 | | | | | | | | | | | | | | | | | | | | | | | | | | | | | |
| | Octadecanoate/Stearate | $C_{18}H_{35}O_2^-$ | $CH_3(CH_2)_{16}COO^-$ | -1 | 18 | 35 | 0 | 2 | 0 | 0 | 0 | 283.47 | | 1 | | | | | | | | | | | | | | | | | | | | | | | | | | | | | |
| | Linoleate | $C_{18}H_{31}O_2^-$ | $COO^-(CH_2)CH=CHCH_2CH=CH(CH_2)_4CH_3$ | -1 | 18 | 31 | 0 | 2 | 0 | 0 | 0 | 279.44 | | 1 | | | | | | | | | | | | | | | | | | 2 | | | | | | | | | | | |
| | Crotonate/2-butenoate/b-methylacrylate | $C_4H_5O_2^-$ | $CH_3HC=CHCOO^-$ | -1 | 4 | 5 | 0 | 2 | 0 | 0 | 0 | 85.08 | | 1 | | | | | | | | | | | | | | | | | | 1 | | | | | | | | | | | |
| | Cinnamate | $C_9H_7O_2^-$ | $C_6H_5CH=CHCOO^-$ | -1 | 9 | 7 | 0 | 2 | 0 | 0 | 0 | 147.15 | | 1 | | | | | | | | | | | | | | | | | | 1 | | 1 | | | | | | | | | |
| | Pyruvate | $C_3H_3O_3^-$ | $CH_3COCOO^-$ | -1 | 3 | 3 | 0 | 3 | 0 | 0 | 0 | 87.05 | | 1 | | | | | | | | | 1 | | | | | | | | | | | | | | | | | | | | |
| | Lactate | $C_3H_5O_3^-$ | $CH_3CH(OH)COO^-$ | -1 | 3 | 5 | 0 | 3 | 0 | 0 | 0 | 89.07 | | 1 | 1 | | | | | | | | | | | | | | | | | | | | | | | | | | | | |
| | Fumarate | $C_4H_2O_4^-$ | $HOOC(CH)=(CH)COO^-$ | -1 | 4 | 2 | 0 | 4 | 0 | 0 | 0 | 114.06 | 1 | 1 | | | | | | | | | | | | | | | | | | 1 | | | | | | | | | | | |
| | Glutarate | $C_5H_7O_4^-$ | $COOH(CH_2)_3COO^-$ | -1 | 5 | 7 | 0 | 4 | 0 | 0 | 0 | 131.11 | 1 | 1 | | | | | | | | | | | | | | | | | | | | | | | | | | | | | |
| | Gluconate | $C_6H_{11}O_7^-$ | $(OH)CH_2(CHOH)_4COO^-$ | -1 | 6 | 11 | 0 | 7 | 0 | 0 | 0 | 195.15 | | 1 | 5 | | | | | | | | | | | | | | | | | | | | | | | | | | | | |
| | Succinate/Butanedioate | $C_4H_4O_4^{2-}$ | $(COO^-)_2(CH_2)_2$ | -2 | 4 | 4 | 0 | 4 | 0 | 0 | 0 | 116.07 | | 2 | | | | | | | | | | | | | | | | | | | | | | | | | | | | | |
| | Malate | $C_4H_4O_5^{2-}$ | $(COO^-)CH_2CH(OH)COO^-$ | -2 | 4 | 4 | 0 | 5 | 0 | 0 | 0 | 132.07 | | 2 | 1 | | | | | | | | | | | | | | | | | | | | | | | | | | | | |
| | Tartrate | $C_4H_4O_6^{2-}$ | $^-OOC(CH)_2COO^-$ | -2 | 4 | 4 | 0 | 6 | 0 | 0 | 0 | 148.07 | | 2 | | | | | | | | | | | | | | | | | | | | | | | | | | | | | |
| | Adipate | $C_6H_{10}O_4^{2-}$ | $(CH_2)_4(COO^-)_2$ | -2 | 6 | 10 | 0 | 4 | 0 | 0 | 0 | 146.14 | | 2 | | | | | | | | | | | | | | | | | | | | | | | | | | | | | |
| Carboxylate | Citrate | $C_6H_5O_7^{2-}$ | $COO^-CH_2C(OH)(COO^-)CH_2COO^-$ | -2 | 6 | 5 | 0 | 7 | 0 | 0 | 0 | 189.10 | | 3 | 1 | | | | | | | | | | | | | | | | | | | | | | | | | | | | |
| | Glyoxylate | $C_2HO_3^-$ | $COH(CH)OO^-$ | -1 | 2 | 1 | 0 | 3 | 0 | 0 | 0 | 73.03 | | 1 | 1 | | | | | | | | | | | | | | | | | | | | | | | | | | | | |
| | Glycolate | $C_2H_3O_3^-$ | $HO(CH2)COO^-$ | -1 | 2 | 3 | 0 | 3 | 0 | 0 | 0 | 75.04 | | 1 | 1 | | | | | | | | | | | | | | | | | | | | | | | | | | | | |
| | Glycerate | $C_3H_5O_4^-$ | $CH_2(OH)CH(OH)COO^-$ | -1 | 3 | 5 | 0 | 4 | 0 | 0 | 0 | 105.07 | 1 | 1 | 2 | | | | | | | | | | | | | | | | | | | | | | | | | | | | |
| | Hydroxybutyrate | $C_4H_7O_3^-$ | $CH_3CH_2CH(OH)COO^-$ | -1 | 4 | 7 | 0 | 3 | 0 | 0 | 0 | 103.10 | | 1 | 1 | | | | | | | | | | | | | | | | | | | | | | | | | | | | |
| | 2-Hydroxybutyrate | $C_4H_7O_3^-$ | $CH_3CH_2CH(OH)COO^-$ | -1 | 4 | 7 | 0 | 3 | 0 | 0 | 0 | 103.10 | | 1 | 1 | | | | | | | | | | | | | | | | | | | | | | | | | | | | |
| | 3-Hydroxybutyrate | $C_4H_7O_3^-$ | $CH_3CH(OH)CH_2COO^-$ | -1 | 4 | 7 | 0 | 3 | 0 | 0 | 0 | 103.10 | | 1 | 1 | | | | | | | | | | | | | | | | | | | | | | | | | | | | |
| | 4-Hydroxybutyrate | $C_4H_7O_3^-$ | $HO(CH_2)_3COO^-$ | -1 | 4 | 7 | 0 | 3 | 0 | 0 | 0 | 103.10 | | 1 | 1 | | | | | | | | | | | | | | | | | | | | | | | | | | | | |
| Hydroxy carboxylate | Ferulate/Ethyl 4-hydroxy-3-methoxycinnamate | $C_{10}H_{10}O_4$ | $HOC_6H_3CH=CHCOO^-$ | 0 | 10 | 10 | 0 | 4 | 0 | 0 | 0 | 194.18 | | 1 | 1 | | | | | | | | | | | | | | | | | 1 | | 1 | | | | | | | | | |
| | Methylpropionate | $C_4H_7O_2^-$ | $CH_3CH_2(COO^-)CH_3$ | -1 | 4 | 7 | 0 | 2 | 0 | 0 | 0 | 87.10 | | 1 | | | | | | | | | | | | 1 | | | | | | | | | | | | | | | | | |
| | 3-methyl-butyrate | $C_5H_9O_2^-$ | $CH_3(CH_2)_2(COO^-)CH_3$ | -1 | 5 | 9 | 0 | 2 | 0 | 0 | 0 | 101.12 | | 1 | | | | | | | | | | | | 1 | | | | | | | | | | | | | | | | | |
| Methyl carboxylate | 4-Methyl-valerate/4-methylpentanoate | $C_6H_{11}O_2^-$ | $CH_3(CH_2)_3(COO^-)CH_3$ | -1 | 6 | 11 | 0 | 2 | 0 | 0 | 0 | 115.15 | | 1 | | | | | | | | | | | | 1 | | | | | | | | | | | | | | | | | |
| | Syringate/4-hydroxy-3,5-dimethoxybenzoate/3,5-dimethoxy-4-hydroxybenzoate | $C_9H_9O_5$ | $COO^-C=CHC(OCH_3)=C(OH)C(OCH_3)=CH$ | 0 | 9 | 9 | 0 | 5 | 0 | 0 | 0 | 197.16 | | 1 | 1 | | | | | | | | | | | | 2 | | | | 3 | | 1 | | | | | | | | | | |
| | 3-methoxycinnamate | $C_{10}H_9O_3^-$ | $(CH_3)C_6H_4CH=CHCOO^-$ | -1 | 10 | 9 | 0 | 3 | 0 | 0 | 0 | 177.18 | | 1 | | | | | | | | | | | | | 1 | | | | 1 | | 1 | | | | | | | | | | |
| | 4-methyoxycinnamate/p-methoxycinnamate | $C_{10}H_9O_3^-$ | NA | -1 | 10 | 9 | 0 | 3 | 0 | 0 | 0 | 177.18 | | 1 | | | | | | | | | | | | | 1 | | | | 1 | | 1 | | | | | | | | | | |
| | 3,4-dimethoxycinnamate | $C_{11}H_{11}O_4^-$ | NA | -1 | 11 | 11 | 0 | 4 | 0 | 0 | 0 | 207.20 | | 1 | | | | | | | | | | | | | 1 | | | | 1 | | 1 | | | | | | | | | | |
| Methyloxy carboxylate | 3,4,5-trimethoxycinnamate | $C_{12}H_{13}O_5^-$ | NA | -1 | 12 | 13 | 0 | 5 | 0 | 0 | 0 | 237.23 | | 1 | | | | | | | | | | | | | 1 | | | | 1 | | 1 | | | | | | | | | | |
| | Vanillate/4-hydroxy-3-methoxybenzoate | $C_8H_8O_4$ | $COO^-C=CHC(OCH_3)=C(OH)HC=CH$ | 0 | 8 | 8 | 0 | 4 | 0 | 0 | 0 | 168.15 | | 1 | 1 | | | | | | | | | | | | 1 | | | | 3 | | 1 | | | | | | | | | | |
| Hydoxyl methyloxy carboxylate | Sinapate/3,5-dimethoxy-4-hydroxycinnamate | $C_{11}H_{12}O_5^-$ | $COO^-HC=CH(C=CHC(OCH_3)=COHC(OCH_3)=CH)$ | -1 | 11 | 12 | 0 | 5 | 0 | 0 | 0 | 224.21 | | 1 | 1 | | | | | | | | | | | | 2 | | | | 4 | | 1 | | | | | | | | | | |
| | Phenyl acetate | $C_8H_8O_2$ | $CH3COO(C_6H_5)$ | 0 | 8 | 8 | 0 | 2 | 0 | 0 | 0 | 136.15 | | 1 | | | | | | | | | | | 1 | 1 | | | | | | | 1 | | | | | | | | | | |
| Phenyl carboxylate | 3-phenylpropionate/Hydrocinnamic acid | $C_9H_{10}O_2$ | $COOH(CH_2)_2(C_6H_5)$ | 0 | 9 | 10 | 0 | 2 | 0 | 0 | 0 | 150.18 | 1 | | | | | | | | | | | | | | | | | | | | 1 | | | | | | | | | | |
| | 4-amino-butyrate/γ-aminobutyrate | $C_4H_8NO_2^-$ | $NH_2(CH_2)_3COO^-$ | -1 | 4 | 8 | 1 | 2 | 0 | 0 | 0 | 102.11 | | 1 | | 1 | | | | | | | | | | | | | | | | | | | | | | | | | | | |
| | Aspartate | $C_4H_6NO_4^-$ | $COOHCH2CH(NH2)COO^-$ | -1 | 4 | 6 | 1 | 4 | 0 | 0 | 0 | 132.09 | 1 | 1 | | 1 | | | | | | | | | | | | | | | | | | | | | | | | | | | |
| | Aminovalerate/Aminopentanoate | $C_5H_{11}NO_2$ | $NH_2(CH_2)_4COO^-$ | 0 | 5 | 11 | 1 | 2 | 0 | 0 | 0 | 117.15 | | 1 | | 1 | | | | | | | | | | | | | | | | | | | | | | | | | | | |
| | Glutamate | $C_5H_8NO_4^-$ | $COO^-(CH_2)_2CH(NH_3)COO^-$ | -1 | 5 | 8 | 1 | 4 | 0 | 0 | 0 | 146.12 | | 2 | | | | | 1 | | | | | | | | | | | | | | | | | | | | | | | | |
| Aminocarboxylate | Benzoylamino acetate/Hippurate | $C_9H_{10}NO_2^-$ | $(C_6H_5)CH_2NHCH_2COO^-$ | -1 | 9 | 10 | 1 | 2 | 0 | 0 | 0 | 164.18 | | 1 | | 1 | | | | | | | | | | | | | | | | 1 | | | | | | | | | | | |
| | Methanol | $CH_4O$ | $CH_3OH$ | 0 | 1 | 4 | 0 | 1 | 0 | 0 | 0 | 32.04 | | | 1 | | | | | | | | | | | | | | | | | | | | | | | | | | | | |
| | Ethanol | $C_2H_6O$ | $CH_3CH_2OH$ | 0 | 2 | 6 | 0 | 1 | 0 | 0 | 0 | 46.07 | | | 1 | | | | | | | | | | | | | | | | | | | | | | | | | | | | |
| | Ethylene glycol | $C_2H_6O_2$ | $(CH_2OH)_2$ | 0 | 2 | 6 | 0 | 2 | 0 | 0 | 0 | 62.07 | | | 2 | | | | | | | | | | | | | | | | | | | | | | | | | | | | |
| | 1-propanol | $C_3H_8O$ | $CH_3(CH_2)_2OH$ | 0 | 3 | 8 | 0 | 1 | 0 | 0 | 0 | 60.09 | | | 1 | | | | | | | | | | | | | | | | | | | | | | | | | | | | |
| | 2-propanol/Isopropanol | $C_3H_8O$ | $(CH_3)_2CHOH$ | 0 | 3 | 8 | 0 | 1 | 0 | 0 | 0 | 60.09 | | | 1 | | | | | | | | | | | 1 | | | | | | | | | | | | | | | | | |
| | 1,2-propanediol/Propylene glycol | $C_3H_8O_2$ | $CH_3(CH_2OH)_2$ | 0 | 3 | 8 | 0 | 2 | 0 | 0 | 0 | 76.09 | | | 2 | | | | | | | | | | | | | | | | | | | | | | | | | | | | |
| | 1,3-propanediol | $C_3H_8O_2$ | $CH_2(CH_2OH)_2$ | 0 | 3 | 8 | 0 | 2 | 0 | 0 | 0 | 76.09 | | | 2 | | | | | | | | | | | | | | | | | | | | | | | | | | | | |
| | 1-butanol | $C_4H_{10}O$ | $CH_3(CH_2)_3OH$ | 0 | 4 | 10 | 0 | 1 | 0 | 0 | 0 | 74.12 | | | 1 | | | | | | | | | | | | | | | | | | | | | | | | | | | | |
| | 2-butanol/isobutanol | $C_4H_{10}O$ | $CH_3CH_2CH(OH)CH_3$ | 0 | 4 | 10 | 0 | 1 | 0 | 0 | 0 | 74.12 | | | 1 | | | | | | | | | | | | | | | | | | | | | | | | | | | | |
| | 2-pentanol | $C_5H_{12}O$ | $CH_3CH(OH)(CH_2)_2CH_3$ | 0 | 5 | 12 | 0 | 1 | 0 | 0 | 0 | 88.15 | | | 1 | | | | | | | | | | | | | | | | | | | | | | | | | | | | |
| | 3-pentanol | $C_5H_{12}O$ | $CH_3CH_2CH(OH)CH_2CH_3$ | 0 | 5 | 12 | 0 | 1 | 0 | 0 | 0 | 88.15 | | | 1 | | | | | | | | | | | | | | | | | | | | | | | | | | | | |
| | Cyclopentanol | $C_5H_{10}O$ | $(CH_2)_4CH(OH)$ | 0 | 5 | 10 | 0 | 1 | 0 | 0 | 0 | 86.13 | | | 1 | | | | | | | | | | | | | | | | | | | | | | | | | | | | |
| Alcohol | Adonitol | $C_5H_{12}O_5$ | $CH_2(OH)CH(OH)_3CH_2(OH)$ | 0 | 5 | 12 | 0 | 5 | 0 | 0 | 0 | 152.14 | | | 5 | | | | | | | | | | | | | | | | | | | | | | | | | | | | |
| Amino alcohol | Ethanolamine | $C_2H_7NO$ | $HOCH_2CH_2NH_2$ | 0 | 2 | 7 | 1 | 1 | 0 | 0 | 0 | 61.08 | | | 1 | 1 | | | | | | | | | | | | | | | | | | | | | | | | | | | |
| | Glycerol | $C_3H_8O_3$ | $(CH_2OH)_2CHOH$ | 0 | 3 | 8 | 0 | 3 | 0 | 0 | 0 | 92.09 | | | 3 | | | | | | | | | | | | | | | | | | | | | | | | | | | | |
| | Erythritol | $C_4H_{10}O_4$ | $(CH_2OH)_2(HCOH)_2$ | 0 | 4 | 10 | 0 | 4 | 0 | 0 | 0 | 122.12 | | | 4 | | | | | | | | | | | | | | | | | | | | | | | | | | | | |
| | Arabitol | $C_5H_{12}O_5$ | $(HCOH)_3(CH_2OH)_2$ | 0 | 5 | 12 | 0 | 5 | 0 | 0 | 0 | 152.14 | | | 5 | | | | | | | | | | | | | | | | | | | | | | | | | | | | |
| | Inositol/Cyclohexanehexol | $C_6H_{12}O_6$ | $(CHOH)_6$ | 0 | 6 | 12 | 0 | 6 | 0 | 0 | 0 | 180.15 | | | 6 | | | | | | | | | | | | | | | | | | | | | | | | | | | | |
| | Galactitol/Dulcitol | $C_6H_{14}O_6$ | $(HO)CH_2(HCOH)_4CH_2(OH)$ | 0 | 6 | 14 | 0 | 6 | 0 | 0 | 0 | 182.17 | | | 6 | | | | | | | | | | | | | | | | | | | | | | | | | | | | |
| | Mannitol | $C_6H_{14}O_6$ | $(CH_2OH)_2(CHOH)_4$ | 0 | 6 | 14 | 0 | 6 | 0 | 0 | 0 | 182.17 | | | 6 | | | | | | | | | | | | | | | | | | | | | | | | | | | | |
| Sugar alcohol | Sorbitol/Glucitol | $C_6H_{14}O_6$ | $(CH_2OH)_2(CHOH)_4$ | 0 | 6 | 14 | 0 | 6 | 0 | 0 | 0 | 182.17 | | | 6 | | | | | | | | | | | | | | | | | | | | | | | | | | | | |
| | Ribose | $C_5H_{10}O_5$ | $(CH_2OH)CH_2(CHOH)_3O$ | 0 | 5 | 10 | 0 | 5 | 0 | 0 | 0 | 150.13 | | | 4 | | | | | | | | | 1 | | | | | | | | | | | | | | | | | | | |
| | Xylose | $C_5H_{10}O_5$ | $(CHOH)_5O$ | 0 | 5 | 10 | 0 | 5 | 0 | 0 | 0 | 150.13 | | | 5 | | | | | | | | | | | | | | | | | | | | | | | | | | | | |
| Pentose | Arabinose | $C_5H_{10}O_5$ | $(HO)CH_2(CHOH)_3COOH$ | 0 | 5 | 10 | 0 | 5 | 0 | 0 | 0 | 150.13 | 1 | | 4 | | | | | | | | | 1 | | | | | | | | | | | | | | | | | | | |
| | Galactose | $C_6H_{12}O_6$ | $CH_2(OH)(HCOH)_4COOH$ | 0 | 6 | 12 | 0 | 6 | 0 | 0 | 0 | 180.15 | 1 | | 5 | | | | | | | | | 1 | | | | | | | | | | | | | | | | | | | |
| | Glucose | $C_6H_{12}O_6$ | $H(C=O)(CHOH)_5H$ | 0 | 6 | 12 | 0 | 6 | 0 | 0 | 0 | 180.15 | | | 5 | | | | | | | | | 1 | | | | | | | | | | | | | | | | | | | |
| | Mannose | $C_6H_{12}O_6$ | $CH_2OH(CHOH)_5O$ | 0 | 6 | 12 | 0 | 6 | 0 | 0 | 0 | 180.15 | | | 6 | | | | | | | | | 1 | | | | | | | | | | | | | | | | | | | |
| | Tagatose | $C_6H_{12}O_6$ | $CH_2(OH)(CHOH)_3C=OCH_2OH$ | 0 | 6 | 12 | 0 | 6 | 0 | 0 | 0 | 180.15 | | | 5 | | | | | | | | 1 | | | | | | | | | | | | | | | | | | | | |
| | Sorbose | $C_6H_{12}O_6$ | $CH_2(OH)(CHOH)_4CH_2O$ | 0 | 6 | 12 | 0 | 6 | 0 | 0 | 0 | 180.15 | | | 5 | | | | | | | | 1 | | | | | | | | | | | | | | | | | | | | |
| Hexose | Fructose | $C_6H_{12}O_6$ | NA | 0 | 6 | 12 | 0 | 6 | 0 | 0 | 0 | 180.15 | | | 1 | | | | | | | | 1 | | | | | | | | | | | | | | | | | | | | |
| | Methylmannoside/methyl mannopyranoside | $C_7H_{14}O_6$ | $(HCOH)_2CH_2(OCH_3)COH(O)CHO(CH_3)$ | 0 | 7 | 14 | 0 | 6 | 0 | 0 | 0 | 194.18 | | | 5 | | | | | | | | | | | 1 | 1 | | | | | | | | | | | 1 | | | | | |
| | Methylglucoside | $C_7H_{14}O_6$ | $(HCOH)_3(COHCH_3)O(COHCH_3)$ | 0 | 7 | 14 | 0 | 6 | 0 | 0 | 0 | 194.18 | | | 5 | | | | | | | | | | | 1 | | | | | | | | | | | | 1 | | | | | |
| Methyl hexose | Methylglucopyranoside | $C_7H_{14}O_6$ | $(HCOH)_3CH_2(CH_2OH)O(HCOCH_3)$ | 0 | 7 | 14 | 0 | 6 | 0 | 0 | 0 | 194.18 | | | 4 | | | | | | | | | | | 1 | | | | | | | | | | | | 1 | | | | | |
| | Glucopyranoside | $C_6H_{11}O_6$ | $(HCOH)_3CH_2(H_2COH)OCO$ | 0 | 6 | 11 | 0 | 6 | 0 | 0 | 0 | 179.15 | | | 5 | | | | | | | | | | | | | | | | | | | | | | | 1 | | | | | |
| Methyloxy hexose | Mannopyranoside | $C_6H_{11}O_6$ | $(HCOH)_4COCH_3$ | 0 | 6 | 11 | 0 | 6 | 0 | 0 | 0 | 179.15 | | | 4 | | | | | | | | | | | | | | | | | | | | | | | 1 | | | | | |
| Amino hexose | Glucosamine | $C_6H_{13}NO_5$ | $CH_2(OH)CHOCH(OH)CH(NH_2)(HCOH)_2$ | 0 | 6 | 13 | 1 | 5 | 0 | 0 | 0 | 179.17 | | | 5 | 1 | | | | | | | | | | | | | | | | | | | | | | 1 | | | | | |
| | Acetylglucosamine | $C_8H_{15}NO_6$ | $(HCOH)_2CH(CH_2OH)OCH_2(OH)CH_2(NHCOCH_3)$ | 0 | 8 | 15 | 1 | 6 | 0 | 0 | 0 | 221.21 | | | 4 | | 1 | | | | | 1 | | | | | | | | | | | | | | | | 1 | | | | | |
| Aminoacetylhexose | Acetylgalactosamine | $C_8H_{15}NO_6$ | $(HCOH)_2CH(CH_2OH)OCH_2(OH)CH(NHCOCH_3)$ | 0 | 8 | 15 | 1 | 6 | 0 | 0 | 0 | 221.21 | | | 4 | | 1 | | | | | 1 | | | | | | | | | | | | | | | | 1 | | | | | |
| | Rhamnose/6-deoxy-L-mannopyranose | $C_6H_{12}O_5$ | $(CH_3)CH_2(CHOH)_4O$ | 0 | 6 | 12 | 0 | 5 | 0 | 0 | 0 | 164.16 | | | 4 | | | | | | | | | | | | | | | | | | | | | | | | | | | | |
| Deoxy hexose | Fucose | $C_6H_{12}O_5$ | $CH_3(HCOH)_4COOH$ | 0 | 6 | 12 | 0 | 5 | 0 | 0 | 0 | 164.16 | 1 | | 4 | | | | | | | | | | | | | | | | | | | | | | | | 1 | | | | |
| | Lactose | $C_{12}H_{22}O_{11}$ | NA | 0 | 12 | 22 | 0 | 11 | 0 | 0 | 0 | 342.29 | | | 1 | | | | | | | | | | | | | | | | | | | | | | | | | | | | |
| | Melibiose | $C_{12}H_{22}O_{11}$ | NA | 0 | 12 | 22 | 0 | 11 | 0 | 0 | 0 | 342.29 | | | 1 | | | | | | | | | | | | | | | | | | | | | | | | | | | | |
| | Maltose | $C_{12}H_{22}O_{11}$ | NA | 0 | 12 | 22 | 0 | 11 | 0 | 0 | 0 | 342.29 | | | 1 | | | | | | | | | | | | | | | | | | | | | | | | | | | | |
| | Sucrose/Saccharose | $C_{12}H_{22}O_{11}$ | Glc(α-1,2-β)Fruf | 0 | 12 | 22 | 0 | 11 | 0 | 0 | 0 | 342.29 | | | 1 | | | | | | | | | | | | | | | | | | | | | | | | | | | | |
| | Trehalose | $C_{12}H_{22}O_{11}$ | Glc(α-1,1-α)Glc | 0 | 12 | 22 | 0 | 11 | 0 | 0 | 0 | 342.29 | | | 1 | | | | | | | | | | | | | | | | | | | | | | | | | | | | |
| | Turanose | $C_{12}H_{22}O_{11}$ | Glc(α-1,3)Fruf | 0 | 12 | 22 | 0 | 11 | 0 | 0 | 0 | 342.29 | 1 | | 1 | | | | | | | | | | | | | | | | | | | | | | | | | | | | |
| | Cellobiose | $C_{12}H_{22}O_{11}$ | $(C_6H_7(OH)_4O)_2O$ | 0 | 12 | 22 | 0 | 11 | 0 | 0 | 0 | 342.29 | | | 8 | | | | | | | | | | | | | | | | | | | | | | | | | | | | |
| Disaccharide | Gentiobiose | $C_{12}H_{22}O_{11}$ | NA | 0 | 12 | 22 | 0 | 11 | 0 | 0 | 0 | 342.29 | | | 1 | | | | | | | | | | | | | | | | | | | | | | | | | | | | |
| | Maltotriose | $C_{18}H_{32}O_{16}$ | NA | 0 | 18 | 32 | 0 | 16 | 0 | 0 | 0 | 504.43 | | | 1 | | | | | | | | | | | | | | | | | | | | | | | | | | | | |
| | Melezitose | $C_{18}H_{32}O_{16}$ | NA | 0 | 18 | 32 | 0 | 16 | 0 | 0 | 0 | 504.43 | | | 1 | | | | | | | | | | | | | | | | | | | | | | | | | | | | |
| Trisaccharide | Raffinose | $C_{18}H_{32}O_{16}$ | Gal(α-1,6)Glc(α-1,2β)Fruf | 0 | 18 | 32 | 0 | 16 | 0 | 0 | 0 | 504.43 | | | 1 | | | | | | | | | | | | | | | | | | | | | | | | | | | | |
| | Hemicellulose/polyose | $(C_5H_8O_4)_n$ | NA | 0 | 5 | 8 | 0 | 4 | 0 | 0 | 0 | 132.11 | | | | | | | | | | | | | | | | | | | | | | | | | | | | | | | |
| | Xylan | $C_5H_{10}O_5$ | Xyl(β-2,2-β)Xyl | 0 | 5 | 10 | 0 | 5 | 0 | 0 | 0 | 150.13 | | | 1 | | | | | | | | | | 1 | | 1 | | | | | | | | | | | | | | | | |
| | Amidon/starch | $C_6H_7N_3O$ | $C_5H_4N\text{-}CONH(NH_2)$ | 0 | 6 | 7 | 3 | 1 | 0 | 0 | 0 | 137.14 | | | 1 | | | | | | | | | | | | | | | | | | | | | | | | | | | | |
| | Cellulose | $(C_6H_{10}O_5)_n$ | NA | 0 | 6 | 10 | 0 | 5 | 0 | 0 | 0 | 162.14 | | | | | | | | | | | | | | | | | | | | | | | | | | | | | | | |
| | Lichenin | $(C_6H_{10}O_5)_n$ | NA | 0 | 6 | 10 | 0 | 5 | 0 | 0 | 0 | 162.14 | | | 1 | | | | | | | | | | | | | | | | | | | | | | | | | | | | |
| | Starch | $(C_6H_{10}O_5)_n$ | Glc(α-1,4)Glc | 0 | 6 | 10 | 0 | 5 | 0 | 0 | 0 | 162.14 | | | 1 | | | | | | | | | | | | 1 | | | | | | | | | | | | | | | | |
| | Pectin | $C_6H_{10}O_7$ | $(HCOH)_4OCH(COOH)$ | 0 | 6 | 10 | 0 | 7 | 0 | 0 | 0 | 194.14 | 1 | | 4 | | | | | | | | | | | | | | | | | | | | | | | | | | | | |
| | Arabinoxylan | $(C_{10}H_{16}O_8)_n$ | NA | 0 | 10 | 16 | 0 | 8 | 0 | 0 | 0 | 264.23 | | | | | | | | | | | | | | | | | | | | | | | | | | | | | | | |
| | Glucan | $C_{18}H_{32}O_{16}$ | NA | 0 | 18 | 32 | 0 | 16 | 0 | 0 | 0 | 504.43 | | | 1 | | | | | | | | | | | | | | | | | | | | | | | | | | | | |
| | Laminarin | $C_{18}H_{32}O_{16}$ | NA | 0 | 18 | 32 | 0 | 16 | 0 | 0 | 0 | 504.43 | | | 1 | | | | | | | | | | | | | | | | | | | | | | | | | | | | |
| | Pullulan | $C_{18}H_{32}O_{16}$ | NA | 0 | 18 | 32 | 0 | 16 | 0 | 0 | 0 | 504.43 | | | 1 | | | | | | | | | | | | | | | | | | | | | | | | | | | | |
| | Arabinogalactan/Galactoarabinan | $C_{20}H_{36}O_{14}$ | NA | 0 | 20 | 10 | 0 | 14 | 0 | 0 | 0 | 474.28 | | | 1 | | | | | | | | | | | | 1 | | | | | | | | | | | 1 | | | | | |
| | Glycogen | $C_{24}H_{42}O_{21}$ | NA | 0 | 24 | 42 | 0 | 21 | 0 | 0 | 0 | 666.57 | | | 1 | | | | | | | | | | | | | | | | | | | | | | | | | | | | |
| | Amylopectin | $C_{30}H_{52}O_{26}$ | NA | 0 | 30 | 52 | 0 | 26 | 0 | 0 | 0 | 828.71 | | | 1 | | | | | | | | | | | | | | | | | | | | | | | | | | | | |
| Polysaccharide | Maltodextrin | $C_{6n}H_{10n+2}O_{5n+1}$ | NA | 0 | NA | NA | 0 | NA | 0 | 0 | 0 | NA | | | 1 | | | | | | | | | | | | | | | | | | | | | | | | | | | | |
| Aminopolysaccharide | Chitin | $(C_8H_{13}O_5N)_n$ | NA | 0 | 8 | 13 | 1 | 5 | 0 | 0 | 0 | 203.19 | | | | | 1 | | | | | 1 | | | | 1 | | | | | | | | | | | | | | | 1 | | |
| | Cyclodextrin/Cycloheptaamylose/Cyclomaltoheptaose/Cycloheptaglucan | $(C_6H_{10}O_5)_7$ | $(C_6H_{10}O_5)_7$ | 0 | 6 | 10 | 0 | 5 | 0 | 0 | 0 | 162.14 | | | 5 | | | | | | | | | | | | | | | | | | | | | | | | | | | | |
| | Dextrin | $C_{18}H_{32}O_{16}$ | NA | 0 | 18 | 32 | 0 | 16 | 0 | 0 | 0 | 504.43 | | | 1 | | | | | | | | | | | | | | | | | | | | | | | | | | | | |
| | Galactooligosaccharides | $C_{18}H_{32}O_{16}$ | NA | 0 | 18 | 32 | 0 | 16 | 0 | 0 | 0 | 504.43 | | | 1 | | | | | | | | | | | | | | | | | | | | | | | | | | | | |
| | Fructooligosaccharides | $C_{72}H_{122}O_{61}$ | $Fruf(2\text{-}1)_{11}\text{-}Glc$ | 0 | 72 | 122 | 0 | 61 | 0 | 0 | 0 | 1963.69 | | | 1 | | | | | | | | | | | | | | | | | | | | | | | | | | | | |
| Oligosaccharide | Inulin | $C_{6n}H_{10n+2}O_{5n+1}$ | NA | 0 | NA | NA | 0 | NA | 0 | 0 | 0 | NA | | | 1 | | | | | | | | | | | | | | | | | | | | | | | | 1 | | | | |

[Supplementary Table 1.1 continued]

| Functional Category | Name | Molecular formula | Condensed Formula | Ionic Charge | Elemental composition | | | | | | | Molecular weight | R-COOH, carboxylic acid | $R-COO^-$, carboxylate | R-OH, hydroxyl | $R-NH_2$, amine | R-NH | R-N-R | $R-NH_3^+$ | $R-NO_3^-$ | Nitrile, CN | R-N-(C=O)-R, amide | R-C=O, ketone | R-HC=O, aldehyde | R-O-C=O, Ester | $R-CH_3$, methyl | $R-O-CH_3$, oxymethyl | R-Cl | $R-Fe^{3+}$ | R-S | C=C | Benzene | Phenyl | Pyridine | Imidazole | Indole | Oleate | pyranose | furanose | pyrrolizidine | Glucosamine | Purine | Pteridine |
|---|---|---|---|---|---|---|---|---|---|---|---|---|---|---|---|---|---|---|---|---|---|---|---|---|---|---|---|---|---|---|---|---|---|---|---|---|---|---|---|---|---|---|---|
| | | | | | C | H | N | O | S | Cl | Fe | | | | | | | | | | | | | | | | | | | | | | | | | | | | | | | | |
| | Arbutin | $C_{12}H_{16}O_7$ | NA | 0 | 12 | 16 | 0 | 7 | 0 | 0 | 0 | 272.25 | | | 1 | | | | | | | | | | | | | | | | | 1 | 1 | | | | | | | | | | |
| Glycoside | Salicin | $C_{13}H_{18}O_7$ | NA | 0 | 13 | 18 | 0 | 7 | 0 | 0 | 0 | 286.28 | | | 1 | | | | | | | | | | | | | | | | | 1 | | | | | | | | | | | |
| Methylglycoside | Methyl a-glycoside | $C_7H_{14}O_6$ | $(HCOH)_3COO(CH_3)CH_2(CH_2OH)$ | 0 | 7 | 14 | 0 | 6 | 0 | 0 | 0 | 194.18 | | | 4 | | | | | | | | | | 1 | 1 | 1 | | | | | | | | | | | 1 | | | | | |
| Ester | Polyoxyethylene sorbitan mono-oleate | $C_{64}H_{124}O_{26}$ | NA | 0 | 64 | 124 | 0 | 26 | 0 | 0 | 0 | 1309.65 | | | | | | | | | | | | | 1 | | | | | | | | | | | | 1 | | | | | | |
| | Glycine | $C_2H_5NO_2$ | $NH_2CH_2COOH$ | 0 | 2 | 5 | 1 | 2 | 0 | 0 | 0 | 75.07 | 1 | | | 1 | | | | | | | | | | | | | | | | | | | | | | | | | | | |
| | Alanine | $C_3H_7NO_2$ | $CH_3CH(NH_2)COOH$ | 0 | 3 | 7 | 1 | 2 | 0 | 0 | 0 | 89.09 | 1 | | | 1 | | | | | | | | | | 1 | | | | | | | | | | | | | | | | | |
| | Serine | $C_3H_7NO_3$ | $HOCH_2CH(NH_2)COOH$ | 0 | 3 | 7 | | 3 | 0 | 0 | 0 | 91.08 | 1 | | 1 | 1 | | | | | | | | | | | | | | | | | | | | | | | | | | | |
| | Cysteine | $C_3H_7NO_2S$ | $HSCH_2CH(NH_2)COOH$ | 0 | 3 | 7 | 1 | 2 | 1 | 0 | 0 | 121.16 | 1 | | | 1 | | | | | | | | | | | | | | 1 | | | | | | | | | | | | | |
| | Threonine | $C_4H_9NO_3$ | $CH_3CH(OH)CH(NH_2)COOH$ | 0 | 4 | 9 | 1 | 3 | 0 | 0 | 0 | 119.12 | 1 | | 1 | 1 | | | | | | | | | | 1 | | | | | | | | | | | | | | | | | |
| | Asparagine | $C_4H_8N_2O_3$ | $H_2NCOCH_2CH(NH_2)COOH$ | 0 | 4 | 8 | 2 | 3 | 0 | 0 | 0 | 132.12 | 1 | | | 2 | | | | | | 1 | | | | | | | | | | | | | | | | | | | | | |
| | Aspartic acid | $C_4H_7NO_4$ | $COOHCH_2CH(NH_2)COOH$ | 0 | 4 | 7 | 1 | 4 | 0 | 0 | 0 | 133.10 | 1 | | | 1 | | | | | | | | | | | | | | | | | | | | | | | | | | | |
| | Proline | $C_5H_9NO_2$ | $NH(CH_2)_3CHCOOH$ | 0 | 5 | 9 | 1 | 2 | 0 | 0 | 0 | 115.13 | 1 | | | 1 | | | | | | | | | | | | | | | | | | | | | | | | | | | |
| | Valine | $C_5H_{11}NO_2$ | $(CH_3)_2CHCH(NH_2)COOH$ | 0 | 5 | 11 | 1 | 2 | 0 | 0 | 0 | 117.15 | 1 | | | 1 | | | | | | | | | | 1 | | | | | | | | | | | | | | | | | |
| | Glutamine | $C_5H_{10}N_2O_3$ | $H_2NCO(CH_2)_2CH(NH_2)COOH$ | 0 | 5 | 10 | 2 | 3 | 0 | 0 | 0 | 146.14 | 1 | | | 2 | | | | | | 1 | | | | | | | | | | | | | | | | | | | | | |
| | Glutamic acid | $C_5H_9NO_4$ | $COOH(CH_2)_2CH(NH_2)COOH$ | 0 | 5 | 9 | 1 | 4 | 0 | 0 | 0 | 147.13 | 2 | | | 1 | | | | | | | | | | | | | | | | | | | | | | | | | | | |
| | Leucine | $C_6H_{13}NO_2$ | $(CH_3)_2CHCH_2CH(NH_2)COOH$ | 0 | 6 | 13 | 1 | 2 | 0 | 0 | 0 | 131.17 | 1 | | | 1 | | | | | | | | | | 1 | | | | | | | | | | | | | | | | | |
| | Isoleucine | $C_6H_{13}NO_2$ | $CH_3CH_2CH(CH_3)CH(NH_2)COOH$ | 0 | 6 | 13 | 1 | 2 | 0 | 0 | 0 | 131.17 | 1 | | | 1 | | | | | | | | | | 1 | | | | | | | | | | | | | | | | | |
| | Lysine | $C_6H_{14}N_2O_2$ | $H_2N(CH_2)_4CH(NH_2)COOH$ | 0 | 6 | 14 | 2 | 2 | 0 | 0 | 0 | 146.19 | 1 | | | 2 | | | | | | | | | | | | | | | | | | | | | | | | | | | |
| | Methionine | $C_6H_{14}N_2O_2$ | $CH_3S(CH_2)_2CH(NH_2)COOH$ | 0 | 6 | 14 | 2 | 2 | 0 | 0 | 0 | 146.19 | 1 | | | 1 | | | | | | | | | | 1 | | | | 1 | | | | | | | | | | | | | |
| | Histidine | $C_6H_9N_3O_2$ | $NHCH=NCH=CCH_2CH(NH_2)COOH$ | 0 | 6 | 9 | 3 | 2 | 0 | 0 | 0 | 155.15 | 1 | | | 1 | 1 | 1 | | | | | | | | | | | | | | | | | 1 | | | | | | | | |
| | Arginine | $C_6H_{14}N_4O_2$ | $HN=C(NH_2)NH(CH_2)_3CH(NH_2)COOH$ | 0 | 6 | 14 | 4 | 2 | 0 | 0 | 0 | 174.20 | 1 | | | 1 | 2 | | | | 1 | | | | | | | | | | | | | | | | | | | | | | |
| | Phenylalanine | $C_9H_{11}NO_2$ | $PhCH_2CH(NH_2)COOH$ | 0 | 9 | 11 | 1 | 2 | 0 | 0 | 0 | 165.19 | 1 | | | 1 | | | | | | | | | | | | | | | | | 1 | | | | | | | | | | |
| | Tyrosine | $C_9H_{11}NO_3$ | $HOPhCH_2CH(NH_2)COOH$ | 0 | 9 | 11 | 1 | 3 | 0 | 0 | 0 | 181.19 | 1 | | 1 | 1 | | | | | | | | | | | | | | | | 1 | | | | | | | | | | | |
| Amino acid | Tryptophan | $C_{11}H_{12}N_2O_2$ | $PhNHCH=CH_2CH(NH_2)COOH$ | 0 | 11 | 12 | 2 | 2 | 0 | 0 | 0 | 204.23 | 1 | | | 1 | 1 | | | | | | | | | | | | | | | | | | | 1 | | | | | | | |
| Carboxylic acid | Ketoglutaric acid | $C_5H_6O_5$ | $(COOH)C=O(CH_2)_2COOH$ | 0 | 5 | 6 | 0 | 5 | 0 | 0 | 0 | 146.10 | 2 | | | | | | | | | | 1 | | | | | | | | | | | | | | | | | | | | |
| Pyridinecarboxylic acid | Nicotinic acid/Vitamin B3 | $C_6H_5NO$ | pyridine-COOH | 0 | 6 | 5 | 1 | 1 | 0 | 0 | 0 | 107.11 | 1 | | | | | | | | | | | | | | | | | | | | | 1 | | | | | | | | | |
| | Glucosaminic acid | $C_6H_{12}NO_6$ | $CH_2(OH)(HCOH)_3CH(NH_2)COOH$ | 0 | 6 | 12 | 1 | 6 | 0 | 0 | 0 | 194.16 | 1 | | 4 | 1 | | | | | | | | | | | | | | | | | | | | | | | | | | | |
| Aminocarboxylic acid | Pantothenic acid/Vitamin B5 | $C_9H_{17}NO_5$ | $CH_2(OH)C(CH_3)_2CH(OH)C(=O)NH(CH_2)_2COOH$ | 0 | 9 | 17 | 1 | 5 | 0 | 0 | 0 | 219.23 | 1 | | 2 | | | | | | | 1 | | | | 2 | | | | | | | | | | | | | | | | | |
| | Aminobenzoyl-triglutamic acid/4-aminobenzoyl-pentanedioic acid | $C_{22}H_{28}N_4O_{13}$ | NA | 0 | 22 | 28 | | 13 | 0 | 0 | 0 | 500.45 | 1 | | | 1 | 1 | | | | | 1 | | | | | | | | | | 1 | | | | | | | | | | | |
| | Pteroyl-triglutamic acid/Folic acid/vitamin B9 | $C_{29}H_{33}N_9O_{12}$ | pteridine ring - para-amino-benzoic acid - glutamic acid | 0 | 29 | 33 | 9 | 12 | 0 | 0 | 0 | 699.63 | 1 | | | 1 | 1 | | | | | 1 | | | | | | | | | | | | | | | | | | | | | 1 |
| Aminobenzoylcarboxylic acid | Aminobenzoylpolyglutamic acid | NA | NA | NA | NA | NA | NA | NA | NA | NA | NA | NA | 1 | | | 1 | 1 | | | | | 1 | | | | | | | | | | 1 | | | | | | | | | | | |
| | Monimethylamine | $CH_5N$ | $CH_3NH_2$ | 0 | 1 | 5 | 1 | 0 | 0 | 0 | 0 | 31.06 | | | | 1 | | | | | | | | | | 1 | | | | | | | | | | | | | | | | | |
| | Dimethylamine | $C_2H_7N$ | $NH(CH_3)_2$ | 0 | 2 | 7 | 1 | 0 | 0 | 0 | 0 | 45.08 | | | | | 1 | | | | | | | | | 1 | | | | | | | | | | | | | | | | | |
| Methyl amine | Trimethylamine | $C_3H_9N$ | $N(CH_3)_3$ | 0 | 3 | 9 | 1 | 0 | 0 | 0 | 0 | 59.11 | | | | | | 1 | | | | | | | | 2 | | | | | | | | | | | | | | | | | |
| Polyamine | Putrescine | $C_4H_{12}N_2$ | $(CH_2)_4(NH_2)_2$ | 0 | 4 | 12 | 2 | 0 | 0 | 0 | 0 | 88.15 | | | | 2 | | | | | | | | | | | | | | | | | | | | | | | | | | | |
| | Tryptone | $C_3H_5NO$ | $H_2C=CO(NH_2)$ | 0 | 3 | 5 | 1 | 1 | 0 | 0 | 0 | 71.08 | | | | 1 | | | | | | | | | 1 | | | | | | 1 | | | | | | | | | | | | |
| Peptide | Peptone | NA | NA | 0 | 13 | 24 | 0 | 4 | 0 | 0 | 0 | 244.33 | NA | NA | NA | | | | | | | | | | | | | | | | | | | | | | | | | | | | |
| Hydroxypyridine | Pyridoxamine/Vitamin B6 | $C_8H_{12}N_2O_2$ | $NH_2(CH_2)C=C(OH)C(CH_3)=NC=C(H_2COH)$ | 0 | 8 | 12 | 2 | 2 | 0 | 0 | 0 | 168.19 | | | 2 | 1 | | 1 | | | | | | | | 1 | | | | | 2 | | | 1 | | | | | | | | | |
| Benzoate | Benzoate | $C_7H_5O_2^-$ | NA | -1 | 7 | 5 | 0 | 2 | 0 | 0 | 0 | 121.11 | 1 | | | | | | | | | | | | | | | | | | | 1 | | | | | | | | | | | |
| | 3-hydroxybenzoate | $C_7H_5O_3^-$ | NA | -1 | 7 | 5 | 0 | 3 | 0 | 0 | 0 | 137.11 | 1 | | 1 | | | | | | | | | | | | | | | | | 1 | | | | | | | | | | | |
| Hydroxybenzoate | 4-hydroxy-benzoate | $C_7H_5O_3^-$ | NA | -1 | 7 | 5 | 0 | 3 | 0 | 0 | 0 | 137.11 | 1 | | 1 | | | | | | | | | | | | | | | | | 1 | | | | | | | | | | | |
| Methyoxybenzoate | 3,4,5-trimethyoxybenzoate | $C_{10}H_{12}O_5^-$ | $(CH_3O)_3$-benzoate | -1 | 10 | 12 | 0 | 5 | 0 | 0 | 0 | 212.20 | 1 | | | | | | | | | | | | | | 3 | | | | | 1 | | | | | | | | | | | |
| Phenol | Phenol | $C_6H_6O$ | $(C_6H_5)OH$ | 0 | 6 | 6 | 0 | 1 | 0 | 0 | 0 | 94.11 | | | 1 | | | | | | | | | | | | | | | | | 1 | | | | | | | | | | | |
| Pyrrolizine | Heliotrine | $C_{16}H_{27}NO_5$ | NA | 0 | 16 | 27 | 1 | 5 | 0 | 0 | 0 | 313.39 | | | 1 | | | | | | | | | | 1 | | | | | | | | | | | | | | | 1 | | | |
| Nucleoside | Inosine | $C_{10}H_{12}N_4O_5$ | NA | 0 | 10 | 12 | 4 | 5 | 0 | 0 | 0 | 268.23 | | | | | | | | | | | | | | | | | | | | | | | | | | | 1 | | | 1 | |
| Methylglycine | Betaine/Trimethylglycine | $C_5H_{11}NO_2$ | $(CH_3)_3N^+CH_2COO^-$ | 0 | 5 | 11 | 1 | 2 | 0 | 0 | 0 | 117.15 | | 1 | | | | 1 | | | | | | | | 2 | | | | | | | | | | | | | | | | | |
| | 4-OHB/4-hydroxybutanoic acid | $C_4H_8O_3$ | $COOH(CH_2)_3OH$ | 0 | 4 | 8 | 0 | 3 | 0 | 0 | 0 | 104.10 | 1 | | 1 | | | | | | | | | | | | | | | | | | | | | | | | | | | | |
| Fatty acid | Oleic acid | $C_{18}H_{34}O_2$ | $CH_3(CH_2)_7CH=CH(CH_2)_7COOH$ | 0 | 18 | 34 | 0 | 2 | 0 | 0 | 0 | 282.46 | 1 | | | | | | | | | | | | | | | | | | 1 | | | | | | | | | | | | |

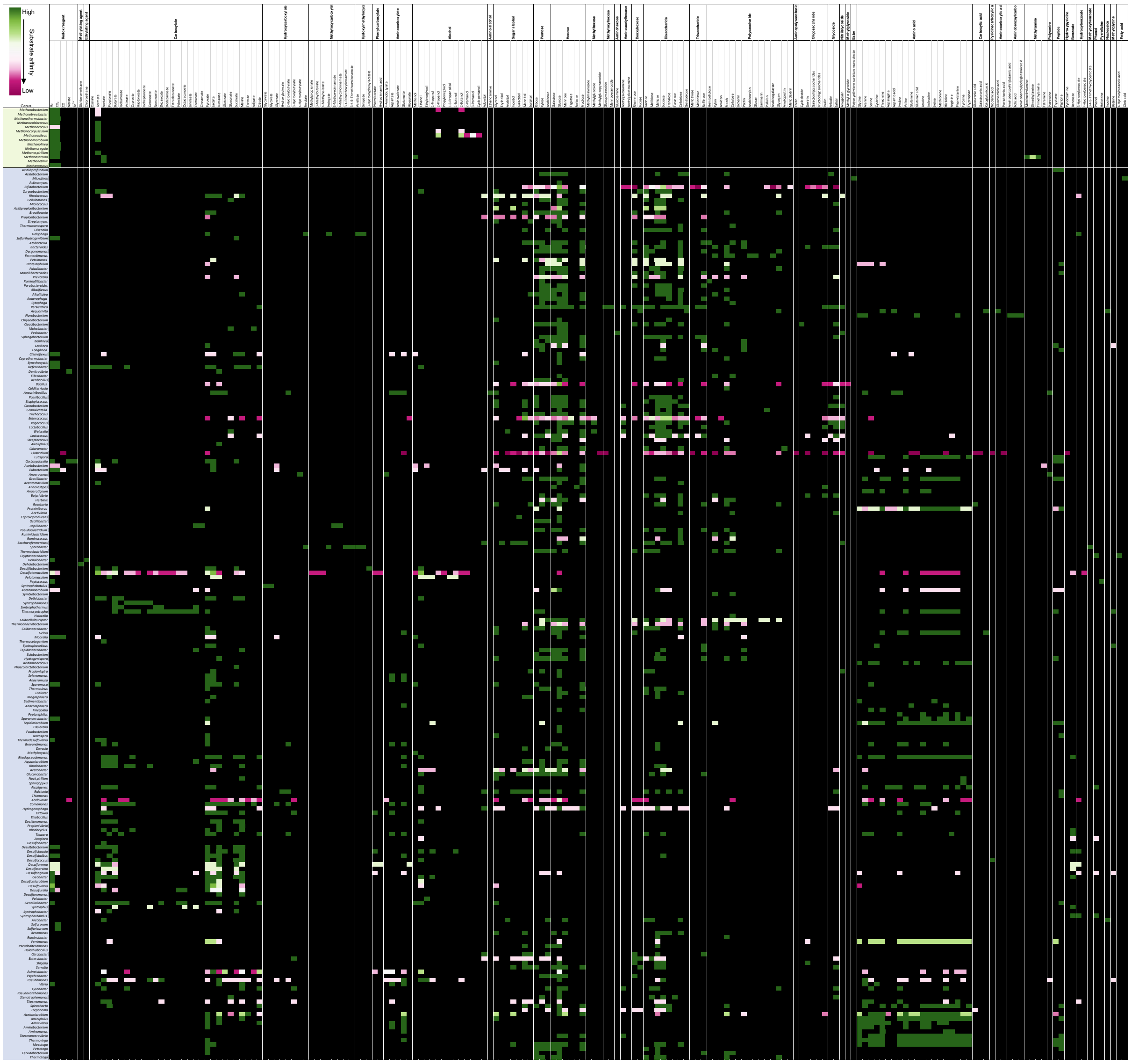

**Supplementary Figure 1.1.** AD substrates, temperatures and pH ranges reported to support microbial growth of different genera from a literature review of 197 papers **a)** Substrates reported to support microbial growth. 187 substrates were investigated. Affinity was based on substrate utilisation where high affinity (green) means that most species within that genera are capable of growth on that substrate, and pink indicates only a proportion of species are capable of growth on that substrate. Black indicates no reported growth on a given substrate. Substrates are categorised by key functional groups [1-195].

#### 1.2. Literature review of genera reported to be involved in syntrophic interactions in anaerobic digestion

**Table 1.6.** Genera reported to be involved in syntrophic relationships in anaerobic digestion. Reported growth conditions are noted including pH range, optimum pH, temperature (temp) range (°C), optimum temp (°C). Whether the genus has been reported to be capable of DIET is indicated by Y for yes, or NA for information not available. Potential function reported in literature is noted and references are provided. *Abbreviations Direct Interspecies Electron Transfer (DIET); Hydrogen Formate Interspecies Transfer (HFIT); Syntrophic Acetate Oxidising Bacteria (SAOB), data not available (NA). [11,14,23,56,67,79,92,99,121,123,127,132,141,142,176,177,196,199-215]

| GENUS | DIET | REPORTED FUNCTION | REF |
|---|---|---|---|
| *Methanomassiliicoccus* | NA | Hydrogenotrophic methanogen | 199,200 |

| | | | |
|---|---|---|---|
| *Methanobacterium* | NA | Hydrogenotrophic methanogen | 56,123,141,177,199,201-203 |
| *Methanobrevibacter* | NA | Hydrogenotrophic methanogen | 23 |
| *Methanothermobacter* | NA | Hydrogenotrophic methanogen | 199,204 |
| *Methanoculleus* | NA | Hydrogenotrophic methanogen, acetate turnover | 92,99,132,204 |
| *Methanolinea* | NA | Hydrogenotrophic methanogen | 203,205 |
| *Methanospirillum* | NA | Hydrogenotrophic methanogen | 23,56,99,123,203 |
| *Methanosarcina* | Y | Acetate turnover | 56,92,177,203,205,206 |
| *Methanothrix* | Y | Obligate acetoclastic methanogen | 56,99,123,141,203,206,207 |
| *Bifidobacterium* | NA | Acid formation and hydrogen release | 14 |
| *Bacillus* | NA | Acid formation and hydrogen release | 14 |
| *Exiguobacterium* | NA | SAOB | 199,208 |
| *Streptococcus* | Y | Acid formation and hydrogen release | 14 |
| *Butyrivibrio* | NA | Acid formation and hydrogen release | 14 |
| *Caloramator* | Y | | 56 |
| *Clostridium* | NA | SAOB | 14,92,207,209 |
| *Desulfotomaculum* | NA | Propionate oxidiser, sulfate-reducer, acetate consumer | 14,23 |
| *Pelotomaculum* | NA | Acetogenic propionate oxidiser | 23,121,123,204 |
| *Syntrophomonas* | Y | 4-8 carbon short-chain fatty acid degrader, propionate/acetate producer, SAOB | 14,56,67,79,121,123,127,142,176,202,204,206,207,209 |
| *Syntrophothermus* | NA | Acetogen | 204 |
| *Caldanaerobacter* | NA | SAOB | 199 |
| *Thermacetogenium* | NA | SAOB | 176,204 |
| *Syntrophaceticus* | NA | | 92,99 |
| *Tepidanaerobacter* | NA | SAOB | 56,199 |
| *Propionispira* | NA | Acetogenesis | 206 |
| *Anaeroarcus* | Y | Acetate and propionate production, amino acid degradation | 56,216 |
| *Sporanaerobacter* | Y | | 56,206 |
| *Bacteroides* | Y | Acid formation and hydrogen release | 14,56,205 |
| *Prosthecochloris* | Y | Sulfide/sulfur reducer | 142,217 |
| *Sulfurospirillum* | Y | | 56,218 |
| *Candidatus Cloacamonas* | NA | Hydrogen producer, amino acid fermenter, butyrate/propionate oxidation | 123,150,219 |
| *Anaerolinea* | NA | Short chain fatty acid degrader, acetate producer | 129,220 |
| *Coprothermobacter* | NA | SAOB | 199 |
| *Deferribacter* | Y | | 56 |
| *Thermodesulfovibrio* | NA | Syntrophic mediator of lactate degradation | 204 |
| *Thauera* | Y | | 56 |
| *Shewanella* | Y | | 56,67,221 |
| *Pseudomonas* | Y | Acid formation and hydrogen release, protein fermentation | 14,196 |
| *Vibrio* | NA | | 67 |
| *Tepidiphilus* | NA | SAOB | 199 |
| *Cloacibacillus* | NA | SAOB, $H_2/CO_2$ producer, acetic acid production, amino acid fermentation | 210,211 |
| *Synergistes* | NA | Amino acid fermenter, VFA producer | 205,212 |

| | | | |
|---|---|---|---|
| *Candidatus Desulfofervidus* | Y | Sulfate-reducer, methane oxidation | 56,222 |
| *Desulfobacterium* | Y | | 56 |
| *Desulfobacula* | Y | | 56 |
| *Desulfovibrio* | Y | Hydrogen-producing acetogen, lactate and ethanol degradation | 14,23,56,207 |
| *Desulfuromonas* | Y | | 56 |
| *Geoalkalibacter* | Y | | 56,142 |
| *Geobacter* | Y | HFIT, ethanol degradation | 56,142,207,213 |
| *Smithella* | Y | Propionate oxidiser, produces acetate and butyrate | 23,79,132,214 |
| *Syntrophus* | Y | HFIT | 121,123,142,205 |
| *Syntrophobacter* | NA | Butyrate/ propionate to acetate conversion, propionate oxidiser | 11,14,23,121,205,207 |
| *Mesotoga* | NA | Acetate oxidiser | 204 |
| *Pseudothermotoga* | NA | Acetate oxidiser | 204,215 |
| *Thermotoga* | NA | Acetate oxidiser | 176 |

### 2. Supplementary Materials and Methods

#### 2.1. Digester description

Two experimental runs were designed to address the latter two research objectives using continuously stirred tank reactor (CSTR) experiments and Quorn™ MFWW as feedstock. In run 1 mesophilic and thermophilic reactors treating MFWW were run, and in run 2 single- and two-stage CSTRs treating MFWW were run (Figure 2b).

CSTRs were selected due to their widespread application (Figure 1f). Anaero technology patented single- and multi-stage CSTRs with temperature and gas sensors were used in this experiment. The patented reactor technology enabled advanced research of AD as the bioreactors mimic commercial-scale AD processes and the inclusion of gas composition sensors, and real-time biogas and temperature detection offer advanced system monitoring. Furthermore, Anaero technology flexible modular bioreactor design enabled the development of a two-stage system for comparison of single- and multi-stage systems.

CSTRs were run with a consistent 14.7 day HRT, 30rpm mixing, and 0.32L/d influent flow rate (Supplementary Table 2.1). OLR was based on influent COD (g COD/L/d). All CSTRs were inoculated with inoculum sourced from a UK-based mesophilic AD plant treating mixed food waste. CSTR's were inoculated with 5.5L of inoculum with continuous mixing of the initial inoculum between inoculations to ensure homogeneity between reactors. Before starting the experimental run, reactors were operated for approximately two months at low OLR (0.18L/d influent flow rate) to ensure all food waste was eliminated and the inoculum was acclimated to the new feedstock. We quantified volatile soluble solids (VSS) as a measure of active biomass on the first and last day of the experimental run. Bioreactor chemistry and influent composition were monitored through the experiment, sampling in triplicate according to Figure 2d-e, including pH, VFA, COD, and sugar and sugar alcohol concentration (Supplementary Table 2.1). Metagenomic sequencing of 16s rRNA was conducted to investigate the microbiome underpinning AD, and OTU assignment and taxonomic classification were carried out over the course of the experiment (Figure 2). Performance was measured in terms of COD removal efficiency, cumulative biogas production, and specific daily biogas production (Figure 2).

**Table 2.1.** CSTR experimental design, including operational parameters, bioreactor chemistry, feedstock composition, and performance parameters. *Abbreviations: continuously stirred tank reactor (CSTR); organic loading rate (OLR), hydraulic retention time (HRT), volatile fatty acid (VFA), chemical oxygen demand (COD).

| Parameter type | Parameter | Value | Unit |
|---|---|---|---|
| **Operational** | Flow rate | 0.32 | L/d |
| | OLR | 0.19 to 1.34 | g COD/L/day |
| | HRT | 14.7 | days |
| | Temperature | 37 or 50 | ºC |
| | Mixing speed | 30.0 | rpm |
| **Chemical** | pH | Variable | n/a |
| **(Influent & reactor)** | VFA | Variable | g/L |
| | Sugar and sugar alcohol | Variable | g/L |
| **Biological** | 16s rRNA sequencing | Variable | n/a |
| **Performance** | Biogas | Variable | L/g COD/d |
| | COD removal | Variable | % |

##### 2.1.1. Run 1

All reactors within run 1 had the same influent through individual peristaltic pumps. Reactors passively outputted effluent through a swan neck outlet. Reactors were sampled in a time-series over a 52-day run time to investigate temporal scales (Figure 2d). Each reactor configuration was run in triplicate, where CSTR1.1-3 represent mesophilic and CSTR2.1-3 represent thermophilic reactors (Figure 2b).

##### 2.1.2. Run 2

All reactors within run 2 had the same influent through individual peristaltic pumps. Single- and second-stage reactors passively outputted effluent through a swan neck outlet. For two-stage reactors, the first-stage effluent transferred to the second-stage through a hose connection (Figure 2c). Reactors were sampled in a time series over a 50-day run time to investigate temporal scales (Figure 2e). Each reactor configuration was run in quadruplicate, where CSTR3.1-4 represent single-stage reactors, CSTR4.1-4 represent first-stage reactors, and CSTR5.1-4 represent second-stage reactors (Figure 2b). DNA quality for day 28 was not sufficient to allow sequencing (Figure 2e).

#### 2.2. Volatile Suspended Solid Content

Total suspended solids (TSS) and volatile suspended solids (VSS) were measured according to APHA standard methods using a 0.22μm glass fibre filter and a vacuum filtration system (Eq. S1, Eq. S2) [223].

Where TSS denotes the total soluble solids (mg/L), $W_1$ represents the dish weight (g), $W_2$ represents the heated sample and dish weight (g) and SV represents the sample volume (mL). VSS denotes volatile soluble solids of total soluble solids (mg/L), $W_2$ represents heated sample weight (g), $W_3$ represents the variable ash and dish weight (g), and SV represents the sample volume (mL).

$$TSS = \frac{W_2 - W_1}{SV} \times 10^6 \quad \text{(S1)}$$

$$VSS = \frac{W_3 - W_2}{SV} \times 10^6 \quad \text{(S2)}$$

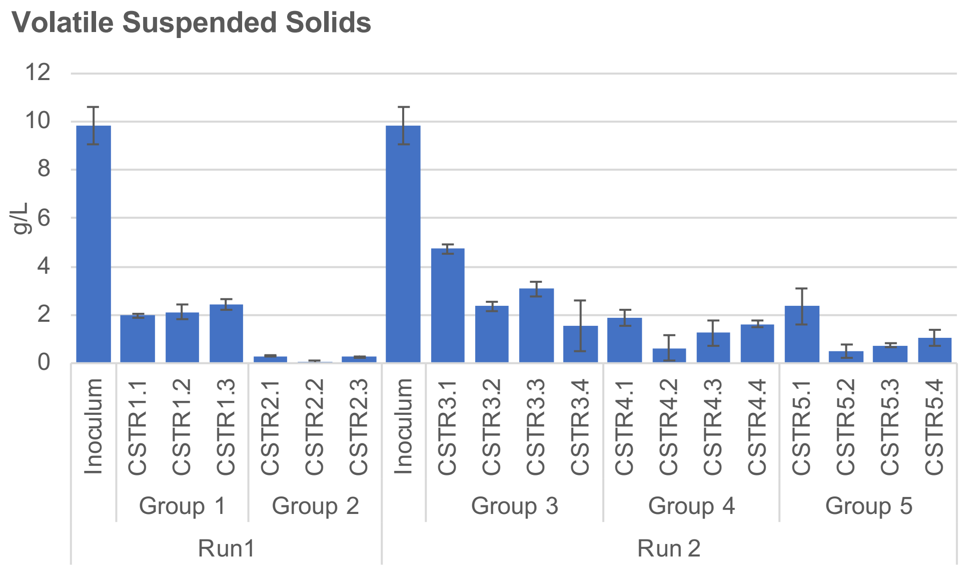

**Supplementary Figure 2.1.** Volatile suspended solid concentration, g/L of inoculum and experimental reactors. *Abbreviations: continuously stirring tank reactor (CSTR).

#### 2.3. Chemical oxygen demand

For high range COD (200mg/L to 15,000mg/L), a 0.2mL aliquot of the test sample was added to separate vials containing 61% sulfuric acid, silver sulfate, and potassium dichromate (Vario HR/COD 200mg/L to 15,000mg/L test reagent, Lovibond). For medium range (20mg/L to 1,500mg/L) and low range (3mg/L to 150mg/L) a 2mL aliquot of test sample was added to separate vials containing 61% sulfuric acid, silver sulfate, and potassium dichromate (Vario MR/COD 200mg/L to 1,500mg/L test reagent, and Vario LR/COD 3mg/L to 150mg/L test reagent, Lovibond).

HPLC-grade water was used as a negative control. 5000mg/L (high range) or 100mg/L (medium and low range) COD standards were used as a positive control (Lovibond). Closed vials were mixed by inverting 2-3 times and incubated at 150°C for 2hrs in the thermoreactor (Thermoreactor RD 125, Lovibond) according to standard methods (ISO, 1989; Moore et al., 1949). Samples were allowed to cool to room temperature and mixed by inverting 2-3 times. Solids were allowed to settle, and samples were measured at 600nm wavelength using the MD-600 spectrophotometer (Lovibond).

COD removal (%) was calculated from measured effluent and influent COD, g/L (Eq. S3) $Inf_{COD}$ denotes influent chemical oxygen demand (g/L), and $Eff_{COD}$ represents effluent chemical oxygen demand (g/L).

$$COD\ removal\ efficiency = \frac{Inf_{COD}}{Eff_{COD}} \quad \text{(S3)}$$

#### 2.4. High performance liquid chromatography

HPLC analysis was conducted using an LC-40D XR solvent delivery pump (Shimadzu) and CTO 40C column oven (Shimadzu). A sample volume of 10µL was injected using a SIL-40C XR Autosampler UHPLC autoinjector (Shimadzu). The mobile phase was degassed by a DGU-405 degassing unit (5 channel; Shimadzu). The column and autosampler were purged with mobile phase before and after every analysis. Calibration curves were constructed by plotting peak area against compound concentration. Linearity of the line of best fit was calculated by the least square regression method within the LabSolutions software. The Limits of Detection (LOD) and Limits of Quantification (LOQ) were calculated using LabSolutions software. These values were used to verify the quality of the calibration curve. Samples (0.6 mL) were filtered through a 0.22µm pore filter (Claristep® Filtration system, Satorius) into a clean 1.5mL HPLC vial. Sample analysis was carried out in triplicate and a mean value with standard deviation was calculated.

##### 2.4.1. Sugar and sugar alcohol

HPLC analysis was conducted using an SPD-M40 photodiode array (PDA) detector (Shimadzu), a RID-20A refractive index detector operating at 40°C (Shimadzu), and a Rezek RCM-Monosaccharide $Ca^{2+}$ column (8µm, 100mm x 7.8mm internal diameter; Phenomenex) maintained at 80°C. HPLC grade water was used as a mobile phase with a flow rate of 0.2mL/min and an isocratic elution run time of 1hr. Instrument parameters for sugar and sugar alcohol HPLC analysis are detailed in Supplementary Table 2.4.1 [224].

Standard stock solutions (10g/L) were prepared using ultrapure HPLC grade water (Arium® Pro Ultrapure Water Systems, Satorius). Five dilution levels (100, 200, 500, 1000 and 2000mg/L) were run in triplicate for the 7 reference compound stock solutions: alpha-D(+)-Melibiose (neat, Ehrenstorfer GMBH), Glucose (1000mg/L in pure water, TraceCERT® Supelco® Merck), D-(+)-Xylose (neat, Ehrenstorfer GMBH), Maltitol (neat, Supelco® Merck), Glycerol (neat, Supelco® Merck), Mannitol (neat, Supelco® Merck), and D-(+)-Arabitol (neat, Ehrenstorfer GMBH). Sample analysis was carried out using the same method as the standard solutions for sugar and sugar alcohol content (Supplementary Table 2.4.1). Peak identification was carried out using the calibration curve for sugar and sugar alcohol (Supplementary Figure 2.4.1).

**Supplementary Table 2.4.1.** HPLC sugar and sugar alcohol content instrument parameters.

| | |
|---|---|
| Detectors | Photodiode Array (210nm) and Refractive Index Detector (40ºC) |
| Column | Phenomenex Rezek-RCM Monosaccharide $Ca^{2+}$ |
| Column Oven Temperature, ºC | 80 |
| Mobile Phase | Ultrapure HPLC-grade Water |
| Flow rate, mL/min | 0.2 |
| Time, min | 60 |
| Injection volume, **μ**L | 10 |

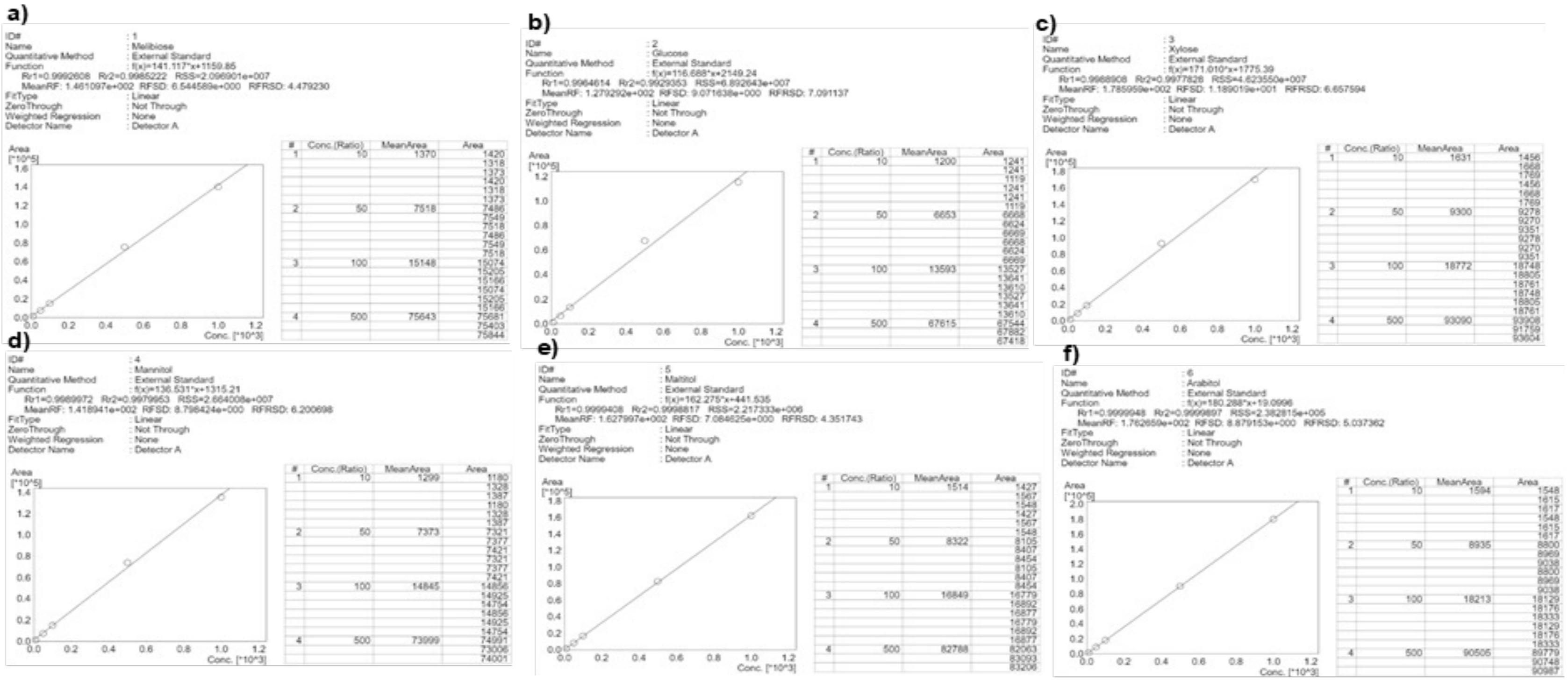

| Compound | $R^2$ |
|---|---|
| Melibiose | 0.999 |
| Glucose | 0.993 |
| Xylose | 0.998 |
| Mannitol | 0.998 |
| Maltitol | >0.999 |
| Arabitol | >0.999 |

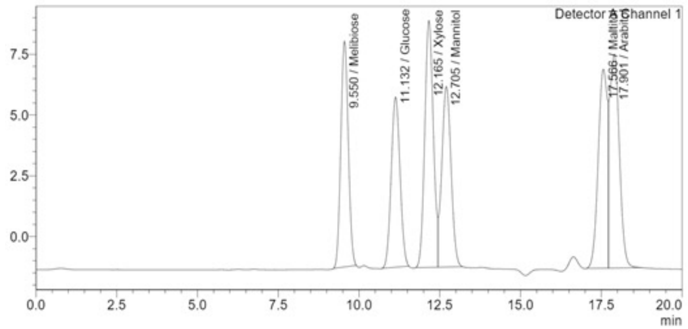

**Supplementary Figure 2.4.1..** Sugar and sugar alcohol calibration curves. **(a)** Melibiose. **(b)** Glucose. **(c)** Xylose. **(d)** Mannitol. **(e)** Maltitol. **(f)** Arabitol. Five levels of concentration were used: 10, 50, 100, 500 and 1000mg/L were used for level 1, 2, 3, 4 and 5 respectively. Chromatogram of standard solutions. Analysis was conducted in triplicate and repeated. Both runs shows high reproducibility and were both included in the final calibration curve. R2 values are provided for each compound calibration.

##### 2.4.2. Volatile fatty acids

A sample volume of 20µL was injected using the autoinjector (Shimadzu) and run through an Aminex® HPX-87H column (8µm, 300mm x 7.8mm internal diameter; BioRad) maintained at 55°C. $H_2SO_4$ (0.008M HPLC grade) was used as a mobile phase with a flow rate of 0.6mL/min and an isocratic elution run time of 40 minutes. Instrument parameters are detailed in Supplementary Table 2.4.2 [225].

**Supplementary Table 2.4.2**. Instrument parameters used for HPLC analysis of VFAs.

| **Detectors** | **Photodiode Array (205nm)** |
|---|---|
| **Column** | Biorad Aminex HPX-87H |
| **Column Oven Temperature, ºC** | 55 |
| **Mobile Phase** | 0.008N $H_2SO_4$ |
| **Flow rate, mL/min** | 0.6 |
| **Time, min** | 40 |
| **Injection volume, µL** | 20 |

Standard stock solutions (10g/L) were prepared using ultrapure HPLC grade water (Arium® Pro Ultrapure Water Systems, Satorius). Five dilution levels (0.01, 0.05, 0.10, 0.50, 1.00 and 2.00g/L) were run in triplicate for the 7 reference compound stock solutions: formic acid (≥99%, HiPerSolv CHROMANORM®), acetic acid (≥99.8%, HiPerSolv CHROMANORM®), propionic acid (neat, Supelco® Merck), iso-butyric acid (neat, EHRENSTORFER GMBH), butyric acid (neat, Supelco® Merck), iso-valeric acid (neat, Supelco® Merck) and valeric acid (neat, Supelco® Merck). Calibration curves were constructed by plotting peak area against standard concentration (Supplementary Figure 2.4.2). Sample analysis was carried out using the same method as the standard solutions for volatile fatty acid content (Supplementary Table 2.4.2). Peak identification was carried out using the calibration curve for sugar and sugar alcohol (Supplementary Figure 2.4.2). Acetic acid was used as a positive control and water was used as a negative control for sample analysis.

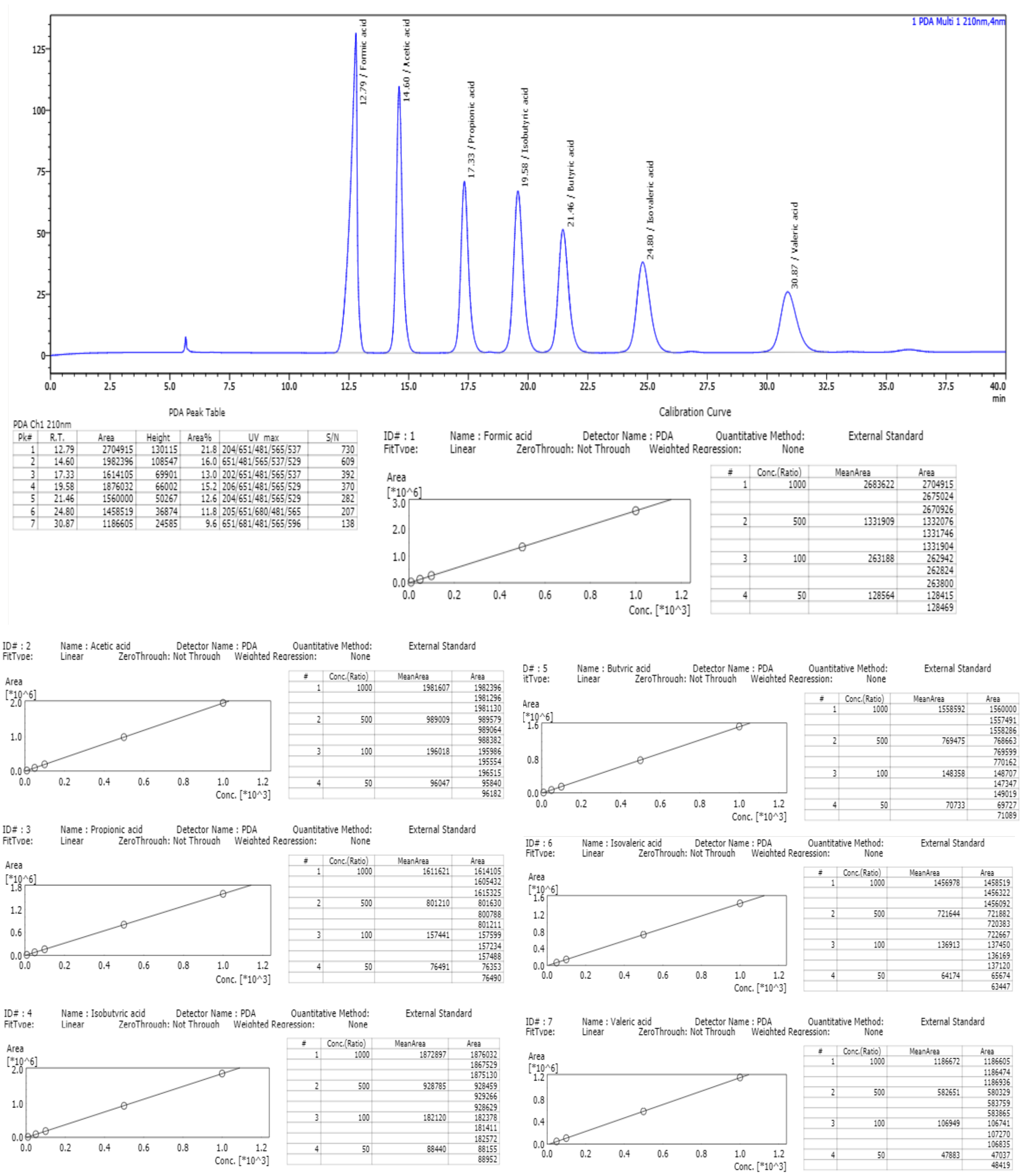

| Compound | $R^2$ |
|---|---|
| **Formic acid** | >0.999 |
| **Acetic acid** | >0.999 |
| **Propionic acid** | >0.999 |
| **Iso-butyric acid** | >0.999 |
| **Butyric acid** | >0.999 |
| **Iso-valeric acid** | >0.999 |
| **Valeric acid** | >0.999 |

**Supplementary Figure 2.4.2.** Combined VFA chromatogram and calibration curves. Calibration curves were constructed using 7 standards, acetic acid, propionic acid, iso-butyric acid, butyric acid, iso-valeric acid and valeric acid. $R^2$ values are given for each compound calibration.

##### 2.4.3. Biomethane potential reactors

Biomethane potential (BMP) CSTR reactors were developed by Anaero technology based on operational research in the UK food waste industry. Each machine consists of a hot water bath incubator for temperature control, 15 1L CSTRs with a centralised mixer, a gas flow meter, a gas composition sensor, built-in bespoke monitoring software (algorithms and controller coded in Python) and cloud data storage, enabling real-time biogas production and composition data.

#### 2.5. DNA extraction, 16S rRNA amplification, barcoding and sequencing

##### 2.5.1. DNA extraction DNeasy® PowerSoil® Pro Kit

2mL of each CSTR sample (in triplicate) was centrifuged at 10,000rpm for 5 minutes, the supernatant was decanted, and the pellet was used for the DNA extraction, which was carried out using the DNeasy® PowerSoil® Pro Kit according to standard protocol (QIAGEN GmbH, Germany). Samples were suspended in solution C6 (QIAGEN GmbH, Germany) and quantified using a NanoDrop™ One (Thermo Scientific, Waltham, MA). Quantified samples were immediately stored at -20°C prior to sequencing.

##### 2.5.2. Illumina 16S rRNA amplicon sequencing

Illumina 16S rRNA V3-V4 amplicon sequencing was conducted externally at the WHRI-Genome-Centre at Queen Mary's University, United Kingdom.

###### 2.5.2.1. DNA sample quantification and quality control

Sample DNA concentration was quantified using a Nanodrop, with Milli-Q water as a negative control and blank. DNA quality was checked using a Qubit Flex Fluorometer. A working solution of 1:200 dilution of Qubit Fluorescent reagent (reagent lot no. 2600187) and Qubit

Dilution Buffer (reagent lot no. 2600189) was prepared. 2 standards were used (Standard lot number 2600187), as well as calf thymus DNA as a positive control. 190µL of the working solution with 10µL of the standards, and 199µL of the working solution with 1µL of the calf thymus DNA, or sample DNA were added to Quibit Flex 8-tube strips. Tubed were capped and vortexed briefly before measuring using the Qubit Flex Fluorometer.

###### 2.5.2.2. Sample PCR using Platinum SuperFi II DNA Polymerase

PCR reagents, forward and reverse primers (Supplementary Figure 2.5.1c) and the sample DNA were combined according to Supplementary Figure 2.5.1a, and vortexed briefly to ensure good mixing, and briefly centrifuged. The PCR cycle was run using a tetrad thermocycle (LG190) according to the program detailed in Supplementary Figure 2.5.1b, with the tetrad lid set to 105ºC.

a)

| Reagent | 1X volume, µL | Component Lot No. |
|---|---|---|
| 5X SuperFi II DNA Buffer | 2.00 | 013011305 |
| Forward and reverse primer, 10µM | 0.20 | 10/01/2024 |
| PCR nucleotide mix, 10µM | 0.20 | 10/01/2021 |
| Platinum SuperFi II DNA Polymerase, 2U/µL | 0.25 | 01299609<br>01337237 |
| RNase-free water | 5.35 | 175027101 |

b)

| Step | Temperature ºC | Time, secs | Cycles |
|---|---|---|---|
| Initial denaturation | 95 | 120 | 1 |
| Denaturation | 95 | 30 | 30 |
| Primer annealing | 55 | 30 | |
| Elongation | 72 | 30 | |
| Final Elongation | 72 | 300 | 1 |
| Hold | 4 | Hold | ∞ |

c)

| | | | | | | |
|---|---|---|---|---|---|---|
| PCR Primer Box 7 | D1 | 16S V3-V4 Forward | ACACTGACGACATGGTTCTACACCTACGGGNGGCWGCAG | Bacteria | 16s v3-v4 | CS1 |
| PCR Primer Box 7 | D2 | 16S V3-V4 Reverse | TACGGTAGCAGAGACTTGGTCTGACTACHVGGGTATCTAATCC | Bacteria | 16s v3-v4 | CS2 |

d)

| Reagent | 1X volume, µL | Component Lot No. |
|---|---|---|
| 10X FastStart high fidelity reaction buffer (without $MgCl_2$) | 1.0 | 74602900 |
| 25mM $MgCl_2$ | 1.8 | 72402300 |
| DMSO | 0.5 | 61443900 |
| 10mM PCR-grade nucleotide mix | 0.2 | 67206126 |
| 5U/µL FastStart high fidelity enzyme blend | 0.1 | 67206126 |
| RNase-free water | 0.4 | 1540233300 |

e)

| Step | Temperature ºC | Time, secs | Cycles |
|---|---|---|---|
| Initial denaturation | 95 | 600 | 1 |
| Denaturation | 95 | 15 | 15 |
| Primer annealing | 60 | 30 | |
| Elongation | 72 | 60 | |
| Final Elongation | 72 | 180 | 1 |
| Hold | 4 | Hold | ∞ |

**Supplementary Figure 2.5.1.** PCR amplification. **(a)** PCR mixture components. **(b)** PCR cycle program. **(c)** Forward and reverse PCR primers for 16S V3-V4 amplicon sequencing. **(d)** Barcoding PCR mixture. **(e)** Amplicon barcoding PCR.

###### 2.5.2.3. Agarose gel electrophoresis

Once the PCR was completed samples were run on a 2% agarose gel (lot no. ESS520-B110700) mixed with 1X TBE Buffer, and GelRed Nucleic acid stain. The gel was added to a gel electrophoresis tank and filled with 1X TBE buffer. Either HyperLadder™ 100bp (Bioline) with a size range of 100 to 1013 bp, or HyperLadder™ 1 Kb (Bioline) with a size range of 200 to 10,037 bp were used as a DNA ladder as appropriate depending on sample DNA size. 2µL of the DNA ladder was loaded into the first and last well of each gel. 1µL of 5X DNA loading buffer and 4µL of the DNA sample were loaded into individual gel channels, and the gel electrophoresis was run at 400V for 30 minutes. The gel was visualised using UVP-BioDoc.

###### 2.5.2.4. Automated Amplicon barcoding PCR

Amplicon barcoding PCR was conducted using BiomekFX robot and automated software (Supplementary Figure 2.5.1d). A master mix of reagents were prepared according to Figure 3.8 and mixed using vortexing and centrifuging. 1X volume of the master mix was added to each sample from the previous PCR. The thermocycling was conducted (Supplementary Figure 2.5.1e). Four wells were run on a D1000 screen tape to check the barcoding had been successful (buffer lot number 0006745353, ScreenTape lot number 0203008-293, Agilent).

###### 2.5.2.5. Library pooling and cleaning

Libraries were cleaned and pooled using AMPure XP Beads. 0.9X the pool volume of AMPure XP beads were added to the library DNA and incubated at room temperature for 10 minutes. The mixture was then spun briefly, and a magnet was used to separate the beads from the liquid to allow supernatant removal. 200µL of freshly prepared 80% ethanol was added to the pellet and incubated for 30 seconds at room temperature. The ethanol was discarded, and this cleaning step was repeated for a total of three times. The tube was then briefly spun, and a magnet was used to remove any residual ethanol. Beads were air dried for 5 to 10 minutes, resuspended in 150uL of EB buffer (lot number 160051261) and incubated at room temperature for 10 minutes. A magnet was used to separate the beads and liquid, and the supernatant was transferred to a clean 1.5mL Lo-bind tube. Quality control was performed on the cleaned pool DNA using the Qubit Fluorometer and D1000 ScreenTape Assay (Supplementary Figure 2.5.1).

###### 2.5.2.6. Illumina MiSeq library denaturation and loading

An equal volume of fresh 0.2N NaOH and DNA library (4nM) were combined in a 1.5mL LoBind microcentrifuge tube and mixed by vortexing and centrifuging. The mixture was incubated at room temperature for 10 minutes. The same volume as used for the library of 200mM Tris-HCL was added to neutralise the NaOH and mixed by briefly vortexing and centrifuging. HT1 Buffer (Lot No. 20789296) was added to a total volume of 1mL, and the sample was mixed by briefly vortexing and centrifuging and stored on ice. The denatured library was diluted using HT1 buffer to the molarity required for loading a total volume of 600uL. 20pM PhiX (Lot No. 20777170) was used as a control. The loading cartridge was mixed by inverting 10 times, and gently tapping, and 600uL of the denatured library was added to the cartridge. Sequencing was run using Illumina MiSeq v3 kit and the run quality was assessed based on quality score (% bases> Q30), raw cluster density (k/mm$^2$), data output (Gbp), read number (Lane 1) and % PhiX aligned (Supplementary Table 2.5.1). Quality control was also assessed using FastQC in the Apocrita environment.

**Supplementary Table 2.5.1.** Quality control of illumina MiSeq v3 kit sequencing

| Metric | Result | Pass/Fail | Comments |
|---|---|---|---|
| **Quality score, % bases > Q30** | 87.17 | Pass | |
| **Raw Cluster Density, k/mm$^2$** | 883±27 | Fail | Good number of reads achieved with high Q30 score |
| **Data Output, Gbp** | 12.69 | Pass | |

| | | |
|---|---|---|
| **Read Number (Lane 1)** | 20,529,084 | Pass |
| **PhiX Aligned, %** | 18.04 | Pass |

##### 2.5.3. Taxonomic classification (Mothur, Epi2me)

Illumina sequencing data taxonomic classification was conducted using the Mothur pipeline available in Galaxy referenced against the Silva_v4 database [226,227]. Additional pre-processing steps were conducted which are presented in Supplementary Figure 2.5.2. Species relative abundance (%) was calculated from taxonomic classification counts divided by total reads. The taxonomic profile was assessed on the phylum level for all reactors. *Firmicutes, Proteobacteria*, and *Bacteroidetes* are commonly reported to significantly affect AD [228,229]. To further investigate the taxonomic profile of these key phyla, the relative abundance of *Firmicutes, Bacteroidetes*, and *Proteobacteria* is presented at the class, order, and family level. The taxonomic profile was investigated in terms of the relative abundance of total archaea for the lowest common ancestor down to the genus level. The relative abundance of methanogens as a percentage of the total archaea population was compared by methanogenic pathway, including obligate acetoclastic (*Methanosaeta*), acetoclastic and hydrogenotrophic (*Methanosarcina*), and hydrogenotrophic. The relative abundance of syntrophic species reported in the literature was compared across reactors. Correlations between syntrophic bacteria and methanogens were investigated by calculating Spearman's rank correlation coefficients.

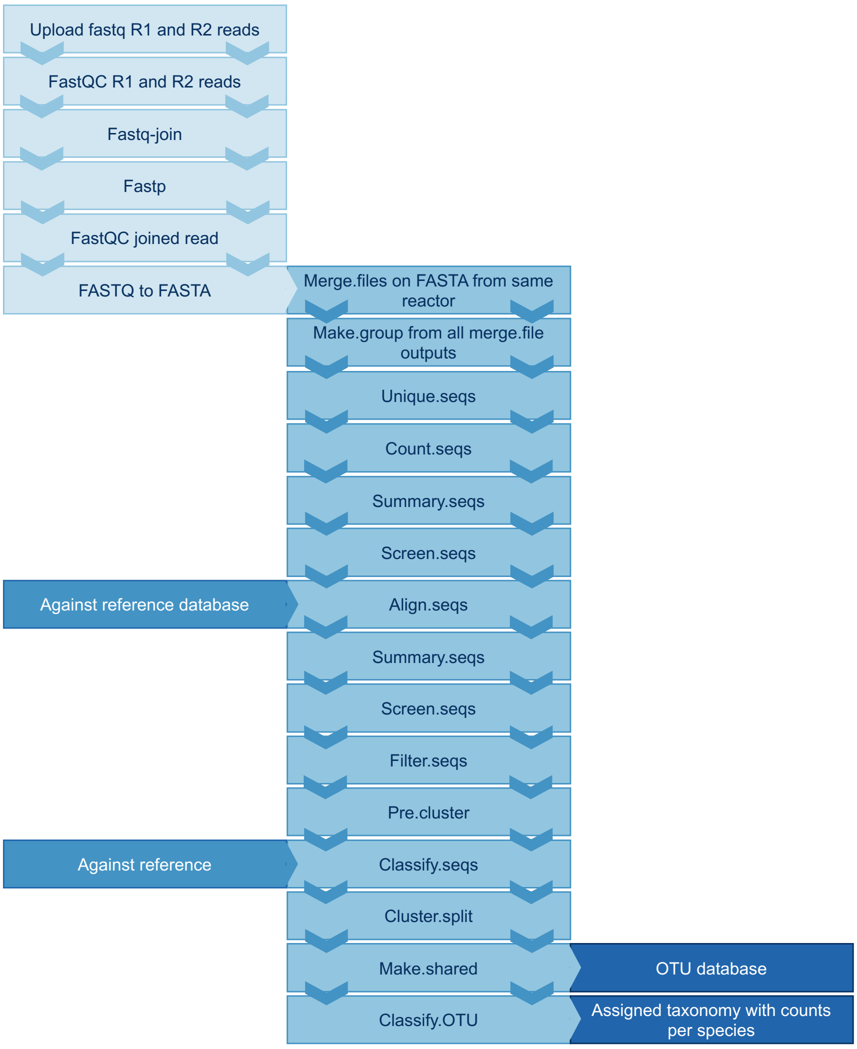

**Supplementary Figure 2.5.2.** Bioinformatic pipeline for metagenomic analysis using a pipeline within galaxy based on Mothur pipeline with added quality control step prior to merging FASTA files [230]. Sequences were aligned against SilvaV4 [227]. Outputs include an OTU database, and taxonomic assignment with read counts.

#### 2.6. Network analysis

The Molecular Ecology Network Analysis (MENA) pipeline was used to conduct network analysis [231,232]. Relative abundance of each reactor was analysed across 17 time points. Genes present in nine out of 17 samples per reactor were retained, and missing data was filled with zero values. A logarithm transformation for non-compositional data was applied. The similarity matrix was constructed based on Pearson correlation in time series, allowing one-time point lag for samples following a time series without replicates for individual reactors, and with biological replicates for reactor groups. The calculation order was based on a decrease in the cut-off from the top, and a regression Poisson distribution was applied. A Chi-square test on the Poisson distribution was applied to calculate the cut-off of $p<0.001$. Global network properties, individual nodes' centrality, module separation, and modularity calculation were performed in MENA [232]. A greedy modularity, leading eigenvector and short-random walks optimisation separations were used, and Z and P values were calculated for all nodes. Networks were analysed using regular power, exponential, and truncated power law. The network was randomised, and the network properties were then recalculated to validate the structure. The network file was imported into Cytoscape for visualisation, filtering by genus-level taxonomic classification to simplify the visualisation [231].

#### 2.7. Alpha diversity

Alpha diversity was measured using Shannon, Simpson's and Chao1 diversity indices. The Shannon index estimates species richness and species evenness based on the number of unique species, and relative abundances (Eq. S4). The Shannon diversity Index, alpha diversity was also characterised using Simpson's Diversity Index to assess species evenness and richness (Eq. S5). The Simpson index is more sensitive to the most abundant species within the community and tends to emphasise the dominance of common or abundant species. Therefore, communities with fewer dominant species will have a lower Simpson diversity. Simpson's Diversity Index values range from 0 to 1, where 1 indicates complete diversity and 0 indicates no diversity. The Chao1 Diversity Index, an estimator of species richness in terms of the number of unique species present in a given community, without taking into account relative abundances [233]. Chao1 provides an estimate of the minimum number of species present in a community, considering both the observed species and an estimate of the unobserved species based on the number of rare species, specifically those observed once or twice (Eq. S6). Significance was calculated using a one-way ANOVA with post hoc-Tukey test for reactor averages.

Where $H'$ denotes the Shannon diversity index and $p_i$ denotes the proportion of individual species $i$ relative to the total number of individuals within the community, $S_{ob}$ denotes the number of observed species, $F_1$ denotes the number of species only observed once, and $F_2$ denotes the number of species only observed twice.

$$Shannon = -\sum(p_i log p_i) \quad \text{(S4)}$$

$$Simpson's = \sum {p_i}^2 \quad \text{(S5)}$$

$$Chao1 = S_{ob} + \frac{{F_1}^2}{2F_2} \quad \text{(S6)}$$

#### 2.8. Beta diversity

The Bray-Curtis similarity index, a non-metric measure of the compositional dissimilarity between two different samples [12]. This method was used to quantify the degree of similarity in species composition between pairs of different reactors at different time points. The Bray-Curtis similarity index was chosen due to its effectiveness in handling ecological relative abundance data and its ability to account for differences in both species presence and abundance.

The datasets used for Bray-Curtis similarity calculation consisted of microbiome taxonomic relative abundance data obtained from reactor samples (Eq. S7). Each sample comprised a list of species along with their corresponding relative abundance values. Prior to analysis, all data were standardised to ensure comparability across different samples. This standardisation involved normalising the species abundance values to relative abundance to account for differences in sampling effort and total counts.

Where $BC_{ij}$ denotes the Bray-Curtis similarity between two samples $i$ and $j$, $x_{ik}$ denotes the abundance of species $k$ in sample $i$, $x_{jk}$ denotes the abundance of species $k$ in sample $j$, and $S$ denotes the number of species across both samples.

$$BC_{ij} = 1 - \frac{\sum_{k=1}^{S} |x_{ik} - x_{jk}|}{\sum_{k=1}^{S} (x_{ik} + x_{jk})} \quad \text{(S7)}$$

The standardised abundance data for each sample were organised into a matrix, where rows corresponded to samples and columns corresponded to species. The Bray-Curtis similarity index was then calculated for all pairs of samples using the pdist function from the scipy.spatial library [234]. The resulting pairwise Bray-Curtis dissimilarity values were converted to similarity values (Eq S7), yielding a similarity matrix with values ranging from 0 (identical communities) to 1 (completely dissimilar communities).

The Bray-Curtis dissimilarity index was implemented to evaluate the dissimilarity in microbial community composition across reactors (Eq. S7). Taxonomic dissimilarity was investigated using principal coordinate analysis (PCoA) and distance-based redundancy analysis (db-RDA), a multivariate statical method used to explore the influence of different variables on the microbial community. The db-RDA was conducted using the Bray-Curtis dissimilarity matrix to investigate the influence of influent and bioreactor chemical variables on microbial community composition. The db-RDA is a constrained ordination method that examines relationships between community dissimilarity and potentially explanatory variables. The db-RDA was performed using the skbio and statsmodels libraries in Python. The dissimilarity matrix served as the response variable, while the influent and bioreactor chemical variables were used as predictors. The results of the db-RDA were visualised in a biplot, where the principal coordinates (axes) represent the variation in microbial community composition explained by the influent and bioreactor chemical variables. The length and direction of the vectors indicate the strength and direction of the relationships between the influent and bioreactor chemical variables and the community composition.

Firstly, PCoA was applied to the Bray-Curtis dissimilarity matrix to reduce the data dimensionality, and db-RDA performs a form of linear regression on the results of the PCoA, treating the variables as predictors to find linear combinations of variables that best explain the variation in the diversity data. The variables investigated in the dissimilarity analysis included:

OLR, temperature, influent COD, pH, acetic acid, propionic acid, isobutyric acid, butyric acid, isovaleric acid, valeric acid, total VFA, melibiose, maltitol, glucose, mannitol, arabitol, and total sugar and sugar alcohol content.

A permutation test was conducted to assess the significance of the influence of measured variables on microbial community composition. The db-RDA was performed repeatedly with randomised data to generate a distribution of test statistics under the null hypothesis of no relationship between community composition and influent and bioreactor chemical variables. The observed test statistics were compared to the distribution obtained from the permutations to determine the significance of the relationships. A non-parametric permutational ANOVA (PERMANOVA) was implemented to test the null hypothesis that centroids and group dispersion were not defined by various influent and bioreactor chemical parameters tested. Statistical significance was tested at the 95% confidence interval ($p<0.05$).

#### 2.9. Machine learning models

This section outlines the methodology used to predict reactor performance using three machine learning algorithms: Lasso Regression, Random Forest Regression, and Bagging Regression. The selection of the best performing model was based on the coefficient of determination ($R^2$) score, and the feature importance was evaluated using SHAP values. The data was split into training and testing sets using 5-fold cross-validation to ensure robustness in model evaluation.

##### 2.9.1. Data Preprocessing and Cross-Validation

The features included in the dataset represent various parameters of the reactor that are expected to influence its performance. For model evaluation, 5-fold cross-validation was employed. In k-fold cross-validation, the dataset is randomly split into k subsets, and random state was set to 42 for reproducibility. In each iteration, the model is trained on k-1 folds and evaluated on the remaining fold. This process is repeated k times, each time with a different fold serving as the validation set. The performance metric used to evaluate the models was the $R^2$ score, which indicates how well the model explains the variance in the target variable, where higher values (closer to 1) indicate better predictive performance.

##### 2.9.2. Lasso Regression

Lasso Regression is a type of linear regression that incorporates L1 regularisation, which penalises the absolute magnitude of the model coefficients. This penalty encourages sparsity in the model, driving some of the coefficients to exactly zero. Lasso can automatically exclude irrelevant or redundant features by shrinking their coefficients to zero. Lasso regression was selected because of its ability to handle high-dimensional data where many features may be correlated with each other, making it prone to overfitting. Additionally, it offers built-in feature selection. The objective of Lasso is to minimise the sum of the residual sum of squares (RSS) with an added penalty proportional to the absolute value of the coefficients. Mathematically, the objective function is presented in Supplementary Equation S8.

Where $N$ denotes the number of observations, $p$ denotes the number of features, $y_i$ denotes the actual target value for the $i$ -th sample, $x_{ij}$ denotes the value of the $j$-th feature for the $i$-th sample, $\beta_0$ denotes the intercept term, $\beta = (\beta_1, \beta_2, \beta_3, \ldots, \beta_p)$ are the regression coefficients, $\lambda$ denotes the regularisation parameter controlling the penalty strength.

$$\min_{\beta} \frac{1}{2N} \sum_{i=1}^{N} (y_i - \beta_0 - \sum_{j=1}^{p} x_{ij}\beta_j)^2 + \lambda \sum_{j=1}^{p} |\beta_j| \quad \text{(S8)}$$

The L1 norm penalty, $\sum_{j=1}^{p} |\beta_j|$, forces some coefficients $\beta_j$ to shrink to zero, effectively excluding those features from the model. For Lasso Regression, the regularisation parameter alpha was set to 0.1. This is a commonly used value, as it provides a good balance between bias and variance. Larger values of alpha increase regularisation, which can lead to underfitting, while smaller values may lead to overfitting.

##### 2.9.3. Random Forest Regression

Random Forest Regression is an ensemble learning method based on decision trees, where multiple decision trees are trained on random subsets of the data and the final prediction is averaged across all trees. This approach helps reduce the variance compared to individual decision trees and increases the robustness of the model. Random Forest was selected due to its ability to model non-linear relationships between the features and target variable, and its robustness against overfitting. Since reactor performance is likely influenced by complex, non-linear interactions between multiple operational parameters, Random Forest is well-suited to capture these relationships. The random forest algorithm is based on an ensemble of decision trees, which work together to make predictions. Each decision tree in a random forest is trained on a subset of the training data. For each tree $T$ in the random forest, it takes a random sample $X_T$ from the dataset $X$. After that, it selects a random subset of features $F_T$ from the feature space to find the best split based on certain criteria. For regression, the criterion is typically Mean Squared Error (MSE) (Supplementary Equation 9).

Where $N$ denotes the number of samples in the node, $y_i$ denotes the actual value, and $y$ denotes the predicted mean value of the node.

$$MSE = \frac{1}{N}\sum_{i=1}^{N}(y_i - y)^2 \qquad \text{(S9)}$$

Once each tree in the random forest is trained, they are combined to make a prediction. The final prediction is the average of all tree predictions (Supplementary Equation 10).

Where $M$ denotes the total number of trees and $y_{T_m}$ denotes the prediction from each individual tree.

$$y = \frac{1}{M}\sum_{m=1}^{M} y_{T_m} \quad \text{(S10)}$$

The random forest approach leverages this ensemble of trees to improve model accuracy and reduce overfitting, which individual decision trees are prone to. The model was trained using the default settings. These include Number of trees = 100, Maximum depth of trees = None (trees are grown until all leaves are pure), Minimum number of samples required to split a node = 2, Minimum number of samples required at a leaf node = 1, Criterion used to evaluate splits = Mean Squared Error (MSE), Random state was set to 42 for reproducibility.

###### 2.9.4. Bagging Regression

Bagging is an ensemble method that combines multiple instances of the same model, each trained on a random bootstrap sample from the training data. The final prediction is obtained by averaging the predictions of the individual models. Bagging helps reduce variance and improves model stability. Bagging Regression was selected as it is an effective method for improving the performance of base regression models that may be prone to overfitting, such as decision trees. By averaging over many different models trained on different subsets of the data, bagging helps increase prediction accuracy and model robustness. Bagging generates multiple datasets $D^{(b)}$ by bootstrapping the original dataset $D$. Each dataset $D^{(b)}$ is formed by sampling $N$ observations with replacement from $D$, where $N$ is the number of original samples. Each base learner $f^{(b)}(X)$ is trained on its corresponding bootstrapped dataset $D^{(b)}$. The final prediction for bagging regression is obtained by averaging the predictions of all the base learners. For a given input $X$, the prediction $y$ is defined in Supplementary Equation 11.

Where $B$ denotes the total number of bootstrapped datasets (and corresponding base learners) and $f^{(b)}(X)$ denotes the prediction from the $b$-th base learner.

$$y = \frac{1}{B}\sum_{b=1}^{B} f^{(b)}(X) \qquad \text{(S11)}$$

##### 2.9.5. Variance reduction

The averaging of predictions reduces the variance of the model without increasing bias significantly, leading to more robust predictions. The model was trained using the default settings. These include Number of base estimators = 10, Maximum number of samples to train each base estimator = 1.0, Maximum number of features used by each base estimator = 1.0, Bootstrap sampling (`bootstrap`) = True, Random state was set to 42 for reproducibility.

##### 2.9.6. Model Evaluation: $R^2$ Score

To evaluate the performance of the models, the $R^2$ score was calculated for each fold in the cross-validation. The $R^2$ score is defined by Supplementary Equation S12. Where: $y_{true}$ are the true target values, $y_{pred}$ are the predicted values, $\overline{y_{pred}}$ is the mean of the true target values. An $R^2$ value closer to 1 indicates that the model explains a large portion of the variance in the target variable, suggesting better performance. The algorithm with the highest mean $R^2$ scores across the 5-fold cross-validation was selected as the final model.

$$R^2 = 1 - \frac{\sum(y_{true} - y_{pred})^2}{\sum(y_{true} - \overline{y_{pred}})^2} \qquad \text{(S12)}$$

##### 2.9.7. Feature Importance Evaluation using SHAP Values

To understand the influence of individual features on the model's predictions, SHAP values were used. SHAP values are based on cooperative game theory and provide a consistent method for explaining the contribution of each feature to a given prediction. They decompose the prediction of a model into contributions from each feature, which allows for a clear understanding of how each input affects the output. SHAP values were used to identify which features had the most significant impact on reactor performance predictions. Since the models used in this study, particularly Random Forest and Bagging, can be seen as "black box" models, SHAP values provide transparency by assigning a numerical importance to each feature, facilitating better interpretability. The SHAP values were computed for the best-performing model (as determined by the $R^2$ score) to visualise and rank feature importances. The SHAP summary plots were used to provide an overall understanding of the relationships between features and reactor performance.

###### 2.9.8. Algorithm Selection

The final model selection was based on the mean $R^2$ score obtained from the 5-fold cross-validation. The algorithm that produced the highest mean $R^2$ score was selected as the best model for predicting reactor performance. Once the best model was identified, SHAP values were used to interpret the model's decisions and assess which features contributed most to the model's predictions. Machine learning models are open source and available at https://github.com/MGuo-Lab/AnaerobicDigestionML.

##### 2.10. Nucleotide sequence accession numbers.

All sequencing data have been submitted to the European Nucleotide Archive (ENA) under the project ID PRJEB80086 and under accession numbers ERS22193437-ERS22335023. The correspondence between accession numbers, sequences and metadata is provided in Supplementary Database 1.

### 3. Supplementary Results

#### 3.1. Operational parameter monitoring

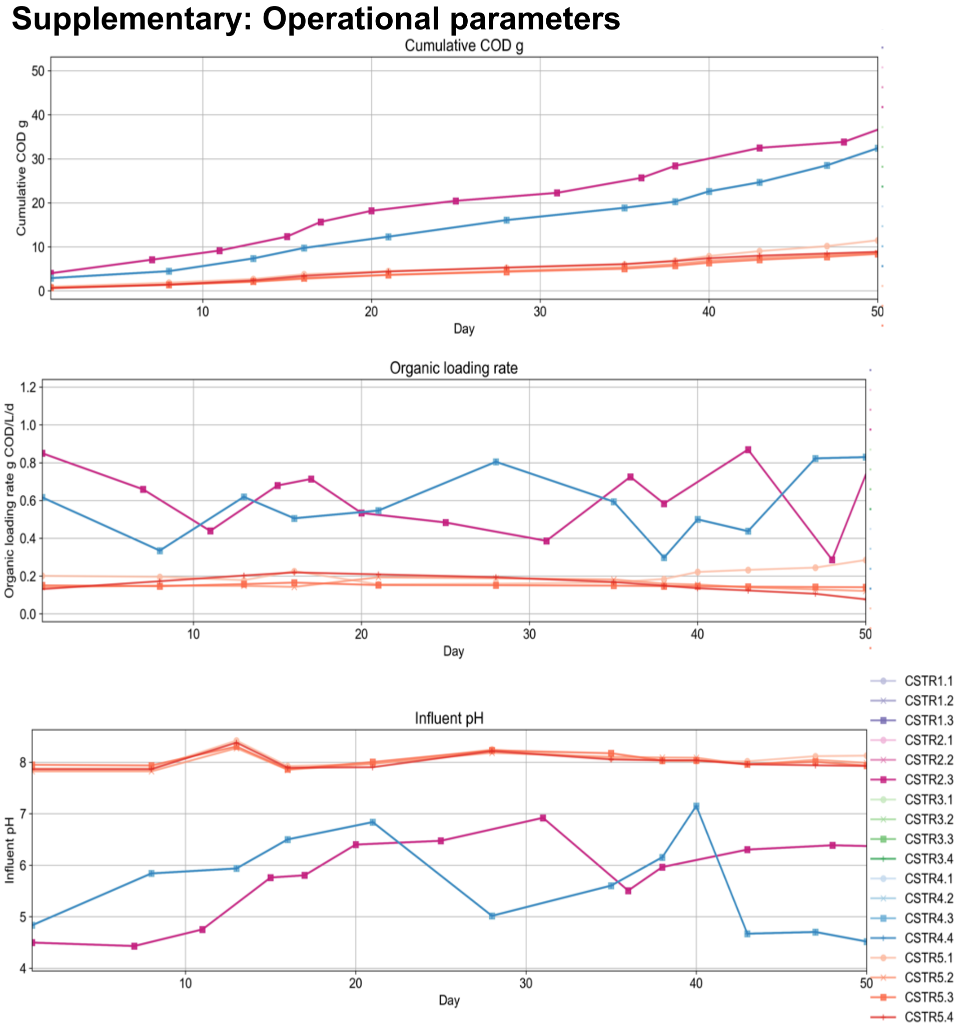

**Supplementary Figure 3.1.** Operational parameter monitoring of experimental runs including cumulative COD loading (g), organic loading rate (g COD/d/L) and influent pH.

#### 3.2. Core microbiome construction per group

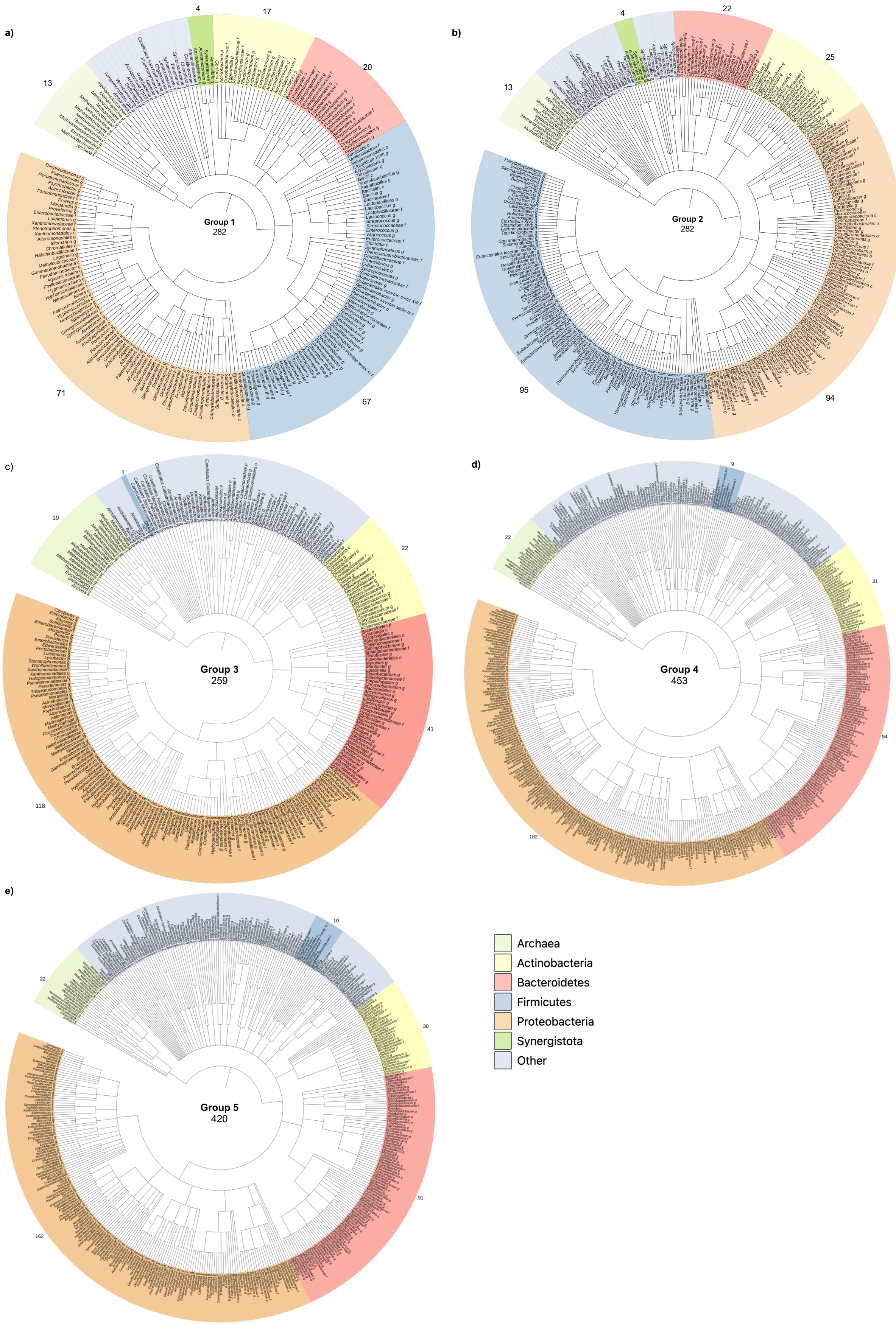

**Figure 3.2.** Core microbiome of genera present across all reactors within **(a)** mesophilic single-stage reactors (run 1), **(b)** thermophilic single-stage reactors (run 1), **(c)** single-stage reactors (run 2), **(d)** first-stage reactors (run 2), and **(e)** second-stage reactors (run 2). Archaea are highlighted as green, *Firmicutes* are highlighted as blue, *Actinobacteria* are highlighted as yellow, *Bacteroidetes* are highlighted as red, *Proteobacteria* are highlighted as orange and other bacterial phyla are highlighted as purple. Phylogenetic trees are constructed using iTOL [235].

#### 3.3. Machine learning microbial community prediction

Predictive machine learning models were developed to estimate the relative abundance of microbial phyla and reactor performance. Reactor performance was measured in terms of COD removal, cumulative biogas and specific daily biogas. The input factors included influent COD, temperature, pH, organic loading rate, sugars, sugar alcohols, amino acids, and volatile fatty acids. Three machine learning models were evaluated: Linear Regression, Random Forest, and Bagging Regression.

Exploratory results are displayed for microbiome prediction using predictive machine learning models (Supplementary Figure 3.3-3.5). The findings of this work indicate that temperature, pH, and organic loading rate are the most influential factors in predicting the relative abundance of phyla and reactor performance. The use of machine learning models, particularly Random Forest, has proven to be highly effective in this work for reactor performance prediction. However, further prediction of microbiome composition will require additional data. We recommend an extended run period of at least 6 months, sampling twice per week to generate enough data for future exploration of microbiome prediction based on operational parameters and feedstock composition.

### Output: Phylum or genus level relative abundance

#### a) Model inputs and outputs

| INPUTS | OUTPUTS |
|---|---|
| Organic Loading Rate | Relative abundance of phyla |
| Temperature | |
| pH | |
| Day | |
| Influent COD | |

#### b) Model performance metrics.

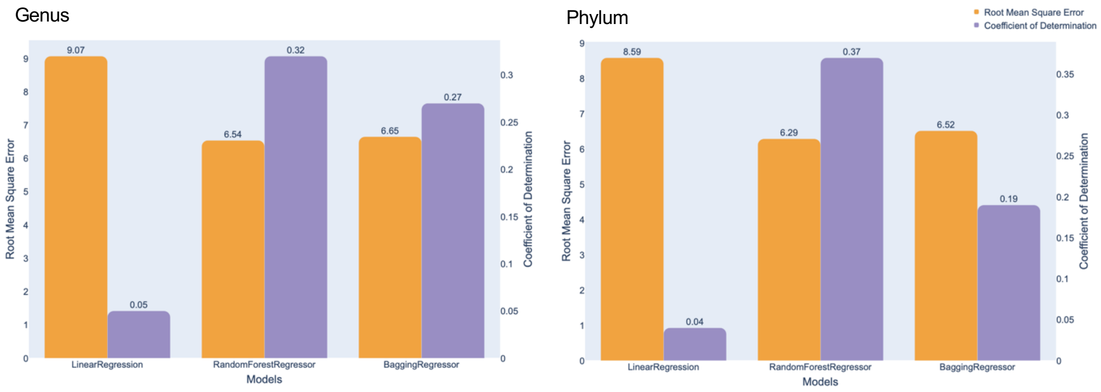

#### c) Top factors for genus and phylum relative abundance prediction

Top factors (genus)
- Temperature
- Day
- pH
- Influent COD
- OLR
- *Clostridiaceae*
- *Sulfurimonas*
- *Alarcobacter*

Top factors (phylum)
- Influent COD
- Day
- OLR
- Temperature
- pH
- *Firmicutes*
- *Proteobacteria*
- *Bacteroidetes*

#### d) Influence of each parameter on the top 5 most influential genera.

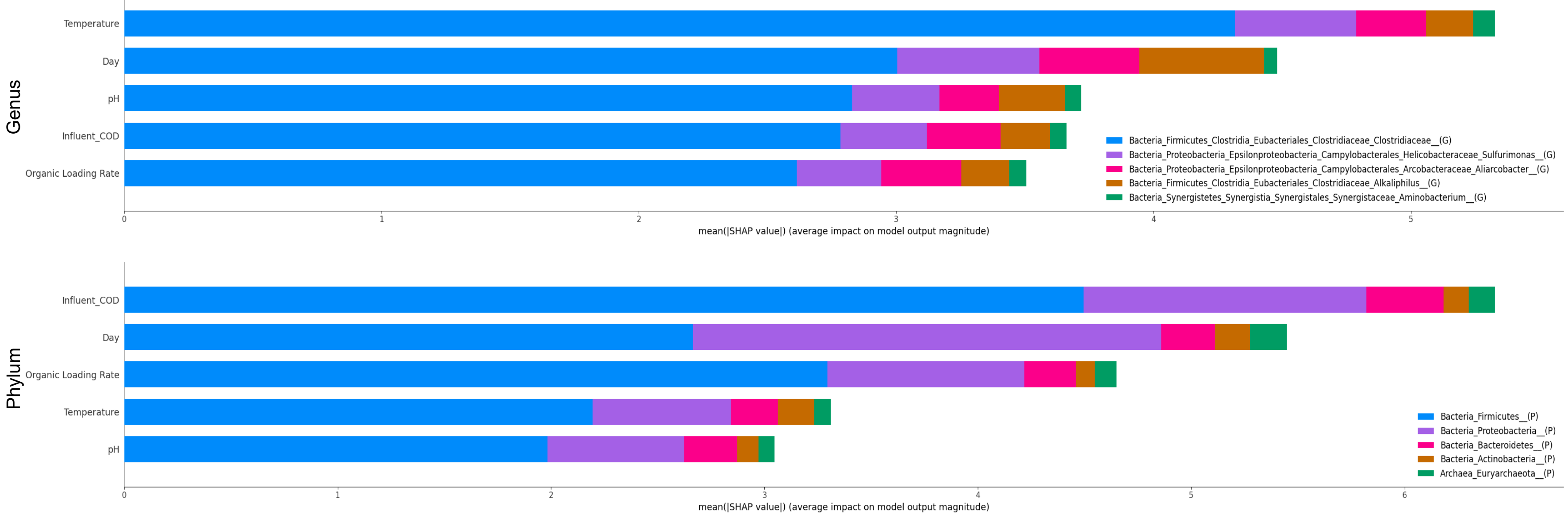

**Supplementary Figure 3.3.** Predictive machine learning models for estimation of relative abundance of the AD microbiome at the genus and phylum taxonomic levels. **(a)** Model inputs and outputs. **(b)** Model performance metrics. **(c)** Top factors influencing the model. **(d)** Influence of each parameter on the top five genera/phyla measured by SHAP values.

**Output: Phylum relative abundance including detailed feedstock composition**

**a) Model inputs and outputs**

| INPUTS | OUTPUTS |
|---|---|
| Organic Loading Rate | Relative abundance of phyla |
| Temperature | |
| pH | |
| Sugar | |
| Sugar Alcohol | |
| Amino Acid | |
| Volatile Fatty Acid | |
| Influent COD | |
| Day | |

**b) Model performance metrics**

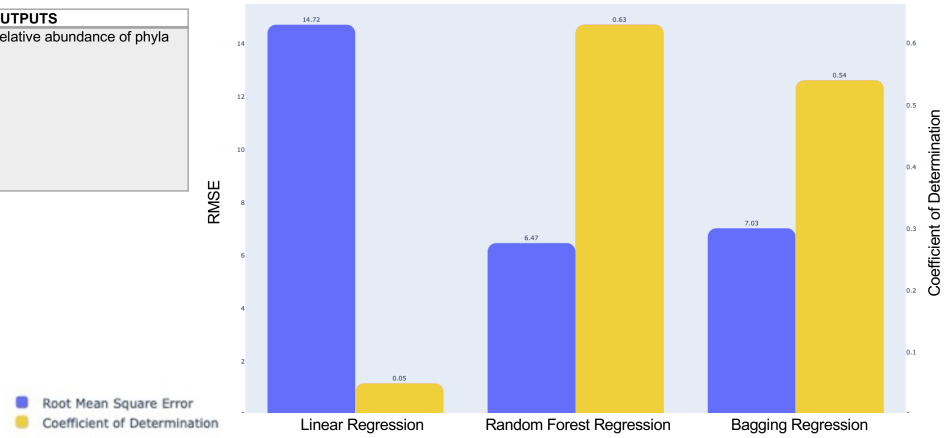

**c) SHAP values of input parameters on top five influential phyla**

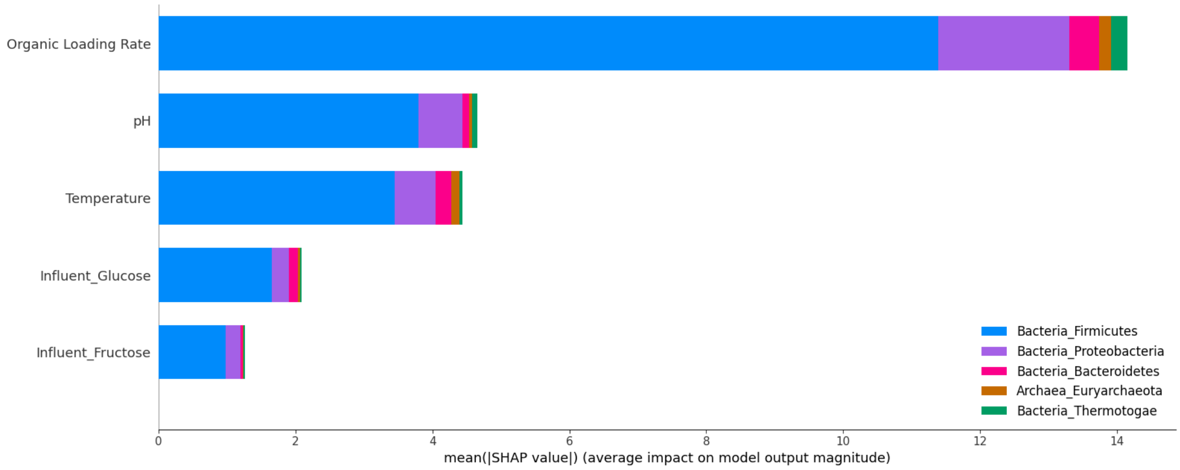

**Supplementary Figure 3.4.** Predictive machine learning models for estimation of phylum relative abundance including detailed feedstock chemical composition. **(a)** Model inputs and outputs. **(b)** Model performance metrics. **(c)** Impact of the top five most influential factors on the top phyla measured by SHAP values.

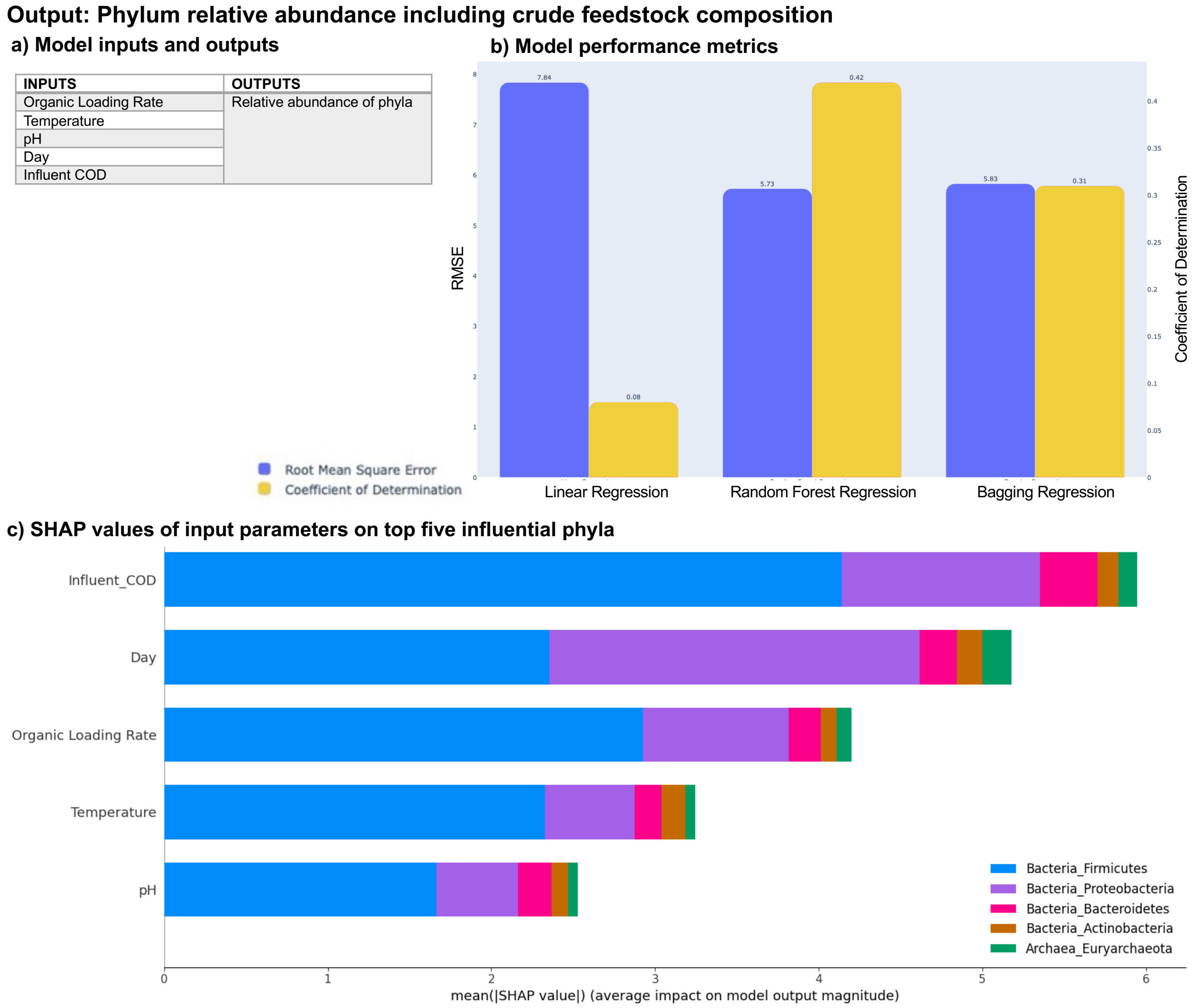

**Supplementary Figure 3.5.** Predictive machine learning models for estimation of phylum relative abundance including crude feedstock characterisation. **(a)** Model inputs and outputs. **(b)** Model performance metrics. **(c)** Impact of the top five most influential factors on the top phyla measured by SHAP values.

#### 3.4. Mycoprotein fermentation wastewater additional detailed chemical composition

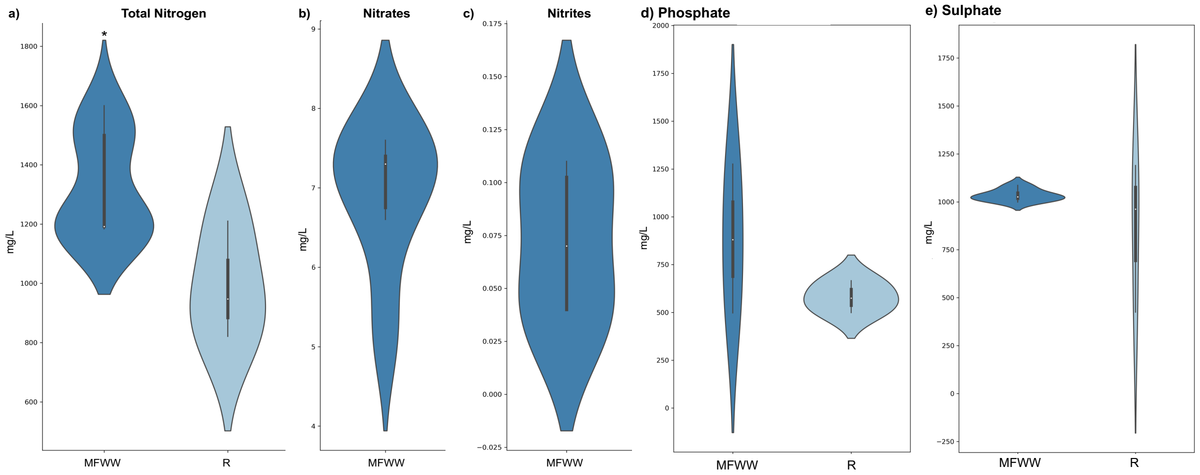

**Supplementary Figure 3.6.** Mycoprotein fermentation wastewater additional detailed chemical composition. **(a)** Total nitrogen, mg/L. **(b)** Nitrates, mg/L. **(c)** Nitrites, mg/L. **(d)** Phosphate, mg/L. **(e)** Sulphate, mg/L.

### 4. Supplementary database 1

Database containing all data relating to:

##### 4.1. Literature Study Details

Relative abundance (%) of species reported to be involved in AD and experimental conditions including: Study reference, feedstock source, feedstock type, inoculum source, inoculum source reactor type, inoculum source location, experimental reactor scale, experimental reactor type, experimental reactor total volume (L). From 197 papers

##### 4.2. Literature Growth Conditions

Contains all information and notes from literature review, including: microorganism name, classification, NCBI taxonomy number, pH range (optimum), temperature preference, substrate, notes, and reference. ** indicates genera has been reported in literature, but not that particular species

##### 4.3. Experimental factors

Experimental factors measured within anaerobic digestion experimental studies for mesophilic (CSTR1.1 to CSTR1.3), thermophilic (CSTR2.1 to CSTR2.3), single-stage (CSTR3.1 to CSTR3.4), first-stage (CSTR4.1 to CSTR4.4), second-stage (CSTR5.1 to CSTR5.4), including: Biogas (L/g COD/d), cumulative biogas (L), specific daily biogas (L/g COD/d), COD removal (%), Organic loading rate (g COD/L/d), Temperature (C), influent COD (g/L), influent pH, Influent volatile fatty acid content (acetic, propionic, isobutyric, butyric, isovaleric, valeric acid, total VFA, g/L), Influent sugar and sugar alcohol (melibiose, maltitol, glucose, mannitol, arabitol, total sugar, g/L), reactor pH, reactor volatile fatty acid, reactor sugar and sugar alcohol

##### 4.4. Taxonomy

Relative abundance of taxonomic classification of sequences obtained from mesophilic (CSTR1.1 to CSTR1.3), thermophilic (CSTR2.1 to CSTR2.3) single-stage (CSTR3.1 to CSTR3.4), first-stage (CSTR4.1 to CSTR4.4), second-stage (CSTR5.1 to CSTR5.4). Taxa are reported to the lowest common ancestor and details are given of taxonomy from the kingdom, phylum, class, order, family and genus level.

##### 4.5. Nucleotide sequence metadata

The correspondence between sequencing sample ID, taxonomy and metadata.

##### 4.6. ENA Accession Numbers

The correspondence between sample ID, accession numbers and taxonomic classification.

##### 4.7. Bibliography

References

224 Rivera, B. Fast Analysis of Sucrose, Glucose, and Fructose Composition in Fruit Juices and Processed Beverages using Simplified HPLC Methodology.

225 Bio-Rad. (ed Bio-Rad) 17-24 (Richmond, CA, USA, 1997).

226 Schloss, P. D. *et al.* Introducing mothur: open-source, platform-independent, community-supported software for describing and comparing microbial communities. *Applied and environmental microbiology* **75**, 7537-7541 (2009).

227 Quast, C. *et al.* The SILVA ribosomal RNA gene database project: improved data processing and web-based tools. *Nucleic acids research* **41**, D590-D596 (2012).

228 Ma, G., Chen, Y. & Ndegwa, P. Association between methane yield and microbiota abundance in the anaerobic digestion process: A meta-regression. *Renewable and Sustainable Energy Reviews* **135**, 110212 (2021).

229 Pasalari, H., Gholami, M., Rezaee, A., Esrafili, A. & Farzadkia, M. Perspectives on microbial community in anaerobic digestion with emphasis on environmental parameters: a systematic review. *Chemosphere* **270**, 128618 (2021).

230 Schloss, P. D. Reintroducing mothur: 10 years later. *Applied and environmental microbiology* **86**, e02343-02319 (2020).

231 Shannon, P. *et al.* Cytoscape: a software environment for integrated models of biomolecular interaction networks. *Genome research* **13**, 2498-2504 (2003).

232 Deng, Y. *et al.* Molecular ecological network analyses. *BMC bioinformatics* **13**, 1-20 (2012).

233 Chao, A. & Chiu, C.-H. Species richness: estimation and comparison. *Wiley StatsRef: statistics reference online* **1**, 26 (2016).

234 Feranchuk, S. *et al.* Tools and a web server for data analysis and presentation in microbial ecology. *Community Ecology* **20**, 230-237 (2019).

235 Letunic, I. & Bork, P. Interactive Tree Of Life (iTOL) v5: an online tool for phylogenetic tree display and annotation. *Nucleic acids research* **49**, W293-W296 (2021).
